# Th1, Th2 and Th17 inflammatory pathways synergistically correlate with cardiometabolic processes. A case study in COVID-19

**DOI:** 10.1101/2021.11.29.470414

**Authors:** James R. Michels, Mohammad Shaheed Nazrul, Sudeep Adhikari, Dawn Wilkins, Ana B. Pavel

## Abstract

A predominant source of complication in SARS-CoV-2 patients arises from the *cytokine storm*, an elevated expression of inflammatory helper T-cell associated cytokines that can lead to tissue damage and organ failure. The high inflammatory burden of this viral infection often results in cardiovascular comorbidities. A better understanding of the interaction between the *cytokine storm* and cardiovascular proteins might inform medical decisions and therapeutic approaches. We hypothesized that all major helper T-cell inflammatory pathways (Th1, Th2 and Th17) synergistically contribute to cardiometabolic modifications in serum of COVID-19 patients. We proved our hypothesis by integrating Th1, Th2 and Th17 cytokines to predict expression of cardiometabolic proteins profiled by OLINK proteomics.

## 1 Introduction

The respiratory virus SARS-CoV-2 has quickly spread around the world, resulting in a total of over 200 million reported cases and over 5 million reported deceased to date. ^1^ Despite incredible advancements in the development of vaccines aiding the prevention of cases, targeted treatments toward COVID-19 symptoms and effects are still needed. Complications such as the acute respiratory distress syndrome (ARDS), respiratory failure, hepatic and renal insufficiency are common in more severe cases. ^2^ Proteomic profiles are currently being investigated in order to study the effect of these cytokines in COVID-19 disease. ^3^ A better understanding of cytokines interactions with the cardiovascular system, might enhance development of novel immunomodulatory therapies and reduce mortality in COVID-19 vulnerable patients.

Previous studies have shown various helper T-cell responses in SARS-CoV-2 infections. One study in elderly SARS-CoV-2 patients found that the predominant response to infection was in the form of Th1-associated cytokines. However, the levels of Th17-associated cytokines were also detected in COVID-19 severe disease. ^4–6^, and Th2 secretions were associated to SARS-CoV-2 susceptibility. ^7,8^

We have recently shown an association between a shift in Th1 towards Th2 expression in patients with increased risk of severe COVID-19 disease. ^8^ Our findings were further validated by a clinical trial conducted in a cohort of atopic patients, which demonstrated a significantly higher rate of asymptomatic cases among patients treated with Dupilumab, a Th2-specific inhibitor, as compared to other systemic treatments. ^9^ Furthermore, Th2 cytokines have been previously linked to vascular inflammation risk in patients with Type 2 inflammatory conditions, such as atopic dermatitis. ^10,11^

In this study we used serum protein profiles from Massachusetts General Hospital COVID-19 registry. ^12^ Immune mediators were previously associated with SARS-Cov-2 infection in this data set, ^12^ however no prior study has integrated all major helper T-cell pathways (Th1, Th2 and Th17) to evaluate the overall impact of COVID-19 *cytokine storm* on cardiometabolic proteins.

We hypothesized that all Th1, Th2, and Th17 immune pathways synergistically impact cardiometabolic processes associated with COVID-19 severe disease. While increased expression in Th1 and Th17 cytokines has been associated with other viral infections such as, influenza ^13^, rhinoviruses ^14^, and other coronaviruses like MERS ^15^, these infections do not show severe cardiovascular manifestations such as those observed in COVID-19 disease. In addition, COVID-19 also has a higher mortality rate (estimated to be between 3% to 4%) compared to influenza, rhinoviruses, and drastically more deaths than the total deaths from MERS. ^16^ Thus, we hypothesized that Th2 cytokines are key players in the immune response triggered by SARS-CoV-2 and that Th1, Th2 and Th17 pathways synergistically contribute to the complex cytokine network that severely affects cardiometabolic processes.

## 2 Materials and Methods

### 2.1 Data

In this study we used the publicly available Massachusetts General Hospital COVID-19 registry ^12^ consisting of 383 patients (after removal of one patient with missing data), where 305 were COVID-19 positive and 78 were COVID-19 negative with other respiratory infections. Patients in this data set were classified by their severity as 42 COVID-19 positive and 7 COVID-19 negative deceased patients (*who max score* = 1), 67 COVID-19 positive and 16 COVID-19 negative intubated patients (*who max score* = 2) and 196 COVID-19 positive and 55 COVID-19 negative non-severe patients (*who max score* ≥ 3).

We included in our analysis serum proteins profiled by OLINK Explore platform, ^12^ such the *Cardiometabolic* panel consisting of 355 detected proteins, and all available Th1, Th2 and Th17 cytokines as previously described. ^8^ OLINK proteomics platform has been extensively used to profile targeted biomarkers and drug targets associated with various diseases and tissue types. ^8,17^

The helper T-cell pathways were defined by 11 Th1 markers (CCL3, CCL4, CXCL11, CXCL10, CXCL9, IL2RA, IFNG, IFNGR1, IFNGR2, IL12B and IL1B), 14 Th2 markers (CCL11, CCL13, CCL17, CCL22, CCL24, CCL26, CCL7, IL10, IL13, IL33, IL4R, IL5, IL7R and TSLP) and 13 Th17 markers (CCL20, S100P, IL6, IL6R, LCN2, S100A12, CXCL1, PI3, IL17A, IL17F, CXCL3, IL12A and IL12B). ^8^

### 2.2 Statistical Modeling

We first applied a linear regression model to associate the normalized expression (NPX) for each immune or cardiometabolic protein (dependent variable) with disease severity (independent variable) in both COVID-19 positive and negative patients. We then displayed heatmaps of the mean estimates of deceased, intubated and non-severe groups for all Th1, Th2 and Th17 cytokines, and all significantly differentially expressed cardiometabolic markers between any comparison (deceased versus non-severe, intubated versus non-severe or deceased versus intubated) by *False Discovery Rate* (*FDR*) < 0.05.

Next, we integrated all Th1, Th2 and Th17 immune mediators by elastic net regularized generalized linear models with 10-fold cross-validation, using *cv*.*glmnet* function from *glmnet* R package, to predict expression levels of each cardiometabolic protein on the OLINK panel based on helper T-cell immune profiles.

We then ranked the best predictions among all 355 cardiometabolic markers by Pearson correlation coefficient calculated between the real and predicted value, and considered as best fit those correlations with *r* ≥ 0.7 and *FDR* < 0.05.

To visualize the interaction between immune and cardiometabolic markers with *r* ≥ 0.7 we used *igraph* R package. The edges of the network represent non-zero coefficients of the elastic net regression model with absolute value ≥ 0.05. We used this threshold to exclude weak interactions and filter noise.

We performed all statistical analyses using R programming language.

## 3 Results

We first analyzed all cytokines of T-helper cell types pathways (Th1, Th2 and Th17) in both COVID-19 positive and COVID-19 negative patients and displayed them as heatmaps of mean expression estimates stratified by disease severity (Figure 1). We observed an increasing trend in protein expression from non-severe to deceased COVID-19 patients in most of Th1 and Th17 mediators and in more than 50% of Th2 mediators. We found significant increases (*p − value* < 0.05) in Th1 (IFNGR1, CCL3, CCL4, CCL5, CXCL9, CXCL10, IL1B, IL2RA), Th2 (CCL11, CCL13, CCL7, CCL24, IL4R) and Th17 (S100A12, S100P, CCL20, LCN2, PI3, CXCL1, IL17A, IL6) mediators in deceased as compared to non-severe COVID-19 patients (Figure 1 A, Supplementary Table 1). In contrast, COVID-19 negative patients with other respiratory infections did not show significant increasing trends in deceased as compared to non-severe patients in the Th1, Th2 and Th17 pathways (Figure 1 B, Supplementary Table 2).

**Fig. 1.**
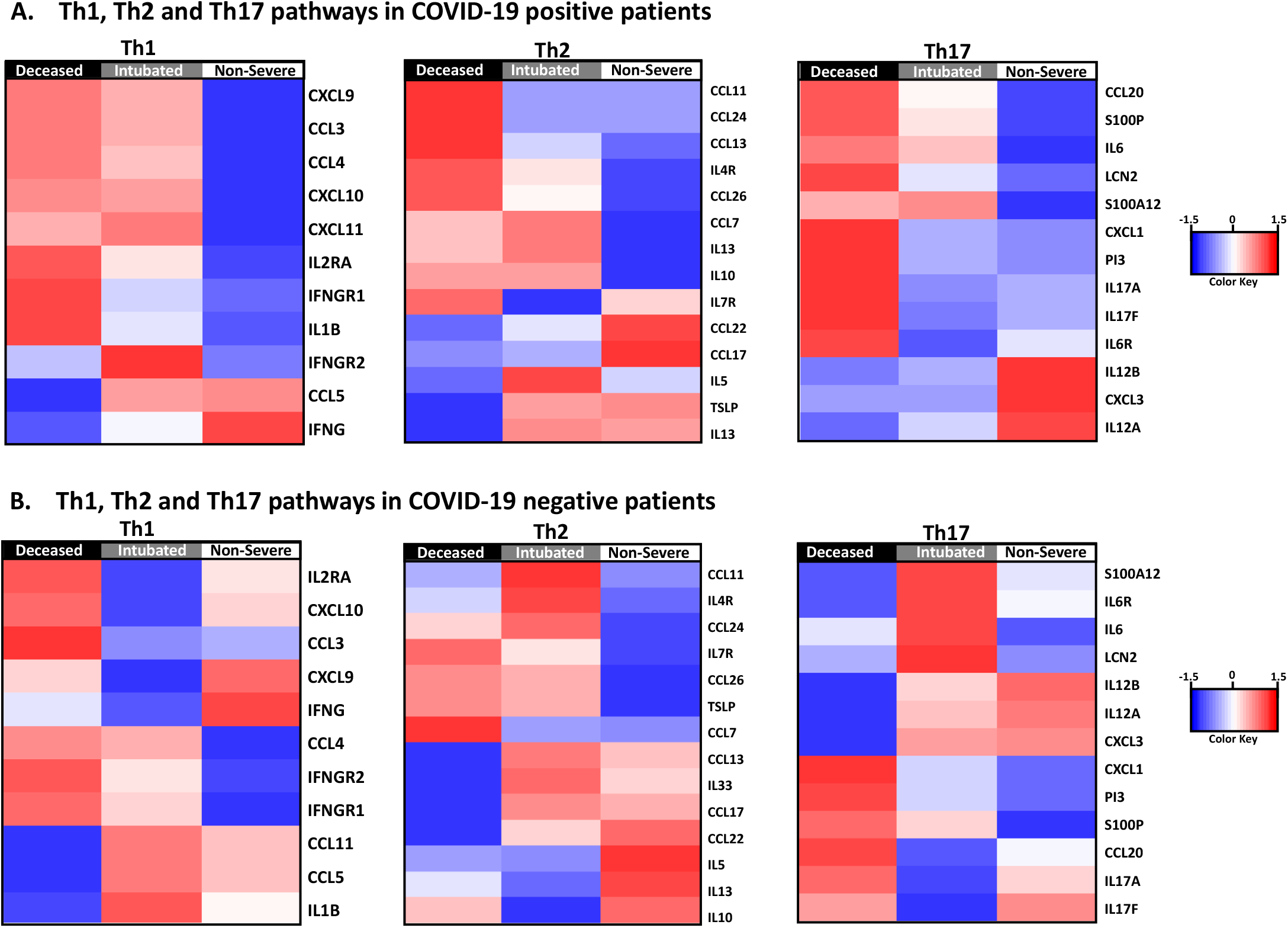
A. Heatmaps of estimated mean expression of Th1, Th2 and Th17 mediators stratified by severity in COVID-19 positive patients. B. Heatmaps of estimated mean expression of Th1, Th2 and Th17 mediators stratified by severity in COVID-19 negative patients. The heatmaps are z-score normalized with blue denoting decreased expression and red denoting increased expression.

Next, we evaluated 355 cardiometabolic proteins detected by OLINK Explore Cardiometabolic panel. We found that 35 of these proteins were strongly associated with COVID-19 severity (| *f old − change*| ≥ 2, *FDR* < 0.05) as shown in Figure 2 A (Supplementary Table 3). All these 35 differentially expressed proteins were significantly increased in deceased compared to non-severe COVID-19 patients, and several of them (i.e. LTBP2, RNASE3, CHI3L1, CSTB, RETN, GDF15, CXCL8, PLA2G2A, IL1RL1 and NADK) also showed significantly elevated expression in intubated compared to the non-severe patients.

**Fig. 2.**
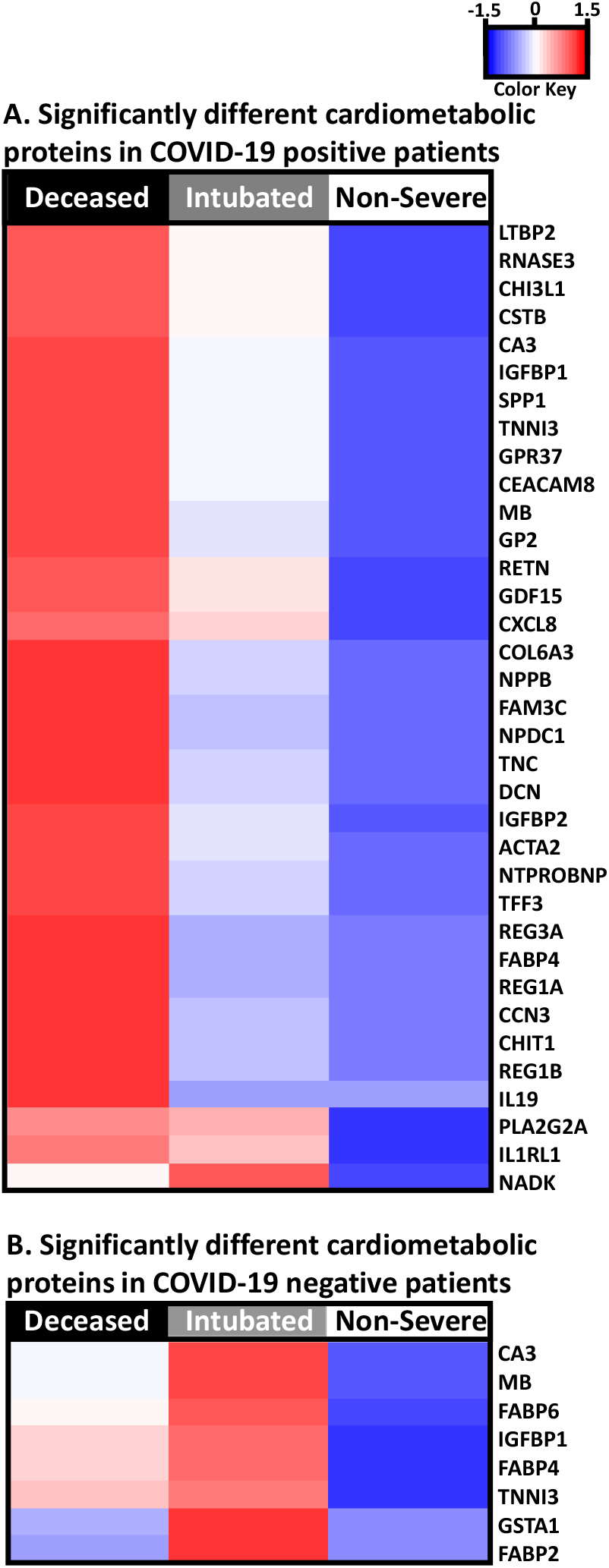
A. Heatmap of estimated mean expression of top significantly differentially expressed cardiometabolic proteins stratified by COVID-19 severity (35 proteins). Differentially expressed proteins were determined by | *f old − change*| ≥ 2 and *FDR* < 0.05 in any comparison (deceased versus non-severe, intubated versus non-severe or deceased versus intubated). All 35 proteins achieved statistical significance between deceased and non-severe groups. B. Heatmap of estimated mean expression of top significantly differentially expressed cardiometabolic proteins stratified by severity of COVID-19 negative patients with other respiratory conditions (8 proteins). Differentially expressed proteins were determined by | *f old − change*| ≥ 2 and *FDR* < 0.05 in any comparison (deceased versus non-severe, intubated versus non-severe or deceased versus intubated). None of these 8 proteins achieved significance between deceased versus non-severe patients. The heatmaps are z-score normalized with blue denoting decreased expression and red denoting increased expression.

In contrast to this strong cardiometabolic signal associated with COVID-19 severity, COVID-19 negative patients with other respiratory infections showed no statistically significant differences between deceased and non-severe patients, and only 8 proteins were increased in intubated as compared to non-severe patients (Figure 2 B, Supplementary Table 4).

Next, we sought to test our hypothesis that all Th1, Th2 and Th17 cytokines contribute to cardiometabolic modifications associated with SARS-CoV-2 infections. We integrated all Th1, Th2 and Th17 immune markers by elastic net regularized linear regression to predict the abundance of each cardiometabolic protein in COVID-19 positive patients. By this approach we were able to rank all the cytokines and immune mediators by their predictive potential.

We evaluated the best fit of our predictions by Pearson correlation coefficient computed between the measured and predicted expression values for each of the 355 cardiometabolic proteins (Supplementary Table 5). We identified 186 significant predictions (*r* ≥ 0.7 and *FDR* < 0.05), and further represented them as a graph structure, where the edges (links) represent the non-zero coefficients (*abs* ≥ 0.05) corresponding to each cytokine. We found that all Th1, Th2 and Th17 immune pathways synergistically describe the expression of these 186 cardiometabolic proteins. Most of the links in our networks were positive connections (Figure 3), suggesting that an increased production in cytokines stimulates the overall production of cardiometabolic proteins. Figure 3 A shows all positive links between Th1 (yellow-green), Th2 (blue) and Th17 (green) cytokines and the significantly predicted cardiometabolic proteins (gray), highlighting the extent of the immune system’s impact on cardiometabolic processes.

**Fig. 3.**
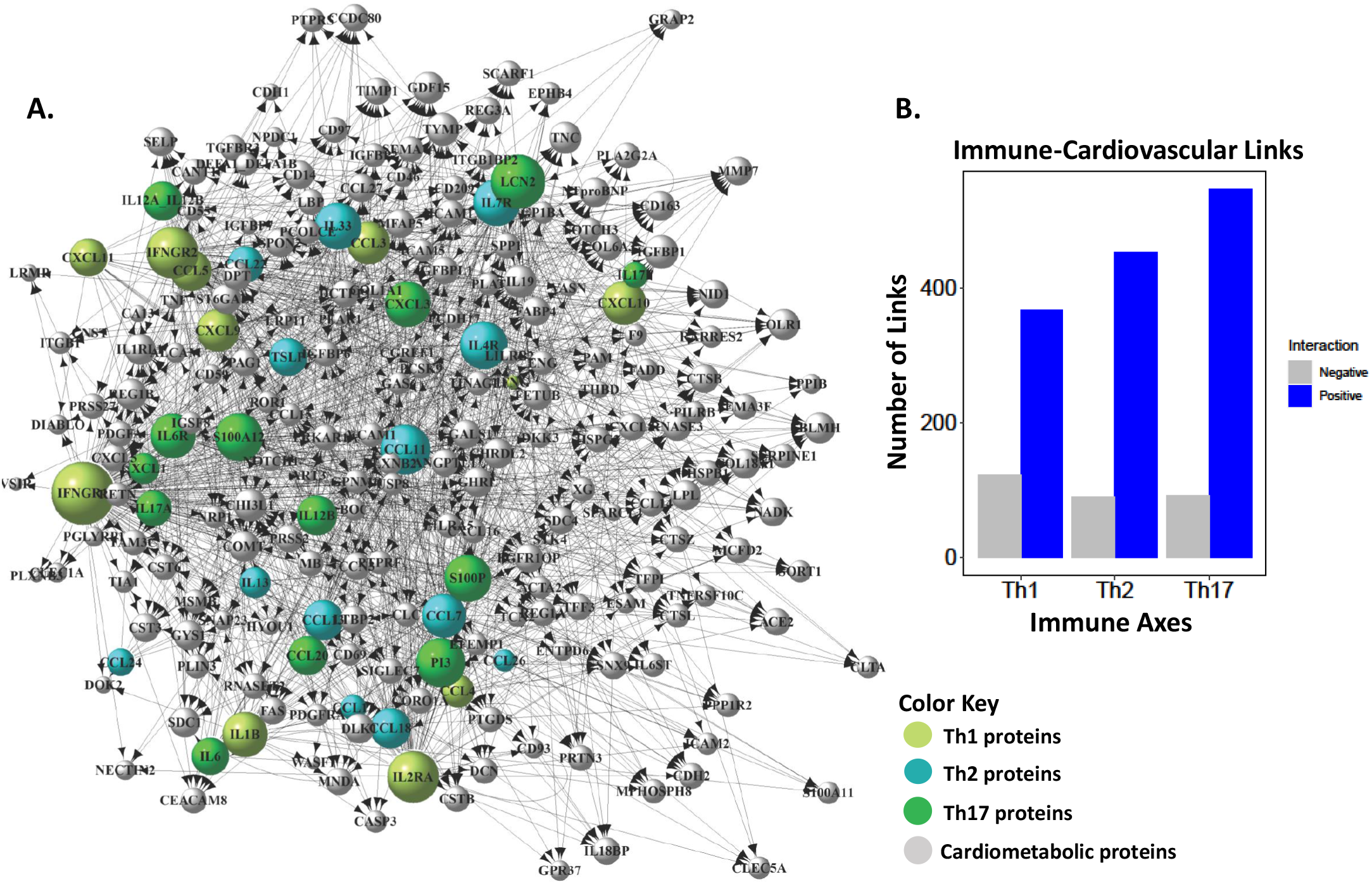
A. Positive associations (links) between the major components of the *cytokine storm*, Th1 (yellow-green), Th2 (blue) and Th17 (green) mediators, and the predicted cardiometabolic proteins (gray). B. Th1 pathway had 367 positive links and 122 negative links, Th2 had 452 positive links and 90 negative links, and Th17 had 546 positive links and 91 negative links with cardiometabolic proteins.

We found that 31 of the 35 cardiometabolic proteins associated with COVID-19 severity (Figure 2 A) were also significantly predicted by the *cytokine storm*, except for CA3, TNNI3, NPPB, CHIT1 which did not reach our prediction threshold for the “goodness of fit” (*r* ≥ 0.7 and *FDR* < 0.05). Most of the cytokines and immune mediators positively contributed to these predictions. In Figure 4 we highlighted 20 of these cardiometabolic proteins that were both significantly associated with COVID-19 severity and significantly predicted by the *cytokine storm*, and in addition were found to be associated with cardiovascular inflammation and hypertension in other studies. ^18–37^ Across these 20 cardiovascular markers, the most common association with Th1 pathway was represented by IFNGR1 (85%), while the most common Th2 associations included CCL11 (50%) and CCL7 (40%), and the most common Th17 associations included PI3 (50%), LCN2 (50%) and IL6 (45%).

**Fig. 4.**
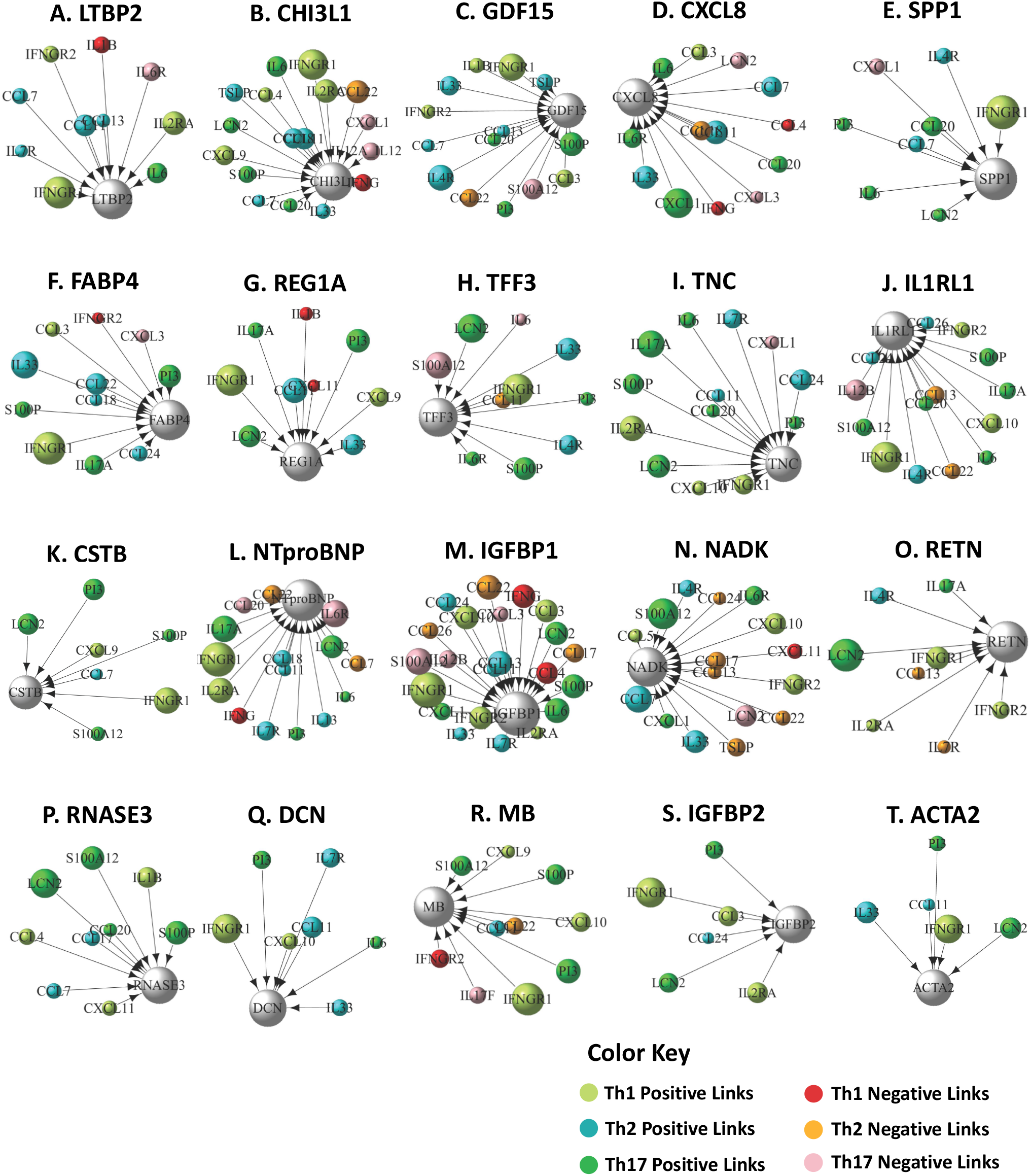
A-T. Immune-cardiometabolic associations for 20 significant predictions (*r* ≥ 0.7 and *FDR* < 0.05). These cardiometabolic predicted proteins were also significantly associated with COVID-19 severity in this data set, as well as previously associated with cardiovascular inflammation by other studies. The vertex color maps with the type of helper T-cell cytokines and immune mediators and the sign of correlation is explained in the color key. The circle size of the cytokines and immune mediators is proportional to their correlation strength with the respective cardiometabolic protein. Cardiometabolic proteins are represented in gray and their size is proportional to the number of associations with helper T-cell mediators.

## 4 Discussion

In this paper we explored the relationships between key immune mediators of COVID-19 *cytokine storm* and cardiometabolic proteins measured by OLINK Explore platform. ^12^ Using predictive modeling, we integrated Th1, Th2 and Th17 cytokines and found 186/355 significant cardiometabolic predictions. Of these 186 cardiometabolic proteins, 31 were also significantly associated with COVID-19 severity. We highlighted 20 of these 31 proteins that were also associated with cardiovascular inflammation and hypertension in previous studies. For example, LTBP2 has been previously reported as a marker of human heart failure ^18^. CHI3L1 levels have been correlated with severity of coronary disease ^19^ and carotid atherosclerotic plaque, ^20^ as well as stroke ^21^. Elevated GDF15 levels have been associated with higher risks in multiple cardiovascular diseases such as stable coronary artery disease, acute coronary syndrome, and heart failure ^22^. Elevated CXCL8 expression levels have been identified in cases of atherosclerotic plaque ^23^. SPP1 expression has been found to be higher in response to ischemia associated with stroke ^24^, myocardial infarction ^25^, and peripheral artery disease ^26^. FABP4 has been found to contribute to the development of atherosclerosis, and studies had shown that lower levels of FABP4 protect against atherosclerosis to a degree ^38^. REG1A has been shown to have high levels of expression in hearts of patients who died of myocardial infarction ^27^. TFF3 levels in sera have been linked to the prediction of major adverse cardiovascular events ^28^ as well as being identified as a possible biomarker for myocardial infarction ^29^. TNC has been previously linked to many cardiovascular diseases in humans such as pulmonary thromboembolism ^30^ and hypertension ^31^. IL1RL1 has been studied as a marker of cardiac disease, and found to have increased expression in the lung of patients with heart failure ^32^. CSTB has been identified as a relevant biomarker associated with chronic heart failure patients ^33^. NT-proBNP has been shown to be a reliable biomarker in diagnostic evaluation and outcome prediction in cases of acute heart failure, especially in dyspnoeic patients ^34,35^. One study suggested that NTproBNP been seen as a signal of heart failure, valvular heart diseases, pulmonary hypertension ^36^. Higher IGFBP1 expression has been previously related to lessened cardiovascular risk factors and decreased presence of atherosclerosis in elderly patients ^37^.

Our study highlights the potential role of helper T-cells in the production of cardiometabolic proteins in SARS-CoV-2 infection, suggesting the association of the severe *cytokine storm* with cardiovascular-associated complications. While Type 1 and Type 17 helper T-cells have been extensively associated with the immune system’s response to SARS-CoV-2 infection, the role of Type 2 helper T-cells is still poorly understood. Type 2 inflammation characterizes allergic and autoimmune reactions and has been previously associated with an increased risk of vascular inflammation. ^10,11^ In addition, Th2 inhibition has been suggested to offer protection against COVID-19 symptoms ^17,39^, and hence Th2 inhibitors are currently being tested in cases of patients with atopic conditions ^9^. Our study suggests that in synergy with Th1, both Th2 and Th17 pathways play an important role in the over-production of cardiovascular-associated proteins. While Th1 cytokines are important mediators in fighting against any viral infection, Th2 and Th17 pathways are signals triggered by the severe autoimmune response to the unknown pathogen. Hence inhibiting Th2 and Th17 cytokines specifically with immunomodulators may help reduce cardiovascular inflammation without reducing the body’s immune capabilities to fight the infection.

We acknowledge the use of a limited OLINK assay analysis rather than a whole genome analysis, and the potential misdiagnosis of COVID-19 cases during the first wave of the pandemic as limitations of our study.

In summary, while additional research to explore the link between helper T-cells and cardiovascular inflammation is needed, our data suggests that major immune axes (Th1, Th2 and Th17 pathways) are synergistically linked to a myriad of cardiometabolic proteins, which may potentially explain the cardiovascular complications associated with *cytokine storm* in COVID-19 patients. Future work will include further investigation of the relationships between immune and cardiovascular pathways and a detailed comparisons with other viral infections and autoimmune conditions to create a more complete map of immune-cardiovascular interaction of human body.

## Author Contributions

JRM contributed to statistical analysis, data visualization and writing the manuscript. MSN and SA contributed to data visualization. DW provided valuable insight into the methodology. ABP designed the study, performed statistical analysis and data interpretation, and wrote the manuscript.

## Acknowledgements

The authors would like to a acknowledge MGH for making the COVID-19 registry available and the Department of Biomedical Engineering at UM for in kind support.

## Notes and references

1 H. D. E. Dong and L. Gardner, The Lancet, 2020, 20, 533–534.

2 Y. Yang, C. Shen, J. Li, J. Yuan, M. Yang, F. Wang, G. Li, Y. Li, L. Xing, L. Peng, J. Wei, M. Cao, H. Zheng, W. Wu, R. Zou, D. Li, Z. Xu, H. Wang, M. Zhang, Z. Zhang, L. Liu and Y. Liu, medRxiv, 2020.

3 C. T. L. W. L. Yu, H. S. Toh and W. T. Chang, Acta Cardiologica Sinica, 2021, 37, 9–17.

4 D. Weiskope, K. S. Schmitz, M. P. Raadsen, A. Grifoni, N. M. A. Okba, H. Endeman, J. P. C. V. D. Akker, R. Molenkamp, M. P. G. Koopmans, E. C. M. V. Gorp, B. L. Haagmans, R. L. D. Swart, A. Sette and A. R. D. D. Vries, Science Immunology, 2020, 5, –.

5 D. Wu and X. O. Yang, Journal of Microbiology, Immunology, and Infection, 2020, 53, 368–370.

6 J. J. M. Wong, J. Y. Leong, J. H. Lee, S. Albani and J. G. Yeo, Annals of translational medicine, 2019, 7, 19.

7 C. Huang, Y. Wang, X. Li, L. Ren, J. Zhao, Y. Hu, L. Zhang, M. G. Fan, M. J. Xu, M. X. Gu, Z. Cheng, T. Yu, J. Xia, Y. Wei, W. Wu, X. Xie, W. Yin, H. Li, M. Liu, Y. Xiao, H. Gao, L. Guo, J. Xie, G. Wang, R. Jiang, Z. Gao, Q. Jin, J. Wang and B. Cao, The Lancet, 2019, 395, 497–506.

8 A. Pavel, J. W. Glickman, J. R. Michels, S. K. Schulze, R. L. Miller and E. G. Yassky, Frontiers in Genetics, 2021, 12, 10223.

9 B. Ungar, J. W. Glickman, A. K. Golant, C. Dubin, O. Marushchak, A. Gontzes, D. Mikhaylov, G. K. Singer, D. Baum, N. Wei, A. Sanin, D. Gruenstein, M. G. Lebwohl, A. B. Pavel and E. G. Yassky, The Journal of Allergy and Clinical Immunology, 2021.

10 A. P. Villani, A. B. Pavel, J. Wu, M. Fernandes, C. Maari, E. S. C. Proulx, J. G. C. Jack, S. Choi, H. He, B. Ungar, Y. Estrada, N. Kameyama, N. Zhang, J. Gonzales, J. C. Tardif, J. G. Krueger, R. Bissonnette and E. G. Yassky, Wiley Online Library, 2021, 76, 3107–3121.

11 B. Ungar, A. B. Pavel, P. M. Robson, A. Kaufman, A. Pruzan, P. Brunner, S. Kaushik, J. G. Krueger, M. G. Lebwohl, V. Mani, Z. A. Fayad and E. G. Yassky, The Journal of Allergy and Clinical Immunology In Practice, 2020, 8, 3500–3506.

12 M. R. Filbin, A. Mehta, A. M. Schneider, K. R. Kays, J. R. Guess, M. Gentili, B. G. Fenyves, N. C. Charland, A. L. Gonye, I. Gushterova, H. K. Khanna, T. J. LaSalle, K. M. Lavin-Parsons, B. M. Lilly, C. L. Lodenstein, K. Manakongtreecheep, J. D. Margolin, B. N. McKaig, M. Rojas-Lopez, B. C. Russo, N. Sharma, J. Tantivit, M. F. Thomas, R. E. Gerszten, G. S. Heimberg, P. J. Hoover, D. J. Lieb, B. Lin, D. Ngo, K. Pelka, M. Reyes, C. S. Smillie, A. Waghray, T. E. Wood, A. S. Zajac, L. L. Jennings, I. Grundberg, R. P. Bhattacharyya, B. A. Parry, A. C. Villani, M. S. Feldman, N. Hacohen and M. B. Goldberg, Cell Reports Medicine, 2021, 2,.

13 J. F. B. Martin, R. O. D. Lejarazu, T. Pumarola, J. Rello, R. Almansa, P. Ramírez, I. M. Loeches, D. Varillas, M. C. Gallegos, C. Serón, D. Micheloud, J. Gomez, A. T. Abreu, M. J. Ramos, M. Molina, S. Huidobro, E. Sanchez, M. Gordón, V. Fernández, A. D. Castillo, M. Marcos, B. Villanueva, C. López, M. R. Domínguez, J. C. Galan, R. Cantón, A. Lietor, S. Rojo, J. M. Eiros, C. Hinojosa, I. Gonzalez, N. Torner, D. Banner, A. Leon, P. Cuesta, T. Rowe and D. J. Kelvin, Critical care, 2009, 13, R201.

14 S. Weihler and D. Proud, American Journal of Physiology-Lung Cellular and Molecular Physiology, 2007.

15 A. Badawii and S. G. Ryoo, International journal of infectious diseases, 2016, 49, 129–133.

16 J. Leotte, H. Trombetta, H. Z. Faggion, B. M. Almeida, M. B. Nogueira, L. R. Vidal and S. M. Raboni, Jornal de Pediatria, 2017, 93, 294–300.

17 A. B. Pavel, L. Zhou, A. Diaz, B. Ungar, J. Dan, H. He, Y. D. Estrada, H. Xu, M. Fernandes, Y. Renert-Yuval, J. G. Krueger and E. Guttman-Yassky, American Academy of Dermatology, 2022, 82, 690–699.

18 Y. Bai, P. Zhang, X. Zhang, J. Huang, S. Hu and Y. Wei, Biomarkers, 2012, 17, 407–415.

19 K. Sciborski, W. Kuliczkowski, B. Karolko, D. Bednarczyk, M. Protasiewicz, A. Mysiak and M. N. Kawecka, Pol Arch Intern Med, 2018, 18,.

20 A. E. Michelsen, C. N. Rathcke, M. Skjelland, S. Holm, T. Ranheim, K. K. Sørensen, M. F. Klingvall, F. Brosstad, E. Øie, H. Vestergaard, P. Aukrust and B. Halvorsen, Atherosclerosis, 2010, 211, 589–595.

21 P. M. Ridker, D. I. Chasman, L. Rose, J. Loscalzo and J. A. Elias, J Am Heart Assoc, 2014, 114,.

22 T. Kempf and K. C. Wallet, Hertz, 2009, 34, 594–599.

23 H. G. Rus, R. Vlaicu and F. Niculescu, Atherosclerosis, 1996, 127, 263–271.

24 Q. Zhu, X. Luo, J. Zhang, Y. Liu, H. Luo, Q. Huang, Y. Cheng and Z. Xie, Curr Drug Deliv, 2017, 14,.

25 O. Muller, L. Delrue, M. Hamilos, S. Vercauteren, A. Ntalianis, C. Trana, F. Mangiacapra, K. Dierickx, B. D. Bruyne, W. Wijns, A. Behfar, E. Barbato, A. Terzic, M. Vanderheyden and J. Bartunek, Clinical Research, 2011, 7,.

26 M. Koshikawa, K. Aizawa, H. Kasai, A. Izawa, T. Tomita, S. Kumazaki, H. Tsutsui, J. Koyama, S. Shimodaira, M. Takahashi and U. Keda, Sage Journals, 2009, 60, 42–45.

27 T. Kiji, Y. Dohi, S. Takasawa, H. Okamoto, A. Nonomura and S. Taniguchi, Am. J. Physiol. Heart Circ Physiol, 2005.

28 F. Obendorf, C. T. Herz, C. Höbaus, G. Pesau, R. Koppensteiner and G. Schernthaner, Vascular disease, 2015, 132,.

29 A. F. Cisnal, S. G. Blas, E. Valero, G. Miñana, J. Núñez and J.S. Forés, Science Direct, 2020, 73, 418–420.

30 A. Celik, I. Kocyigit, B. Calapkorur, H. Korkmaz, E. Doganay, D. Elcik and I. Ozdogrul, Journal of Ather. Throm., 2011, 18, 487–493.

31 C. Schumann, P. M. Lepper, H. Frank, R. Schneiderbauer, W.T C. Kropf, K. M. Stoiber, S. Rüdiger, L. Kruska, T. Krahn and F. Kramer, Biomarkers, 2010, 15, 523–532.

32 D. A. Pascual-Figal, M.T. Pérez-Martínez, M. C. Asensio-Lopez, J. Sanchez-Más, M.E. García-García, C. M. Martinez, M. Lencina, R. Jara, J. L. Januzzi and A. Lax, Circul. Heart Failure, 2018, 11,.

33 E. Bouwens, M. Brankovic, H. Mouthaan, S. Baart, D. Rizopoulos, N. van Boven, K. Caliskan, O. Manintveld, T. Germans, J. V. Ramshorst, V. Umans, K. M. Akkerhuis and I. Kardys, Jour. of Ame. Heart Ass., 2019, 8,.

34 V. Panagopoulou, S. Deftereos, C. Kossyvakis, K. Raisakis, G. Giannopoulos, G. Bouras, V. Pyrgakis and M. W. Cleman, Curr Top Med Chem., 2013, 13,.

35 J. L. Januzzi, R. V. Kimmenade, J. Lainchbury, A. Bayes-Genis, J. Ordonez-Llanos, M. Santalo-Bel, Y. M. Pinto and M. Richards, European Heart Journal, 2006, 27, 330–337.

36 V. Kimmende and Januzzi, Biomarker Insights, 2006, 1,.

37 J. A. M. J. L. Janssen, R. P. Stolk, H. A. P. Pols, D. E. Grobbee and S. W. J. Lamberts, Arterioscler Thromb Vasc Biol, 1998, 18, 277–282.

38 L. Makowski, J. B. Boord, K. Maeda, V. R. Babaev, K. T. Uysal, M. A. Morgan, R. A. Parker, J. Suttles, S. Fazio, G. S. Hotamisligil and M. F. Linton, naturemedicine, 2001, 7, 699–705.

39 A. C. C. C. Branco, M. N. Sato and R. W. Alberca, Frontiers in cellular and infection microbiology, 2020, 106,.

