## Supplementary Tables for "Th1, Th2 and Th17 inflammatory pathways synergistically correlate with cardiometabolic processes. A case study in COVID-19"

**Supplementary Table 1. Statistical comparisons between deceased, intubated and non-severe groups in COVID-19 positive patients for all Th1, Th2 and Th17 cytokines and immune mediators.**

**A. Th1**

| marker | Mean_Estimate_Deceased | Mean_Estimate_Intubated | Mean_Estimate_NonSevere | lgFCH_Deceased_vs_Intubated | FCH_Deceased_vs_Intubated | p_Deceased_vs_Intubated | lgFCH_Deceased_vs_NonSevere | FCH_Deceased_vs_NonSevere | p_Deceased_vs_NonSevere | lgFCH_Intubated_vs_NonSevere | FCH_Intubated_vs_NonSevere | p_Intubated_vs_NonSevere |
|---|---|---|---|---|---|---|---|---|---|---|---|---|
| CCL3 | 3.839667 | 3.758557 | 3.375698 | 0.08111 | 1.057832 | 0.632851 | 0.463969 | 1.379331 | 0.001702 | 0.382859 | 1.303923 | 0.001862 |
| CCL4 | 4.533055 | 4.450697 | 4.183864 | 0.082358 | 1.058747 | 0.609644 | 0.349191 | 1.273846 | 0.012653 | 0.266833 | 1.203164 | 0.021959 |
| CCL5 | 3.058536 | 3.497901 | 3.521671 | -0.43937 | -1.35601 | 0.089238 | -0.46314 | -1.37853 | 0.038352 | -0.02377 | -1.01661 | 0.898016 |
| CXCL10 | 8.067081 | 8.027064 | 7.060412 | 0.040017 | 1.028126 | 0.869028 | 1.006669 | 2.009266 | 2.44E-06 | 0.966652 | 1.9543 | 6.45E-08 |
| CXCL11 | 5.420621 | 5.473434 | 5.093661 | -0.05281 | -1.03729 | 0.79439 | 0.326961 | 1.254368 | 0.062564 | 0.379774 | 1.301138 | 0.009544 |
| CXCL9 | 4.755986 | 4.626119 | 3.958236 | 0.129866 | 1.094192 | 0.541683 | 0.79775 | 1.738388 | 1.91E-05 | 0.667884 | 1.588741 | 1.71E-05 |
| IFNG | 2.912634 | 3.200185 | 3.559888 | -0.28755 | -1.22057 | 0.444137 | -0.64725 | -1.56618 | 0.047345 | -0.3597 | -1.28316 | 0.180299 |
| IFNGR1 | 1.735405 | 1.364157 | 1.174299 | 0.371248 | 1.293471 | 0.000109 | 0.561105 | 1.475399 | 3.87E-11 | 0.189857 | 1.140651 | 0.005615 |
| IFNGR2 | 2.288057 | 2.554254 | 2.214384 | -0.2662 | -1.20263 | 0.037079 | 0.073673 | 1.052393 | 0.502824 | 0.33987 | 1.265642 | 0.000239 |
| IL1B | 0.972867 | 0.715585 | 0.551015 | 0.257282 | 1.195224 | 0.027092 | 0.421851 | 1.339646 | 3.30E-05 | 0.16457 | 1.120832 | 0.049095 |
| IL2RA | 5.35906 | 5.168139 | 4.904144 | 0.190921 | 1.141492 | 0.145873 | 0.454916 | 1.370703 | 7.32E-05 | 0.263995 | 1.200799 | 0.005378 |

**B. Th2**

| marker | Mean_Estimate_Deceased | Mean_Estimate_Intubated | Mean_Estimate_NonSevere | lgFCH_Deceased_vs_Intubated | FCH_Deceased_vs_Intubated | p_Deceased_vs_Intubated | lgFCH_Deceased_vs_NonSevere | FCH_Deceased_vs_NonSevere | p_Deceased_vs_NonSevere | lgFCH_Intubated_vs_NonSevere | FCH_Intubated_vs_NonSevere | p_Intubated_vs_NonSevere |
|---|---|---|---|---|---|---|---|---|---|---|---|---|
| CCL11 | 6.867267 | 6.340584 | 6.343758 | 0.526683 | 1.440613 | 0.000114 | 0.523509 | 1.437447 | 9.80E-06 | -0.00317 | -1.0022 | 0.973879 |
| CCL13 | 6.441117 | 6.23976 | 6.163189 | 0.201357 | 1.149779 | 0.204937 | 0.277927 | 1.212452 | 0.043277 | 0.07657 | 1.054508 | 0.502207 |
| CCL17 | 4.20771 | 4.247852 | 4.606879 | -0.04014 | -1.02822 | 0.845314 | -0.39917 | -1.31875 | 0.02533 | -0.35903 | -1.28256 | 0.015733 |
| CCL22 | 5.19659 | 5.28663 | 5.448229 | -0.09004 | -1.0644 | 0.488051 | -0.25164 | -1.19056 | 0.025435 | -0.1616 | -1.11853 | 0.084137 |
| CCL24 | 5.229133 | 4.652407 | 4.639222 | 0.576726 | 1.491461 | 0.031271 | 0.589911 | 1.505154 | 0.010901 | 0.013185 | 1.009181 | 0.9452 |
| CCL26 | 1.947276 | 1.858636 | 1.743453 | 0.08864 | 1.063368 | 0.668685 | 0.203824 | 1.151747 | 0.255128 | 0.115183 | 1.083113 | 0.439454 |
| CCL7 | 5.453148 | 5.706034 | 4.334088 | -0.25289 | -1.19159 | 0.35047 | 1.11906 | 2.172054 | 2.62E-06 | 1.371947 | 2.588195 | 1.18E-11 |
| TSLP | 0.124193 | 0.165269 | 0.166628 | -0.04108 | -1.02888 | 0.640193 | -0.04244 | -1.02985 | 0.576227 | -0.00136 | -1.00094 | 0.982833 |
| IL10 | 0.392479 | 0.395799 | 0.121696 | -0.00332 | -1.0023 | 0.989468 | 0.270782 | 1.206462 | 0.213273 | 0.274102 | 1.209241 | 0.130328 |
| IL13 | -0.18518 | -0.08663 | -0.08922 | -0.09855 | -1.0707 | 0.303685 | -0.09596 | -1.06878 | 0.246463 | 0.00259 | 1.001797 | 0.969987 |
| IL33 | 0.145867 | 0.17389 | 0.053408 | -0.02802 | -1.01961 | 0.716487 | 0.092459 | 1.066186 | 0.166098 | 0.120482 | 1.087098 | 0.030523 |
| IL4R | 3.974879 | 3.615075 | 3.073927 | 0.359804 | 1.283251 | 0.047431 | 0.900952 | 1.867297 | 1.97E-08 | 0.541148 | 1.45513 | 4.09E-05 |
| IL5 | 0.087957 | 0.389097 | 0.188048 | -0.30114 | -1.23212 | 0.171847 | -0.10009 | -1.07184 | 0.598658 | 0.201049 | 1.149534 | 0.204498 |
| IL7R | 4.057112 | 3.708964 | 3.962354 | 0.348148 | 1.272925 | 0.008226 | 0.094758 | 1.067886 | 0.402652 | -0.25339 | -1.192 | 0.007484 |

**C. Th17**

| marker | Mean_Estimate_Deceased | Mean_Estimate_Intubated | Mean_Estimate_NonSevere | lgFCH_Deceased_vs_Intubated | FCH_Deceased_vs_Intubated | p_Deceased_vs_Intubated | lgFCH_Deceased_vs_NonSevere | FCH_Deceased_vs_NonSevere | p_Deceased_vs_NonSevere | lgFCH_Intubated_vs_NonSevere | FCH_Intubated_vs_NonSevere | p_Intubated_vs_NonSevere |
|---|---|---|---|---|---|---|---|---|---|---|---|---|
| S100A12 | 6.28179 | 6.370781 | 5.712131 | -0.08899 | -1.06363 | 0.517601 | 0.569659 | 1.484173 | 2.50E-06 | 0.658649 | 1.578604 | 1.23E-10 |
| S100P | 2.406164 | 2.023994 | 1.42661 | 0.38217 | 1.303301 | 0.032771 | 0.979554 | 1.971856 | 7.32E-10 | 0.597384 | 1.51297 | 4.70E-06 |
| CCL20 | 6.250036 | 5.6281 | 4.885216 | 0.621936 | 1.538939 | 0.0198 | 1.364819 | 2.575441 | 7.41E-09 | 0.742884 | 1.673518 | 0.000123 |
| LCN2 | 3.086124 | 2.520122 | 2.200199 | 0.566001 | 1.480415 | 0.000796 | 0.885925 | 1.847949 | 2.61E-09 | 0.319923 | 1.248264 | 0.008147 |
| PI3 | 2.858407 | 1.784919 | 1.671535 | 1.073488 | 2.104515 | 2.17E-05 | 1.186872 | 2.276586 | 7.24E-08 | 0.113384 | 1.081763 | 0.526691 |
| CXCL1 | 3.875114 | 3.572672 | 3.533874 | 0.302443 | 1.233231 | 0.111447 | 0.34124 | 1.266845 | 0.037924 | 0.038798 | 1.027257 | 0.775992 |
| CXCL3 | 4.791669 | 4.784497 | 5.078272 | 0.007172 | 1.004984 | 0.97909 | -0.2866 | -1.21976 | 0.225943 | -0.29377 | -1.22584 | 0.136135 |
| IL12A | 4.276736 | 4.333357 | 4.444208 | -0.05662 | -1.04003 | 0.782909 | -0.16747 | -1.12309 | 0.345851 | -0.11085 | -1.07987 | 0.453321 |
| IL12B | 5.547005 | 5.566451 | 5.679793 | -0.01945 | -1.01357 | 0.918413 | -0.13279 | -1.09641 | 0.4184 | -0.11334 | -1.08173 | 0.406623 |
| IL17A | 0.827764 | 0.410578 | 0.449506 | 0.417187 | 1.335321 | 3.26E-05 | 0.378258 | 1.299772 | 1.34E-05 | -0.03893 | -1.02735 | 0.584485 |
| IL17F | 0.561443 | 0.381466 | 0.410569 | 0.179977 | 1.132866 | 0.223551 | 0.150874 | 1.110242 | 0.237566 | -0.0291 | -1.02038 | 0.784056 |
| IL6 | 7.188969 | 6.763184 | 5.177767 | 0.425785 | 1.343304 | 0.144768 | 2.011202 | 4.03118 | 2.80E-14 | 1.585417 | 3.000945 | 4.53E-13 |
| IL6R | 4.358988 | 4.280809 | 4.312342 | 0.078179 | 1.055685 | 0.402947 | 0.046646 | 1.032861 | 0.563405 | -0.03153 | -1.0221 | 0.638831 |

**Supplementary Table 2. Statistical comparisons between deceased, intubated and non-severe in COVID-19 negative patients with other respiratory conditions for all Th1, Th2 and Th17 cytokines and immune mediators.**

## A. Th1

| marker | Mean_Estimate_Deceased | Mean_Estimate_Intubated | Mean_Estimate_NonSevere | lgFCH_Deceased_vs_Intubated | FCH_Deceased_vs_Intubated | p_Deceased_vs_Intubated | lgFCH_Deceased_vs_NonSevere | FCH_Deceased_vs_NonSevere | p_Deceased_vs_NonSevere | lgFCH_Intubated_vs_NonSevere | FCH_Intubated_vs_NonSevere | p_Intubated_vs_NonSevere |
|---|---|---|---|---|---|---|---|---|---|---|---|---|
| CCL3 | 3.8184143 | 3.478556 | 3.511958 | 0.339858036 | 1.26563 | 0.499544 | 0.306456 | 1.236666 | 0.491767 | -0.0334 | -1.02342 | 0.915559 |
| CCL4 | 4.6149 | 4.609681 | 4.569096 | 0.00521875 | 1.00362 | 0.991049 | 0.045804 | 1.032258 | 0.91147 | 0.040585 | 1.028531 | 0.889309 |
| CCL5 | 1.8490571 | 3.40845 | 3.02596 | -1.55939286 | -2.9473 | 0.007281 | -1.1769 | -2.26091 | 0.021332 | 0.38249 | 1.30359 | 0.283761 |
| CXCL10 | 5.2372857 | 4.696006 | 5.052038 | 0.541279464 | 1.45526 | 0.372925 | 0.185248 | 1.137012 | 0.729995 | -0.35603 | -1.2799 | 0.349913 |
| CXCL11 | 3.4802857 | 3.755888 | 3.691851 | -0.27560179 | -1.2105 | 0.639699 | -0.21157 | -1.15794 | 0.684838 | 0.064037 | 1.045387 | 0.86215 |
| CXCL9 | 4.3315286 | 4.049456 | 4.437155 | 0.282072321 | 1.21594 | 0.664552 | -0.10563 | -1.07596 | 0.854434 | -0.3877 | -1.3083 | 0.342817 |
| IFNG | 0.8061571 | 0.217613 | 1.646167 | 0.588544643 | 1.50373 | 0.622502 | -0.84001 | -1.79006 | 0.428101 | -1.42855 | -2.69177 | 0.05939 |
| IFNGR1 | 1.7429286 | 1.69275 | 1.563549 | 0.050178571 | 1.03539 | 0.864894 | 0.179379 | 1.132397 | 0.49281 | 0.129201 | 1.093688 | 0.48526 |
| IFNGR2 | 2.0041143 | 1.9355 | 1.82002 | 0.068614286 | 1.04871 | 0.80086 | 0.184094 | 1.136104 | 0.445526 | 0.11548 | 1.083335 | 0.498799 |
| IL1B | 0.4707571 | 1.017263 | 0.784813 | -0.54650536 | -1.4605 | 0.188913 | -0.31406 | -1.2432 | 0.392315 | 0.23245 | 1.174828 | 0.37116 |
| IL2RA | 5.1694 | 5.060188 | 5.123436 | 0.1092125 | 1.07864 | 0.764208 | 0.045964 | 1.032373 | 0.886612 | -0.06325 | -1.04482 | 0.781658 |

**B. Th2**

| marker | Mean_Estimate_Deceased | Mean_Estimate_Intubated | Mean_Estimate_NonSevere | lgFCH_Deceased_vs_Intubated | FCH_Deceased_vs_Intubated | p_Deceased_vs_Intubated | lgFCH_Deceased_vs_NonSevere | FCH_Deceased_vs_NonSevere | p_Deceased_vs_NonSevere | lgFCH_Intubated_vs_NonSevere | FCH_Intubated_vs_NonSevere | p_Intubated_vs_NonSevere |
|---|---|---|---|---|---|---|---|---|---|---|---|---|
| CCL11 | 6.5765 | 7.028294 | 6.53106 | -0.45179 | -1.36774 | 0.198314 | 0.04544 | 1.031998 | 0.883207 | 0.497234 | 1.411505 | 0.025523 |
| CCL13 | 6.098514 | 6.262263 | 6.223949 | -0.16375 | -1.12019 | 0.690231 | -0.12543 | -1.09084 | 0.730247 | 0.038313 | 1.026913 | 0.881681 |
| CCL17 | 4.779471 | 5.398713 | 5.340918 | -0.61924 | -1.53607 | 0.234315 | -0.56145 | -1.47575 | 0.223448 | 0.057794 | 1.040873 | 0.858792 |
| CCL22 | 4.951286 | 5.417663 | 5.598204 | -0.46638 | -1.38164 | 0.166933 | -0.64692 | -1.56582 | 0.031923 | -0.18054 | -1.13331 | 0.391456 |
| CCL24 | 6.521729 | 6.846756 | 5.876842 | -0.32503 | -1.25269 | 0.564903 | 0.644887 | 1.563617 | 0.199176 | 0.969914 | 1.958724 | 0.007415 |
| CCL26 | 1.706957 | 1.674981 | 1.337822 | 0.031976 | 1.022411 | 0.945263 | 0.369135 | 1.291578 | 0.372048 | 0.337159 | 1.263267 | 0.250208 |
| CCL7 | 3.133514 | 2.633488 | 2.609289 | 0.500027 | 1.41424 | 0.282295 | 0.524225 | 1.438161 | 0.203751 | 0.024198 | 1.016915 | 0.933587 |
| TSLP | 0.175114 | 0.173731 | 0.164376 | 0.001383 | 1.000959 | 0.995387 | 0.010738 | 1.007471 | 0.95957 | 0.009355 | 1.006505 | 0.950249 |
| IL10 | 0.288514 | -0.03083 | 0.399467 | 0.319346 | 1.247764 | 0.464754 | -0.11095 | -1.07994 | 0.773917 | -0.4303 | -1.34751 | 0.11841 |
| IL13 | -0.17047 | -0.24991 | -0.02553 | 0.079441 | 1.056609 | 0.763805 | -0.14494 | -1.10569 | 0.53624 | -0.22439 | -1.16828 | 0.178217 |
| IL33 | 0.017157 | 0.223506 | 0.166816 | -0.20635 | -1.15376 | 0.319631 | -0.14966 | -1.10931 | 0.41451 | 0.05669 | 1.040077 | 0.661841 |
| IL4R | 3.350886 | 3.553581 | 3.258693 | -0.2027 | -1.15085 | 0.668005 | 0.092193 | 1.065989 | 0.82557 | 0.294889 | 1.22679 | 0.320813 |
| IL5 | 0.356757 | 0.340706 | 0.693925 | 0.016051 | 1.011188 | 0.976996 | -0.33717 | -1.26327 | 0.494641 | -0.35322 | -1.27741 | 0.313012 |
| IL7R | 4.729986 | 4.626925 | 4.440115 | 0.103061 | 1.07405 | 0.77501 | 0.289871 | 1.222531 | 0.365154 | 0.18681 | 1.138244 | 0.409421 |

**C. Th17**

| marker | Mean_Estimate_Deceased | Mean_Estimate_Intubated | Mean_Estimate_NonSevere | lgFCH_Deceased_vs_Intubated | FCH_Deceased_vs_Intubated | p_Deceased_vs_Intubated | lgFCH_Deceased_vs_NonSevere | FCH_Deceased_vs_NonSevere | p_Deceased_vs_NonSevere | lgFCH_Intubated_vs_NonSevere | FCH_Intubated_vs_NonSevere | p_Intubated_vs_NonSevere |
|---|---|---|---|---|---|---|---|---|---|---|---|---|
| S100A12 | 5.728757 | 6.064206 | 5.852045 | -0.33545 | -1.26177 | 0.351617 | -0.12329 | -1.08921 | 0.698372 | 0.212161 | 1.158422 | 0.347308 |
| S100P | 2.959171 | 2.700819 | 2.074989 | 0.258353 | 1.196112 | 0.609955 | 0.884182 | 1.845718 | 0.05139 | 0.62583 | 1.543098 | 0.051394 |
| CCL20 | 5.498871 | 5.127169 | 5.28746 | 0.371703 | 1.293879 | 0.638851 | 0.211411 | 1.15782 | 0.762991 | -0.16029 | -1.11751 | 0.746693 |
| LCN2 | 2.941471 | 3.17935 | 2.905856 | -0.23788 | -1.17926 | 0.651348 | 0.035615 | 1.024994 | 0.939059 | 0.273494 | 1.208731 | 0.407929 |
| PI3 | 2.871914 | 2.453788 | 2.236811 | 0.418127 | 1.336191 | 0.465291 | 0.635103 | 1.553049 | 0.211994 | 0.216977 | 1.162295 | 0.545291 |
| CXCL1 | 3.8285 | 3.342113 | 3.150798 | 0.486388 | 1.400933 | 0.4052 | 0.677702 | 1.59959 | 0.191815 | 0.191314 | 1.141803 | 0.600926 |
| CXCL3 | 3.810729 | 4.569119 | 4.626156 | -0.75839 | -1.6916 | 0.278115 | -0.81543 | -1.75982 | 0.188707 | -0.05704 | -1.04033 | 0.896059 |
| IL12A | 3.302486 | 4.294444 | 4.566915 | -0.99196 | -1.98888 | 0.096154 | -1.26443 | -2.40232 | 0.017698 | -0.27247 | -1.20787 | 0.462585 |
| IL12B | 4.511771 | 5.376981 | 5.739651 | -0.86521 | -1.8216 | 0.128247 | -1.22788 | -2.34222 | 0.015995 | -0.36267 | -1.2858 | 0.306998 |
| IL17A | 0.796386 | 0.63395 | 0.743722 | 0.162436 | 1.119175 | 0.53716 | 0.052664 | 1.037178 | 0.821057 | -0.10977 | -1.07906 | 0.505924 |
| IL17F | 0.540414 | 0.308581 | 0.549616 | 0.231833 | 1.174326 | 0.581332 | -0.0092 | -1.0064 | 0.980261 | -0.24104 | -1.18184 | 0.361212 |
| IL6 | 5.299943 | 5.773325 | 4.979655 | -0.47338 | -1.38836 | 0.659955 | 0.320288 | 1.24858 | 0.736682 | 0.79367 | 1.733479 | 0.241117 |
| IL6R | 4.077571 | 4.296531 | 4.175715 | -0.21896 | -1.16389 | 0.369367 | -0.09814 | -1.07039 | 0.648915 | 0.120817 | 1.08735 | 0.429127 |

**Supplementary Table 3. Statistical comparisons between deceased, intubated and non-severe groups in COVID-19 positive patients for all cardiometabolic proteins.**

| marker | Mean_Estimate_Deceased | Mean_Estimate_Intubated | Mean_Estimate_NonSevere | lgFCH_Deceased_vs_Intubated | FCH_Deceased_vs_Intubated | p_Deceased_vs_Intubated | lgFCH_Deceased_vs_NonSevere | FCH_Deceased_vs_NonSevere | p_Deceased_vs_NonSevere | lgFCH_Intubated_vs_NonSevere | FCH_Intubated_vs_NonSevere | p_Intubated_vs_NonSevere | FDR_Deceased_vs_Intubated | FDR_Deceased_vs_NonSevere | FDR_Intubated_vs_NonSevere |
|---|---|---|---|---|---|---|---|---|---|---|---|---|---|---|---|
| QDPR | 3.52 | 3.74 | 3.22 | -0.21 | -1.16 | 0.21 | 0.30 | 1.23 | 0.04 | 0.51 | 1.43 | 0.00 | 0.36 | 0.07 | 0.00 |
| QPCT | 3.56 | 3.26 | 3.08 | 0.30 | 1.23 | 0.03 | 0.49 | 1.40 | 0.00 | 0.18 | 1.14 | 0.07 | 0.09 | 0.00 | 0.13 |
| RARRES2 | 2.65 | 2.58 | 2.56 | 0.08 | 1.05 | 0.52 | 0.10 | 1.07 | 0.35 | 0.02 | 1.01 | 0.82 | 0.66 | 0.42 | 0.86 |
| RNASE3 | 2.79 | 2.26 | 1.67 | 0.53 | 1.45 | 0.04 | 1.12 | 2.18 | 0.00 | 0.59 | 1.50 | 0.00 | 0.11 | 0.00 | 0.01 |
| RNASET2 | 4.74 | 4.61 | 4.35 | 0.14 | 1.10 | 0.17 | 0.39 | 1.31 | 0.00 | 0.25 | 1.19 | 0.00 | 0.32 | 0.00 | 0.00 |
| ROR1 | 0.86 | 0.50 | 0.55 | 0.37 | 1.29 | 0.00 | 0.31 | 1.24 | 0.00 | -0.06 | -1.04 | 0.42 | 0.00 | 0.00 | 0.52 |
| S100A11 | 1.83 | 1.83 | 1.29 | 0.00 | 1.00 | 0.98 | 0.54 | 1.46 | 0.00 | 0.54 | 1.45 | 0.00 | 0.99 | 0.00 | 0.00 |
| SCARF1 | 3.50 | 3.29 | 3.13 | 0.22 | 1.16 | 0.12 | 0.37 | 1.29 | 0.00 | 0.15 | 1.11 | 0.13 | 0.24 | 0.00 | 0.21 |
| RCOR1 | 0.62 | 0.66 | 0.50 | -0.04 | -1.03 | 0.54 | 0.12 | 1.09 | 0.04 | 0.16 | 1.12 | 0.00 | 0.67 | 0.06 | 0.00 |
| REG1A | 6.36 | 5.30 | 5.08 | 1.06 | 2.08 | 0.00 | 1.28 | 2.42 | 0.00 | 0.22 | 1.16 | 0.20 | 0.00 | 0.00 | 0.29 |
| REG1B | 4.69 | 3.61 | 3.31 | 1.08 | 2.12 | 0.00 | 1.39 | 2.62 | 0.00 | 0.31 | 1.24 | 0.10 | 0.00 | 0.00 | 0.18 |
| REG3A | 5.08 | 4.02 | 3.80 | 1.06 | 2.09 | 0.00 | 1.28 | 2.42 | 0.00 | 0.21 | 1.16 | 0.25 | 0.00 | 0.00 | 0.34 |
| REN | 2.80 | 2.19 | 1.85 | 0.61 | 1.53 | 0.00 | 0.95 | 1.93 | 0.00 | 0.34 | 1.26 | 0.02 | 0.01 | 0.00 | 0.05 |
| RETN | 7.18 | 6.79 | 6.15 | 0.39 | 1.31 | 0.03 | 1.03 | 2.04 | 0.00 | 0.64 | 1.56 | 0.00 | 0.08 | 0.00 | 0.00 |
| SEMA3F | 2.19 | 1.82 | 1.64 | 0.37 | 1.29 | 0.00 | 0.55 | 1.46 | 0.00 | 0.18 | 1.13 | 0.02 | 0.00 | 0.00 | 0.06 |
| SEMA7A | 4.74 | 4.61 | 4.39 | 0.13 | 1.09 | 0.25 | 0.35 | 1.28 | 0.00 | 0.22 | 1.17 | 0.01 | 0.40 | 0.00 | 0.02 |
| SERPINA11 | 6.59 | 6.75 | 6.50 | -0.16 | -1.12 | 0.20 | 0.09 | 1.06 | 0.41 | 0.25 | 1.19 | 0.01 | 0.34 | 0.48 | 0.02 |
| SERPINB5 | 0.37 | 0.15 | 0.18 | 0.22 | 1.16 | 0.00 | 0.19 | 1.14 | 0.00 | -0.03 | -1.02 | 0.58 | 0.01 | 0.01 | 0.67 |
| SERPINE1 | 6.68 | 6.86 | 6.25 | -0.18 | -1.13 | 0.40 | 0.43 | 1.35 | 0.02 | 0.61 | 1.53 | 0.00 | 0.54 | 0.03 | 0.00 |
| SFTPD | 3.76 | 3.47 | 3.11 | 0.29 | 1.22 | 0.20 | 0.65 | 1.57 | 0.00 | 0.36 | 1.28 | 0.03 | 0.35 | 0.00 | 0.06 |

| | | | | | | | | | | | | | | | |
|---|---|---|---|---|---|---|---|---|---|---|---|---|---|---|---|
| SDC1 | 4.03 | 3.95 | 3.22 | 0.08 | 1.06 | 0.71 | 0.81 | 1.75 | 0.00 | 0.73 | 1.66 | 0.00 | 0.81 | 0.00 | 0.00 |
| SDC4 | 2.90 | 3.10 | 2.91 | -0.20 | -1.15 | 0.24 | -0.01 | -1.01 | 0.93 | 0.18 | 1.14 | 0.13 | 0.39 | 0.95 | 0.20 |
| SELE | 5.68 | 5.86 | 5.65 | -0.18 | -1.14 | 0.19 | 0.03 | 1.02 | 0.80 | 0.21 | 1.16 | 0.03 | 0.33 | 0.84 | 0.07 |
| SELP | 2.95 | 2.87 | 2.76 | 0.08 | 1.05 | 0.62 | 0.19 | 1.14 | 0.16 | 0.11 | 1.08 | 0.31 | 0.75 | 0.20 | 0.40 |
| SLITRK6 | 2.37 | 2.33 | 2.29 | 0.04 | 1.03 | 0.66 | 0.08 | 1.06 | 0.32 | 0.04 | 1.03 | 0.56 | 0.78 | 0.39 | 0.66 |
| SNAP23 | 3.90 | 3.52 | 3.92 | 0.38 | 1.30 | 0.26 | -0.02 | -1.01 | 0.94 | -0.40 | -1.32 | 0.10 | 0.41 | 0.95 | 0.17 |
| SNX9 | 2.88 | 2.67 | 2.49 | 0.21 | 1.16 | 0.32 | 0.40 | 1.32 | 0.03 | 0.19 | 1.14 | 0.21 | 0.46 | 0.05 | 0.30 |
| SOD1 | 1.94 | 1.98 | 1.51 | -0.04 | -1.03 | 0.82 | 0.43 | 1.34 | 0.01 | 0.47 | 1.39 | 0.00 | 0.89 | 0.02 | 0.00 |
| SORT1 | 4.46 | 4.38 | 4.18 | 0.08 | 1.05 | 0.48 | 0.27 | 1.21 | 0.00 | 0.20 | 1.15 | 0.01 | 0.62 | 0.01 | 0.03 |
| SOST | 5.19 | 4.98 | 4.69 | 0.21 | 1.16 | 0.10 | 0.50 | 1.41 | 0.00 | 0.29 | 1.22 | 0.00 | 0.22 | 0.00 | 0.01 |
| SPARCL1 | 3.79 | 3.39 | 3.28 | 0.40 | 1.32 | 0.00 | 0.51 | 1.43 | 0.00 | 0.11 | 1.08 | 0.11 | 0.00 | 0.00 | 0.19 |
| SPON2 | 5.14 | 4.80 | 4.25 | 0.34 | 1.27 | 0.04 | 0.90 | 1.86 | 0.00 | 0.55 | 1.47 | 0.00 | 0.10 | 0.00 | 0.00 |
| SPP1 | 9.28 | 8.61 | 8.06 | 0.67 | 1.59 | 0.00 | 1.22 | 2.33 | 0.00 | 0.55 | 1.47 | 0.00 | 0.00 | 0.00 | 0.00 |
| SSC4D | 1.90 | 1.99 | 2.47 | -0.08 | -1.06 | 0.75 | -0.56 | -1.48 | 0.01 | -0.48 | -1.40 | 0.01 | 0.83 | 0.02 | 0.03 |
| SSC5D | 2.98 | 3.12 | 3.10 | -0.14 | -1.11 | 0.17 | -0.12 | -1.09 | 0.18 | 0.02 | 1.02 | 0.77 | 0.32 | 0.23 | 0.82 |
| ST6GAL1 | 6.14 | 6.11 | 5.62 | 0.03 | 1.02 | 0.81 | 0.51 | 1.43 | 0.00 | 0.48 | 1.40 | 0.00 | 0.88 | 0.00 | 0.00 |
| STK11 | 2.44 | 2.37 | 1.91 | 0.07 | 1.05 | 0.66 | 0.53 | 1.44 | 0.00 | 0.45 | 1.37 | 0.00 | 0.78 | 0.00 | 0.00 |
| STK4 | 0.36 | 0.36 | 0.39 | 0.00 | 1.00 | 0.99 | -0.02 | -1.02 | 0.80 | -0.03 | -1.02 | 0.74 | 0.99 | 0.84 | 0.81 |
| SUSD1 | 1.06 | 1.06 | 1.06 | 0.00 | -1.00 | 0.99 | -0.01 | -1.00 | 0.89 | -0.01 | -1.00 | 0.89 | 0.99 | 0.92 | 0.91 |
| TCL1B | 0.02 | 0.03 | 0.17 | 0.00 | -1.00 | 0.96 | -0.15 | -1.11 | 0.08 | -0.14 | -1.10 | 0.04 | 0.99 | 0.11 | 0.08 |
| TCN2 | 8.18 | 8.12 | 7.87 | 0.06 | 1.05 | 0.53 | 0.31 | 1.24 | 0.00 | 0.24 | 1.18 | 0.00 | 0.66 | 0.00 | 0.00 |
| TFF3 | 2.51 | 1.79 | 1.46 | 0.71 | 1.64 | 0.00 | 1.05 | 2.07 | 0.00 | 0.33 | 1.26 | 0.05 | 0.01 | 0.00 | 0.09 |
| TFPI | 4.68 | 4.69 | 4.49 | -0.01 | -1.01 | 0.90 | 0.19 | 1.14 | 0.02 | 0.20 | 1.15 | 0.00 | 0.95 | 0.03 | 0.01 |
| TFRC | 2.31 | 2.21 | 2.05 | 0.10 | 1.08 | 0.33 | 0.27 | 1.20 | 0.00 | 0.16 | 1.12 | 0.04 | 0.48 | 0.01 | 0.08 |
| TGFBI | 4.77 | 4.93 | 4.79 | -0.16 | -1.11 | 0.07 | -0.02 | -1.02 | 0.75 | 0.13 | 1.10 | 0.04 | 0.17 | 0.80 | 0.08 |
| TGFBR3 | 3.98 | 3.62 | 3.51 | 0.36 | 1.28 | 0.00 | 0.47 | 1.39 | 0.00 | 0.11 | 1.08 | 0.21 | 0.02 | 0.00 | 0.30 |
| TGM2 | 4.78 | 4.86 | 4.39 | -0.07 | -1.05 | 0.76 | 0.40 | 1.32 | 0.05 | 0.47 | 1.39 | 0.01 | 0.84 | 0.07 | 0.02 |
| THBD | 3.44 | 3.07 | 2.85 | 0.36 | 1.29 | 0.00 | 0.59 | 1.50 | 0.00 | 0.22 | 1.17 | 0.01 | 0.01 | 0.00 | 0.02 |
| THBS4 | 3.65 | 3.21 | 3.06 | 0.43 | 1.35 | 0.01 | 0.59 | 1.50 | 0.00 | 0.16 | 1.11 | 0.21 | 0.04 | 0.00 | 0.31 |
| THOP1 | 5.00 | 5.49 | 4.90 | -0.49 | -1.40 | 0.00 | 0.10 | 1.07 | 0.41 | 0.59 | 1.50 | 0.00 | 0.00 | 0.48 | 0.00 |
| THPO | 1.58 | 1.68 | 1.61 | -0.09 | -1.06 | 0.35 | -0.02 | -1.01 | 0.80 | 0.07 | 1.05 | 0.32 | 0.50 | 0.84 | 0.41 |
| TIA1 | 0.83 | 0.84 | 0.76 | -0.01 | -1.01 | 0.88 | 0.07 | 1.05 | 0.40 | 0.08 | 1.06 | 0.22 | 0.93 | 0.47 | 0.31 |
| TIE1 | 4.50 | 4.31 | 4.39 | 0.19 | 1.14 | 0.04 | 0.12 | 1.08 | 0.15 | -0.08 | -1.05 | 0.25 | 0.10 | 0.19 | 0.35 |
| TIMD4 | 3.27 | 3.07 | 2.87 | 0.20 | 1.15 | 0.17 | 0.39 | 1.31 | 0.00 | 0.20 | 1.15 | 0.06 | 0.32 | 0.00 | 0.12 |

| | | | | | | | | | | | | | | | |
|---|---|---|---|---|---|---|---|---|---|---|---|---|---|---|---|
| TIMP1 | 6.98 | 6.80 | 6.22 | 0.18 | 1.13 | 0.18 | 0.76 | 1.70 | 0.00 | 0.58 | 1.50 | 0.00 | 0.33 | 0.00 | 0.00 |
| TINAGL1 | 4.48 | 4.34 | 3.96 | 0.14 | 1.10 | 0.19 | 0.53 | 1.44 | 0.00 | 0.38 | 1.30 | 0.00 | 0.34 | 0.00 | 0.00 |
| SIGLEC7 | 2.72 | 2.53 | 2.47 | 0.20 | 1.15 | 0.02 | 0.25 | 1.19 | 0.00 | 0.05 | 1.04 | 0.38 | 0.06 | 0.00 | 0.48 |
| SIRPA | 6.47 | 6.21 | 6.08 | 0.26 | 1.20 | 0.08 | 0.39 | 1.31 | 0.00 | 0.13 | 1.10 | 0.22 | 0.19 | 0.00 | 0.31 |
| PTGDS | 4.72 | 4.06 | 4.08 | 0.66 | 1.58 | 0.00 | 0.64 | 1.56 | 0.00 | -0.02 | -1.01 | 0.90 | 0.00 | 0.00 | 0.91 |
| PTN | 0.55 | -0.03 | -0.25 | 0.58 | 1.50 | 0.00 | 0.80 | 1.74 | 0.00 | 0.22 | 1.16 | 0.06 | 0.00 | 0.00 | 0.12 |
| PTPRF | 3.87 | 3.76 | 3.57 | 0.11 | 1.08 | 0.20 | 0.30 | 1.23 | 0.00 | 0.19 | 1.14 | 0.00 | 0.35 | 0.00 | 0.01 |
| PTPRS | 2.54 | 2.35 | 2.43 | 0.19 | 1.14 | 0.02 | 0.11 | 1.08 | 0.11 | -0.08 | -1.06 | 0.18 | 0.06 | 0.15 | 0.26 |
| TNC | 4.01 | 3.17 | 2.81 | 0.84 | 1.79 | 0.00 | 1.20 | 2.30 | 0.00 | 0.36 | 1.28 | 0.01 | 0.00 | 0.00 | 0.03 |
| TNF | 1.67 | 1.63 | 1.42 | 0.04 | 1.02 | 0.69 | 0.24 | 1.18 | 0.00 | 0.21 | 1.15 | 0.00 | 0.80 | 0.00 | 0.01 |
| TNFRSF10C | 3.07 | 2.79 | 2.69 | 0.28 | 1.22 | 0.02 | 0.39 | 1.31 | 0.00 | 0.10 | 1.07 | 0.25 | 0.06 | 0.00 | 0.34 |
| TNFSF13B | 4.98 | 5.01 | 4.77 | -0.03 | -1.02 | 0.85 | 0.21 | 1.16 | 0.10 | 0.24 | 1.18 | 0.02 | 0.90 | 0.13 | 0.06 |
| TNNI3 | 2.02 | 1.21 | 0.52 | 0.81 | 1.75 | 0.00 | 1.50 | 2.83 | 0.00 | 0.69 | 1.61 | 0.00 | 0.01 | 0.00 | 0.00 |
| TSHB | 1.65 | 1.46 | 1.53 | 0.18 | 1.13 | 0.21 | 0.11 | 1.08 | 0.35 | -0.07 | -1.05 | 0.52 | 0.35 | 0.42 | 0.62 |
| TSPAN1 | 0.82 | 0.52 | 0.26 | 0.29 | 1.23 | 0.06 | 0.56 | 1.47 | 0.00 | 0.26 | 1.20 | 0.02 | 0.15 | 0.00 | 0.05 |
| TYMP | 5.54 | 5.83 | 5.28 | -0.30 | -1.23 | 0.11 | 0.25 | 1.19 | 0.12 | 0.55 | 1.46 | 0.00 | 0.24 | 0.15 | 0.00 |
| TYRO3 | 2.35 | 2.26 | 2.36 | 0.09 | 1.06 | 0.23 | -0.01 | -1.01 | 0.85 | -0.10 | -1.07 | 0.06 | 0.38 | 0.88 | 0.11 |
| UMOD | 3.06 | 3.36 | 3.55 | -0.31 | -1.24 | 0.04 | -0.50 | -1.41 | 0.00 | -0.19 | -1.14 | 0.08 | 0.11 | 0.00 | 0.14 |
| USP8 | 1.48 | 1.28 | 1.34 | 0.20 | 1.14 | 0.37 | 0.13 | 1.10 | 0.48 | -0.06 | -1.04 | 0.70 | 0.52 | 0.55 | 0.77 |
| VAMP5 | 1.54 | 1.51 | 1.22 | 0.03 | 1.02 | 0.82 | 0.32 | 1.25 | 0.00 | 0.29 | 1.22 | 0.00 | 0.89 | 0.01 | 0.00 |
| VASN | 2.05 | 1.93 | 1.88 | 0.12 | 1.09 | 0.07 | 0.17 | 1.13 | 0.00 | 0.06 | 1.04 | 0.25 | 0.17 | 0.00 | 0.34 |
| VCAM1 | 4.19 | 3.76 | 3.60 | 0.43 | 1.35 | 0.00 | 0.59 | 1.51 | 0.00 | 0.16 | 1.11 | 0.06 | 0.00 | 0.00 | 0.11 |
| VIM | 1.61 | 1.52 | 1.13 | 0.09 | 1.07 | 0.56 | 0.48 | 1.40 | 0.00 | 0.39 | 1.31 | 0.00 | 0.68 | 0.00 | 0.00 |
| VSIR | 1.40 | 1.29 | 1.18 | 0.12 | 1.08 | 0.32 | 0.22 | 1.17 | 0.03 | 0.11 | 1.08 | 0.20 | 0.47 | 0.04 | 0.29 |
| VSTM2L | 0.20 | 0.13 | 0.12 | 0.07 | 1.05 | 0.54 | 0.08 | 1.06 | 0.41 | 0.01 | 1.01 | 0.89 | 0.67 | 0.48 | 0.91 |
| VWF | 6.05 | 5.97 | 5.53 | 0.08 | 1.05 | 0.51 | 0.52 | 1.43 | 0.00 | 0.44 | 1.36 | 0.00 | 0.65 | 0.00 | 0.00 |
| WASF1 | 0.75 | 0.66 | 0.74 | 0.09 | 1.07 | 0.49 | 0.01 | 1.01 | 0.94 | -0.08 | -1.06 | 0.39 | 0.63 | 0.95 | 0.49 |
| XG | 2.38 | 1.75 | 1.70 | 0.62 | 1.54 | 0.00 | 0.68 | 1.60 | 0.00 | 0.06 | 1.04 | 0.65 | 0.00 | 0.00 | 0.74 |
| ZBTB17 | 1.69 | 1.85 | 1.32 | -0.16 | -1.12 | 0.20 | 0.37 | 1.29 | 0.00 | 0.53 | 1.44 | 0.00 | 0.35 | 0.00 | 0.00 |
| ITGB1 | 2.25 | 2.16 | 2.14 | 0.09 | 1.06 | 0.21 | 0.10 | 1.07 | 0.10 | 0.01 | 1.01 | 0.79 | 0.36 | 0.13 | 0.85 |
| ITGB1BP2 | 2.64 | 2.27 | 2.46 | 0.36 | 1.29 | 0.20 | 0.17 | 1.13 | 0.47 | -0.19 | -1.14 | 0.35 | 0.34 | 0.55 | 0.44 |
| ITGB2 | 4.94 | 4.80 | 4.75 | 0.14 | 1.10 | 0.11 | 0.20 | 1.15 | 0.01 | 0.06 | 1.04 | 0.36 | 0.24 | 0.02 | 0.45 |
| ITIH3 | 2.39 | 2.30 | 2.07 | 0.09 | 1.06 | 0.48 | 0.32 | 1.25 | 0.00 | 0.23 | 1.18 | 0.01 | 0.62 | 0.01 | 0.03 |
| KIT | 1.25 | 1.44 | 1.54 | -0.19 | -1.14 | 0.02 | -0.29 | -1.22 | 0.00 | -0.10 | -1.07 | 0.09 | 0.06 | 0.00 | 0.15 |

| | | | | | | | | | | | | | | |
|---|---|---|---|---|---|---|---|---|---|---|---|---|---|---|
| KITLG | 2.68 | 2.31 | 2.55 | 0.37 | 1.29 | 0.01 | 0.14 | 1.10 | 0.28 | -0.23 | -1.18 | 0.03 | 0.04 | 0.34 | 0.06 |
| KYAT1 | 4.47 | 4.73 | 4.15 | -0.26 | -1.20 | 0.16 | 0.31 | 1.24 | 0.05 | 0.57 | 1.49 | 0.00 | 0.30 | 0.08 | 0.00 |
| LACTB2 | 1.33 | 1.28 | 0.88 | 0.05 | 1.04 | 0.66 | 0.45 | 1.36 | 0.00 | 0.39 | 1.31 | 0.00 | 0.78 | 0.00 | 0.00 |
| LBP | 5.19 | 5.32 | 4.80 | -0.12 | -1.09 | 0.40 | 0.39 | 1.31 | 0.00 | 0.51 | 1.43 | 0.00 | 0.54 | 0.00 | 0.00 |
| LDLR | 3.02 | 3.05 | 2.77 | -0.03 | -1.02 | 0.83 | 0.25 | 1.19 | 0.03 | 0.27 | 1.21 | 0.00 | 0.89 | 0.05 | 0.01 |
| LEP | 2.94 | 2.75 | 2.81 | 0.19 | 1.14 | 0.48 | 0.13 | 1.09 | 0.58 | -0.06 | -1.04 | 0.75 | 0.62 | 0.65 | 0.81 |
| LEPR | 0.49 | 0.33 | 0.29 | 0.16 | 1.11 | 0.02 | 0.20 | 1.15 | 0.00 | 0.04 | 1.03 | 0.41 | 0.07 | 0.00 | 0.51 |
| LGALS1 | 3.55 | 3.14 | 2.77 | 0.41 | 1.32 | 0.00 | 0.78 | 1.72 | 0.00 | 0.38 | 1.30 | 0.00 | 0.01 | 0.00 | 0.00 |
| LGALS3 | 4.14 | 4.14 | 3.77 | 0.00 | 1.00 | 0.98 | 0.37 | 1.30 | 0.00 | 0.37 | 1.29 | 0.00 | 0.99 | 0.00 | 0.00 |
| LILRA5 | 4.85 | 4.71 | 4.41 | 0.14 | 1.10 | 0.17 | 0.43 | 1.35 | 0.00 | 0.29 | 1.23 | 0.00 | 0.32 | 0.00 | 0.00 |
| LILRB1 | 4.15 | 4.20 | 4.07 | -0.06 | -1.04 | 0.64 | 0.07 | 1.05 | 0.49 | 0.13 | 1.09 | 0.14 | 0.76 | 0.56 | 0.21 |
| LILRB2 | 6.84 | 6.68 | 6.54 | 0.16 | 1.12 | 0.12 | 0.30 | 1.23 | 0.00 | 0.14 | 1.10 | 0.07 | 0.24 | 0.00 | 0.12 |
| LILRB5 | 5.88 | 6.02 | 6.00 | -0.13 | -1.10 | 0.45 | -0.12 | -1.09 | 0.44 | 0.02 | 1.01 | 0.90 | 0.58 | 0.51 | 0.91 |
| LPL | 5.15 | 4.42 | 4.58 | 0.73 | 1.66 | 0.00 | 0.57 | 1.48 | 0.00 | -0.16 | -1.12 | 0.14 | 0.00 | 0.00 | 0.21 |
| LRMP | 1.62 | 1.62 | 1.47 | 0.00 | 1.00 | 0.99 | 0.15 | 1.11 | 0.27 | 0.15 | 1.11 | 0.19 | 0.99 | 0.33 | 0.28 |
| LRP11 | 2.83 | 2.59 | 2.40 | 0.24 | 1.18 | 0.05 | 0.44 | 1.35 | 0.00 | 0.20 | 1.15 | 0.02 | 0.12 | 0.00 | 0.05 |
| LTBP2 | 5.83 | 5.15 | 4.39 | 0.68 | 1.60 | 0.00 | 1.44 | 2.71 | 0.00 | 0.76 | 1.70 | 0.00 | 0.00 | 0.00 | 0.00 |
| MARCO | 4.51 | 4.47 | 4.51 | 0.04 | 1.02 | 0.71 | 0.00 | -1.00 | 0.99 | -0.04 | -1.03 | 0.59 | 0.81 | 0.99 | 0.68 |
| MB | 6.62 | 5.48 | 4.66 | 1.14 | 2.21 | 0.00 | 1.96 | 3.89 | 0.00 | 0.82 | 1.76 | 0.00 | 0.00 | 0.00 | 0.00 |
| MCAM | 0.87 | 0.58 | 0.55 | 0.28 | 1.22 | 0.00 | 0.32 | 1.25 | 0.00 | 0.03 | 1.02 | 0.54 | 0.00 | 0.00 | 0.64 |
| MCFD2 | 3.49 | 2.98 | 2.53 | 0.50 | 1.42 | 0.00 | 0.96 | 1.95 | 0.00 | 0.46 | 1.37 | 0.00 | 0.00 | 0.00 | 0.00 |
| MEGF9 | 4.55 | 4.74 | 4.72 | -0.19 | -1.14 | 0.01 | -0.17 | -1.13 | 0.01 | 0.02 | 1.01 | 0.74 | 0.03 | 0.01 | 0.81 |
| MEP1B | 0.35 | 0.75 | 0.81 | -0.39 | -1.31 | 0.00 | -0.45 | -1.37 | 0.00 | -0.06 | -1.04 | 0.56 | 0.02 | 0.00 | 0.66 |
| MET | 4.95 | 5.07 | 4.93 | -0.12 | -1.09 | 0.12 | 0.02 | 1.01 | 0.75 | 0.14 | 1.10 | 0.01 | 0.24 | 0.80 | 0.03 |
| MFAP3 | 0.22 | 0.15 | 0.17 | 0.07 | 1.05 | 0.37 | 0.05 | 1.03 | 0.48 | -0.02 | -1.01 | 0.69 | 0.52 | 0.55 | 0.77 |
| MFAP5 | 3.26 | 2.71 | 2.73 | 0.54 | 1.46 | 0.00 | 0.52 | 1.44 | 0.00 | -0.02 | -1.01 | 0.86 | 0.00 | 0.00 | 0.90 |
| MMP7 | 1.57 | 1.26 | 1.04 | 0.31 | 1.24 | 0.02 | 0.53 | 1.44 | 0.00 | 0.22 | 1.17 | 0.02 | 0.07 | 0.00 | 0.05 |
| MNDA | 2.56 | 2.39 | 1.74 | 0.17 | 1.12 | 0.38 | 0.82 | 1.77 | 0.00 | 0.66 | 1.58 | 0.00 | 0.53 | 0.00 | 0.00 |
| MPHOSPH8 | 0.96 | 0.97 | 0.52 | -0.01 | -1.01 | 0.95 | 0.44 | 1.36 | 0.00 | 0.45 | 1.37 | 0.00 | 0.98 | 0.00 | 0.00 |
| MSMB | 0.67 | 0.25 | 0.06 | 0.43 | 1.34 | 0.00 | 0.61 | 1.53 | 0.00 | 0.19 | 1.14 | 0.08 | 0.02 | 0.00 | 0.15 |
| MSTN | 0.49 | 0.89 | 1.00 | -0.40 | -1.32 | 0.00 | -0.50 | -1.42 | 0.00 | -0.11 | -1.08 | 0.30 | 0.02 | 0.00 | 0.40 |
| MTPN | 0.15 | 0.14 | 0.07 | 0.01 | 1.01 | 0.88 | 0.08 | 1.06 | 0.08 | 0.07 | 1.05 | 0.06 | 0.93 | 0.11 | 0.11 |
| NADK | 5.19 | 5.75 | 4.56 | -0.56 | -1.47 | 0.01 | 0.64 | 1.56 | 0.00 | 1.20 | 2.29 | 0.00 | 0.03 | 0.00 | 0.00 |
| NCAM1 | 1.69 | 1.56 | 1.57 | 0.13 | 1.09 | 0.08 | 0.12 | 1.08 | 0.06 | -0.01 | -1.01 | 0.80 | 0.17 | 0.09 | 0.85 |

| | | | | | | | | | | | | | | |
|---|---|---|---|---|---|---|---|---|---|---|---|---|---|---|
| NECTIN2 | 6.48 | 6.05 | 5.77 | 0.42 | 1.34 | 0.01 | 0.71 | 1.63 | 0.00 | 0.29 | 1.22 | 0.01 | 0.02 | 0.00 | 0.03 |
| NID1 | 4.67 | 4.75 | 4.37 | -0.08 | -1.06 | 0.45 | 0.30 | 1.23 | 0.00 | 0.38 | 1.30 | 0.00 | 0.59 | 0.00 | 0.00 |
| NOTCH1 | 5.88 | 5.83 | 5.79 | 0.05 | 1.03 | 0.44 | 0.09 | 1.06 | 0.08 | 0.04 | 1.03 | 0.31 | 0.58 | 0.12 | 0.40 |
| NOTCH3 | 3.12 | 2.71 | 2.45 | 0.41 | 1.33 | 0.00 | 0.67 | 1.59 | 0.00 | 0.26 | 1.19 | 0.00 | 0.00 | 0.00 | 0.01 |
| NPDC1 | 3.13 | 2.31 | 2.02 | 0.81 | 1.76 | 0.00 | 1.10 | 2.15 | 0.00 | 0.29 | 1.22 | 0.03 | 0.00 | 0.00 | 0.07 |
| NPPB | 3.59 | 2.14 | 1.61 | 1.45 | 2.73 | 0.00 | 1.98 | 3.95 | 0.00 | 0.54 | 1.45 | 0.04 | 0.00 | 0.00 | 0.08 |
| NPTXR | 2.46 | 2.34 | 2.32 | 0.13 | 1.09 | 0.41 | 0.15 | 1.11 | 0.28 | 0.02 | 1.01 | 0.87 | 0.55 | 0.34 | 0.91 |
| NRCAM | 4.30 | 4.27 | 4.11 | 0.04 | 1.03 | 0.76 | 0.19 | 1.14 | 0.08 | 0.15 | 1.11 | 0.10 | 0.84 | 0.12 | 0.16 |
| NRP1 | 1.94 | 1.89 | 1.74 | 0.06 | 1.04 | 0.44 | 0.21 | 1.15 | 0.00 | 0.15 | 1.11 | 0.01 | 0.58 | 0.00 | 0.02 |
| NTproBNP | 3.37 | 1.88 | 1.15 | 1.49 | 2.80 | 0.00 | 2.22 | 4.64 | 0.00 | 0.73 | 1.66 | 0.01 | 0.00 | 0.00 | 0.02 |
| NTRK2 | 1.04 | 0.91 | 0.92 | 0.13 | 1.09 | 0.10 | 0.12 | 1.08 | 0.09 | -0.01 | -1.01 | 0.82 | 0.22 | 0.12 | 0.86 |
| OLR1 | 3.11 | 2.96 | 2.58 | 0.15 | 1.11 | 0.28 | 0.53 | 1.45 | 0.00 | 0.38 | 1.30 | 0.00 | 0.44 | 0.00 | 0.00 |
| OSMR | 0.80 | 0.71 | 0.62 | 0.09 | 1.06 | 0.07 | 0.17 | 1.13 | 0.00 | 0.09 | 1.06 | 0.01 | 0.17 | 0.00 | 0.03 |
| PAG1 | 4.01 | 3.92 | 3.25 | 0.09 | 1.06 | 0.55 | 0.75 | 1.69 | 0.00 | 0.66 | 1.58 | 0.00 | 0.68 | 0.00 | 0.00 |
| PAM | 4.38 | 4.22 | 4.13 | 0.17 | 1.12 | 0.07 | 0.25 | 1.19 | 0.00 | 0.08 | 1.06 | 0.20 | 0.16 | 0.00 | 0.30 |
| PCDH17 | 1.79 | 1.53 | 1.40 | 0.26 | 1.19 | 0.01 | 0.39 | 1.31 | 0.00 | 0.13 | 1.09 | 0.08 | 0.04 | 0.00 | 0.14 |
| PCOLCE | 4.43 | 4.26 | 4.36 | 0.17 | 1.13 | 0.22 | 0.07 | 1.05 | 0.56 | -0.10 | -1.07 | 0.32 | 0.38 | 0.63 | 0.41 |
| PCSK9 | 3.04 | 3.05 | 2.63 | -0.01 | -1.01 | 0.92 | 0.41 | 1.33 | 0.00 | 0.42 | 1.34 | 0.00 | 0.96 | 0.00 | 0.00 |
| PDCD6 | 2.48 | 2.47 | 2.35 | 0.02 | 1.01 | 0.89 | 0.14 | 1.10 | 0.16 | 0.12 | 1.09 | 0.14 | 0.94 | 0.21 | 0.21 |
| PDGFA | 1.58 | 2.06 | 1.93 | -0.48 | -1.39 | 0.01 | -0.35 | -1.27 | 0.03 | 0.13 | 1.09 | 0.34 | 0.03 | 0.04 | 0.44 |
| PDGFRA | 2.36 | 1.81 | 1.58 | 0.55 | 1.46 | 0.00 | 0.78 | 1.72 | 0.00 | 0.23 | 1.18 | 0.00 | 0.00 | 0.00 | 0.01 |
| PDGFRB | 2.69 | 2.53 | 2.57 | 0.17 | 1.12 | 0.05 | 0.12 | 1.09 | 0.10 | -0.04 | -1.03 | 0.46 | 0.13 | 0.13 | 0.56 |
| PEAR1 | 1.10 | 1.01 | 1.03 | 0.09 | 1.06 | 0.07 | 0.07 | 1.05 | 0.11 | -0.02 | -1.01 | 0.58 | 0.17 | 0.14 | 0.67 |
| PGLYRP1 | 5.82 | 5.27 | 5.00 | 0.55 | 1.46 | 0.00 | 0.82 | 1.76 | 0.00 | 0.27 | 1.21 | 0.02 | 0.00 | 0.00 | 0.05 |
| PILRB | 3.18 | 2.53 | 2.34 | 0.66 | 1.58 | 0.00 | 0.84 | 1.80 | 0.00 | 0.19 | 1.14 | 0.15 | 0.00 | 0.00 | 0.23 |
| PLA2G1B | 4.76 | 4.48 | 4.21 | 0.28 | 1.22 | 0.08 | 0.56 | 1.47 | 0.00 | 0.27 | 1.21 | 0.02 | 0.18 | 0.00 | 0.05 |
| PLA2G2A | 4.65 | 4.47 | 3.20 | 0.19 | 1.14 | 0.53 | 1.45 | 2.73 | 0.00 | 1.26 | 2.40 | 0.00 | 0.66 | 0.00 | 0.00 |
| PLAT | 5.16 | 4.70 | 4.20 | 0.46 | 1.37 | 0.01 | 0.97 | 1.95 | 0.00 | 0.51 | 1.42 | 0.00 | 0.02 | 0.00 | 0.00 |
| PLIN3 | 2.26 | 2.08 | 1.59 | 0.18 | 1.13 | 0.13 | 0.67 | 1.59 | 0.00 | 0.49 | 1.40 | 0.00 | 0.25 | 0.00 | 0.00 |
| PLPBP | 2.74 | 2.76 | 2.38 | -0.01 | -1.01 | 0.95 | 0.37 | 1.29 | 0.04 | 0.38 | 1.30 | 0.01 | 0.98 | 0.07 | 0.03 |
| PLTP | 2.91 | 2.72 | 2.71 | 0.19 | 1.14 | 0.14 | 0.19 | 1.14 | 0.08 | 0.00 | 1.00 | 0.96 | 0.27 | 0.11 | 0.97 |
| PLXNB2 | 5.74 | 5.64 | 5.35 | 0.10 | 1.07 | 0.32 | 0.40 | 1.32 | 0.00 | 0.30 | 1.23 | 0.00 | 0.47 | 0.00 | 0.00 |
| PLXNB3 | 0.93 | 0.86 | 0.82 | 0.06 | 1.05 | 0.38 | 0.11 | 1.08 | 0.09 | 0.04 | 1.03 | 0.41 | 0.52 | 0.12 | 0.51 |
| PON2 | -0.09 | -0.02 | -0.08 | -0.08 | -1.05 | 0.30 | -0.01 | -1.01 | 0.82 | 0.06 | 1.04 | 0.24 | 0.45 | 0.86 | 0.34 |

| | | | | | | | | | | | | | | | |
|---|---|---|---|---|---|---|---|---|---|---|---|---|---|---|---|
| PPIB | 0.82 | 0.93 | 0.72 | -0.11 | -1.08 | 0.50 | 0.09 | 1.07 | 0.51 | 0.20 | 1.15 | 0.08 | 0.63 | 0.58 | 0.15 |
| PPP1R2 | 1.78 | 1.64 | 1.20 | 0.14 | 1.10 | 0.29 | 0.58 | 1.50 | 0.00 | 0.44 | 1.35 | 0.00 | 0.44 | 0.00 | 0.00 |
| PRCP | 2.63 | 2.95 | 2.84 | -0.33 | -1.25 | 0.00 | -0.21 | -1.16 | 0.02 | 0.11 | 1.08 | 0.12 | 0.01 | 0.03 | 0.19 |
| PRKAR1A | 3.49 | 3.17 | 3.06 | 0.32 | 1.25 | 0.15 | 0.43 | 1.35 | 0.03 | 0.11 | 1.08 | 0.50 | 0.29 | 0.04 | 0.60 |
| PROC | 2.89 | 3.14 | 3.31 | -0.24 | -1.19 | 0.00 | -0.42 | -1.34 | 0.00 | -0.17 | -1.13 | 0.00 | 0.01 | 0.00 | 0.01 |
| PRSS2 | 3.39 | 2.78 | 2.55 | 0.62 | 1.53 | 0.00 | 0.84 | 1.79 | 0.00 | 0.23 | 1.17 | 0.12 | 0.01 | 0.00 | 0.20 |
| PRSS27 | 2.81 | 2.71 | 2.71 | 0.10 | 1.07 | 0.41 | 0.10 | 1.07 | 0.33 | 0.00 | 1.00 | 0.98 | 0.55 | 0.40 | 0.98 |
| PRTN3 | 3.57 | 3.43 | 3.00 | 0.13 | 1.10 | 0.34 | 0.56 | 1.48 | 0.00 | 0.43 | 1.35 | 0.00 | 0.49 | 0.00 | 0.00 |
| ACAN | 2.81 | 2.53 | 2.37 | 0.28 | 1.21 | 0.01 | 0.44 | 1.35 | 0.00 | 0.16 | 1.12 | 0.04 | 0.03 | 0.00 | 0.08 |
| ACE2 | 1.70 | 1.82 | 1.25 | -0.12 | -1.09 | 0.49 | 0.45 | 1.36 | 0.00 | 0.57 | 1.48 | 0.00 | 0.63 | 0.01 | 0.00 |
| ACOX1 | 0.16 | 0.17 | 0.12 | -0.01 | -1.01 | 0.86 | 0.04 | 1.03 | 0.53 | 0.05 | 1.04 | 0.32 | 0.91 | 0.60 | 0.41 |
| ACP5 | 4.33 | 4.53 | 4.20 | -0.21 | -1.15 | 0.09 | 0.13 | 1.09 | 0.22 | 0.33 | 1.26 | 0.00 | 0.20 | 0.27 | 0.00 |
| ACTA2 | 0.87 | 0.22 | -0.13 | 0.65 | 1.57 | 0.00 | 1.00 | 2.00 | 0.00 | 0.36 | 1.28 | 0.00 | 0.00 | 0.00 | 0.01 |
| ACY1 | 3.70 | 4.35 | 3.69 | -0.65 | -1.56 | 0.00 | 0.01 | 1.01 | 0.94 | 0.66 | 1.58 | 0.00 | 0.01 | 0.95 | 0.00 |
| ADA2 | 5.37 | 5.13 | 5.17 | 0.25 | 1.19 | 0.05 | 0.21 | 1.15 | 0.06 | -0.04 | -1.03 | 0.66 | 0.13 | 0.09 | 0.74 |
| ADAM15 | 3.49 | 3.46 | 3.40 | 0.03 | 1.02 | 0.72 | 0.09 | 1.06 | 0.24 | 0.06 | 1.04 | 0.36 | 0.82 | 0.30 | 0.46 |
| ADAMTS13 | 3.84 | 4.08 | 4.20 | -0.24 | -1.18 | 0.00 | -0.36 | -1.28 | 0.00 | -0.12 | -1.09 | 0.03 | 0.01 | 0.00 | 0.07 |
| ADAMTS16 | 0.58 | 0.41 | 0.31 | 0.18 | 1.13 | 0.01 | 0.27 | 1.21 | 0.00 | 0.09 | 1.07 | 0.07 | 0.04 | 0.00 | 0.12 |
| ADGRG2 | 2.87 | 2.89 | 2.96 | -0.02 | -1.01 | 0.83 | -0.09 | -1.07 | 0.27 | -0.07 | -1.05 | 0.31 | 0.89 | 0.33 | 0.40 |
| ADH4 | 2.30 | 2.96 | 2.66 | -0.67 | -1.59 | 0.01 | -0.36 | -1.29 | 0.10 | 0.31 | 1.24 | 0.10 | 0.03 | 0.13 | 0.17 |
| AGXT | 1.54 | 2.30 | 1.65 | -0.76 | -1.70 | 0.00 | -0.11 | -1.08 | 0.57 | 0.65 | 1.57 | 0.00 | 0.01 | 0.64 | 0.00 |
| AHCY | 2.01 | 2.07 | 1.61 | -0.06 | -1.04 | 0.62 | 0.39 | 1.31 | 0.00 | 0.46 | 1.37 | 0.00 | 0.75 | 0.00 | 0.00 |
| AK1 | 2.74 | 2.73 | 2.20 | 0.01 | 1.01 | 0.97 | 0.53 | 1.45 | 0.02 | 0.53 | 1.44 | 0.01 | 0.99 | 0.03 | 0.02 |
| AKR1C4 | 0.48 | 0.65 | 0.45 | -0.16 | -1.12 | 0.11 | 0.03 | 1.02 | 0.72 | 0.20 | 1.15 | 0.01 | 0.23 | 0.78 | 0.02 |
| ALCAM | 1.90 | 1.82 | 1.71 | 0.08 | 1.06 | 0.28 | 0.19 | 1.14 | 0.01 | 0.10 | 1.07 | 0.07 | 0.44 | 0.01 | 0.13 |
| AMY2A | 1.86 | 1.63 | 1.61 | 0.24 | 1.18 | 0.13 | 0.25 | 1.19 | 0.07 | 0.01 | 1.01 | 0.90 | 0.27 | 0.10 | 0.91 |
| AMY2B | 1.74 | 1.54 | 1.52 | 0.20 | 1.15 | 0.20 | 0.22 | 1.16 | 0.10 | 0.02 | 1.01 | 0.87 | 0.35 | 0.14 | 0.90 |
| ANG | 7.04 | 7.00 | 7.06 | 0.04 | 1.03 | 0.72 | -0.01 | -1.01 | 0.88 | -0.05 | -1.04 | 0.50 | 0.81 | 0.91 | 0.60 |
| ANGPTL1 | 2.69 | 2.56 | 2.08 | 0.13 | 1.10 | 0.29 | 0.61 | 1.53 | 0.00 | 0.48 | 1.39 | 0.00 | 0.45 | 0.00 | 0.00 |
| ANGPTL3 | 4.43 | 4.31 | 4.30 | 0.11 | 1.08 | 0.29 | 0.12 | 1.09 | 0.18 | 0.01 | 1.01 | 0.89 | 0.44 | 0.23 | 0.91 |
| ANPEP | 1.29 | 1.35 | 1.30 | -0.06 | -1.04 | 0.51 | -0.01 | -1.00 | 0.93 | 0.05 | 1.04 | 0.41 | 0.64 | 0.95 | 0.51 |
| ANXA4 | 0.43 | 0.46 | 0.38 | -0.03 | -1.02 | 0.69 | 0.06 | 1.04 | 0.33 | 0.08 | 1.06 | 0.09 | 0.80 | 0.40 | 0.15 |
| AOC3 | 1.70 | 1.57 | 1.53 | 0.13 | 1.09 | 0.15 | 0.17 | 1.13 | 0.02 | 0.05 | 1.03 | 0.46 | 0.29 | 0.04 | 0.56 |
| APLP1 | 1.24 | 1.13 | 1.45 | 0.11 | 1.08 | 0.31 | -0.21 | -1.16 | 0.03 | -0.32 | -1.25 | 0.00 | 0.46 | 0.04 | 0.00 |

| | | | | | | | | | | | | | | |
|---|---|---|---|---|---|---|---|---|---|---|---|---|---|---|
| APOM | 2.92 | 3.04 | 3.33 | -0.12 | -1.09 | 0.25 | -0.41 | -1.33 | 0.00 | -0.29 | -1.22 | 0.00 | 0.39 | 0.00 | 0.00 |
| ART3 | 2.17 | 1.80 | 1.66 | 0.37 | 1.29 | 0.00 | 0.51 | 1.42 | 0.00 | 0.14 | 1.10 | 0.12 | 0.02 | 0.00 | 0.20 |
| AXL | 2.92 | 2.76 | 2.75 | 0.15 | 1.11 | 0.11 | 0.17 | 1.12 | 0.05 | 0.01 | 1.01 | 0.84 | 0.24 | 0.07 | 0.89 |
| AZU1 | 2.69 | 2.50 | 2.02 | 0.18 | 1.14 | 0.39 | 0.67 | 1.59 | 0.00 | 0.49 | 1.40 | 0.00 | 0.54 | 0.00 | 0.01 |
| BAG6 | 4.48 | 4.48 | 4.11 | -0.01 | -1.00 | 0.96 | 0.36 | 1.29 | 0.00 | 0.37 | 1.29 | 0.00 | 0.99 | 0.01 | 0.00 |
| BLMH | 2.86 | 3.11 | 2.52 | -0.25 | -1.19 | 0.06 | 0.34 | 1.26 | 0.00 | 0.59 | 1.50 | 0.00 | 0.14 | 0.01 | 0.00 |
| BMP6 | 2.43 | 2.32 | 2.20 | 0.12 | 1.09 | 0.15 | 0.24 | 1.18 | 0.00 | 0.12 | 1.09 | 0.04 | 0.29 | 0.00 | 0.09 |
| BOC | 1.70 | 1.46 | 1.51 | 0.24 | 1.18 | 0.03 | 0.19 | 1.14 | 0.04 | -0.04 | -1.03 | 0.57 | 0.08 | 0.06 | 0.67 |
| BPIFB1 | 5.99 | 5.60 | 5.69 | 0.38 | 1.31 | 0.03 | 0.30 | 1.23 | 0.05 | -0.08 | -1.06 | 0.51 | 0.08 | 0.07 | 0.61 |
| C1QTNF1 | 3.69 | 3.72 | 3.30 | -0.03 | -1.02 | 0.78 | 0.39 | 1.31 | 0.00 | 0.42 | 1.34 | 0.00 | 0.86 | 0.00 | 0.00 |
| C2 | 4.93 | 4.99 | 4.91 | -0.06 | -1.04 | 0.57 | 0.02 | 1.02 | 0.80 | 0.08 | 1.06 | 0.27 | 0.69 | 0.84 | 0.37 |
| CA1 | 8.09 | 8.08 | 7.65 | 0.01 | 1.01 | 0.98 | 0.44 | 1.35 | 0.06 | 0.43 | 1.35 | 0.02 | 0.99 | 0.08 | 0.06 |
| CA13 | 1.81 | 1.50 | 1.65 | 0.30 | 1.23 | 0.19 | 0.16 | 1.12 | 0.42 | -0.14 | -1.10 | 0.40 | 0.34 | 0.48 | 0.49 |
| CA3 | 7.31 | 6.58 | 5.85 | 0.73 | 1.66 | 0.01 | 1.46 | 2.75 | 0.00 | 0.73 | 1.65 | 0.00 | 0.04 | 0.00 | 0.00 |
| CA4 | 2.34 | 2.31 | 2.14 | 0.03 | 1.02 | 0.71 | 0.21 | 1.16 | 0.00 | 0.18 | 1.13 | 0.00 | 0.81 | 0.01 | 0.01 |
| CA5A | 2.64 | 2.94 | 2.18 | -0.30 | -1.23 | 0.30 | 0.45 | 1.37 | 0.07 | 0.75 | 1.68 | 0.00 | 0.45 | 0.10 | 0.00 |
| CANT1 | 3.52 | 3.49 | 3.41 | 0.02 | 1.02 | 0.74 | 0.11 | 1.08 | 0.10 | 0.08 | 1.06 | 0.13 | 0.83 | 0.13 | 0.20 |
| CASP3 | 3.08 | 3.09 | 2.83 | -0.01 | -1.01 | 0.95 | 0.24 | 1.18 | 0.21 | 0.26 | 1.20 | 0.11 | 0.98 | 0.26 | 0.19 |
| CBLIF | 1.98 | 1.41 | 1.65 | 0.57 | 1.48 | 0.01 | 0.33 | 1.26 | 0.06 | -0.24 | -1.18 | 0.11 | 0.02 | 0.09 | 0.18 |
| CCDC80 | 4.92 | 4.48 | 4.01 | 0.44 | 1.35 | 0.01 | 0.92 | 1.89 | 0.00 | 0.48 | 1.39 | 0.00 | 0.02 | 0.00 | 0.00 |
| CCL14 | 6.83 | 6.47 | 6.33 | 0.36 | 1.28 | 0.01 | 0.49 | 1.41 | 0.00 | 0.14 | 1.10 | 0.16 | 0.03 | 0.00 | 0.24 |
| CCL15 | 6.93 | 6.68 | 6.45 | 0.25 | 1.19 | 0.14 | 0.48 | 1.40 | 0.00 | 0.24 | 1.18 | 0.05 | 0.27 | 0.00 | 0.09 |
| CCL16 | 4.89 | 4.58 | 4.92 | 0.31 | 1.24 | 0.18 | -0.02 | -1.02 | 0.91 | -0.33 | -1.26 | 0.04 | 0.32 | 0.93 | 0.09 |
| CCL27 | 3.55 | 3.06 | 2.94 | 0.50 | 1.41 | 0.00 | 0.62 | 1.54 | 0.00 | 0.12 | 1.09 | 0.26 | 0.01 | 0.00 | 0.36 |
| CCN3 | 3.54 | 2.62 | 2.41 | 0.92 | 1.90 | 0.00 | 1.13 | 2.19 | 0.00 | 0.21 | 1.15 | 0.16 | 0.00 | 0.00 | 0.24 |
| CD14 | 7.54 | 7.41 | 7.20 | 0.13 | 1.10 | 0.38 | 0.34 | 1.26 | 0.01 | 0.21 | 1.15 | 0.06 | 0.53 | 0.02 | 0.11 |
| CD163 | 3.33 | 3.24 | 3.06 | 0.09 | 1.06 | 0.44 | 0.26 | 1.20 | 0.01 | 0.17 | 1.13 | 0.03 | 0.58 | 0.01 | 0.07 |
| CD209 | 4.03 | 3.85 | 3.70 | 0.18 | 1.13 | 0.06 | 0.32 | 1.25 | 0.00 | 0.14 | 1.10 | 0.04 | 0.16 | 0.00 | 0.08 |
| CD2AP | 4.58 | 4.32 | 4.22 | 0.26 | 1.20 | 0.29 | 0.36 | 1.28 | 0.10 | 0.10 | 1.07 | 0.58 | 0.45 | 0.13 | 0.67 |
| CD46 | 4.25 | 3.92 | 3.80 | 0.33 | 1.26 | 0.00 | 0.44 | 1.36 | 0.00 | 0.11 | 1.08 | 0.16 | 0.01 | 0.00 | 0.24 |
| CD55 | 3.67 | 3.43 | 3.18 | 0.24 | 1.18 | 0.04 | 0.49 | 1.40 | 0.00 | 0.25 | 1.19 | 0.00 | 0.10 | 0.00 | 0.01 |
| CD59 | 4.71 | 4.16 | 3.84 | 0.56 | 1.47 | 0.00 | 0.87 | 1.83 | 0.00 | 0.32 | 1.25 | 0.01 | 0.00 | 0.00 | 0.02 |
| CD69 | 4.95 | 4.59 | 4.83 | 0.35 | 1.28 | 0.24 | 0.11 | 1.08 | 0.67 | -0.24 | -1.18 | 0.26 | 0.39 | 0.73 | 0.36 |
| CD93 | 4.38 | 3.97 | 3.81 | 0.41 | 1.33 | 0.00 | 0.56 | 1.48 | 0.00 | 0.15 | 1.11 | 0.08 | 0.00 | 0.00 | 0.14 |

| | | | | | | | | | | | | | | | |
|---|---|---|---|---|---|---|---|---|---|---|---|---|---|---|---|
| CD97 | 4.62 | 4.39 | 4.22 | 0.23 | 1.17 | 0.04 | 0.41 | 1.33 | 0.00 | 0.18 | 1.13 | 0.03 | 0.11 | 0.00 | 0.07 |
| CDH1 | 2.66 | 2.43 | 2.24 | 0.23 | 1.17 | 0.03 | 0.42 | 1.34 | 0.00 | 0.19 | 1.14 | 0.01 | 0.08 | 0.00 | 0.03 |
| CDH17 | 0.38 | 0.11 | 0.16 | 0.27 | 1.21 | 0.02 | 0.22 | 1.16 | 0.02 | -0.05 | -1.04 | 0.53 | 0.05 | 0.04 | 0.62 |
| CDH2 | 2.44 | 2.43 | 2.14 | 0.00 | 1.00 | 0.98 | 0.30 | 1.23 | 0.00 | 0.29 | 1.22 | 0.00 | 0.99 | 0.00 | 0.00 |
| CDH5 | 1.80 | 1.58 | 1.69 | 0.23 | 1.17 | 0.00 | 0.12 | 1.08 | 0.07 | -0.11 | -1.08 | 0.05 | 0.01 | 0.11 | 0.09 |
| CDH6 | 1.23 | 0.89 | 0.89 | 0.33 | 1.26 | 0.00 | 0.34 | 1.26 | 0.00 | 0.00 | 1.00 | 0.97 | 0.01 | 0.00 | 0.97 |
| CDHR5 | 2.44 | 2.56 | 2.74 | -0.12 | -1.08 | 0.31 | -0.30 | -1.23 | 0.00 | -0.18 | -1.13 | 0.03 | 0.45 | 0.01 | 0.06 |
| CEACAM8 | 3.36 | 2.70 | 2.19 | 0.65 | 1.57 | 0.00 | 1.17 | 2.25 | 0.00 | 0.52 | 1.43 | 0.00 | 0.01 | 0.00 | 0.00 |
| CEBPB | 0.55 | 0.64 | 0.33 | -0.10 | -1.07 | 0.29 | 0.22 | 1.16 | 0.01 | 0.31 | 1.24 | 0.00 | 0.45 | 0.01 | 0.00 |
| CELA3A | 3.42 | 3.49 | 3.24 | -0.07 | -1.05 | 0.67 | 0.18 | 1.14 | 0.20 | 0.25 | 1.19 | 0.03 | 0.79 | 0.25 | 0.07 |
| CES1 | 4.65 | 5.34 | 4.76 | -0.69 | -1.61 | 0.00 | -0.11 | -1.08 | 0.58 | 0.58 | 1.49 | 0.00 | 0.01 | 0.64 | 0.00 |
| CGREF1 | 2.93 | 2.38 | 2.08 | 0.56 | 1.47 | 0.00 | 0.85 | 1.81 | 0.00 | 0.30 | 1.23 | 0.01 | 0.00 | 0.00 | 0.03 |
| CHI3L1 | 5.15 | 4.13 | 2.91 | 1.01 | 2.02 | 0.00 | 2.24 | 4.72 | 0.00 | 1.23 | 2.34 | 0.00 | 0.00 | 0.00 | 0.00 |
| CHIT1 | 5.76 | 4.78 | 4.50 | 0.98 | 1.97 | 0.01 | 1.26 | 2.39 | 0.00 | 0.28 | 1.21 | 0.29 | 0.03 | 0.00 | 0.39 |
| CHL1 | 2.36 | 2.01 | 2.10 | 0.36 | 1.28 | 0.00 | 0.26 | 1.20 | 0.00 | -0.10 | -1.07 | 0.16 | 0.00 | 0.00 | 0.24 |
| CHRDL2 | 2.70 | 2.37 | 2.12 | 0.32 | 1.25 | 0.03 | 0.58 | 1.49 | 0.00 | 0.26 | 1.19 | 0.02 | 0.09 | 0.00 | 0.05 |
| CLC | 1.13 | 0.60 | 0.63 | 0.53 | 1.45 | 0.00 | 0.50 | 1.42 | 0.00 | -0.03 | -1.02 | 0.70 | 0.00 | 0.00 | 0.78 |
| CLEC1A | 2.08 | 1.83 | 1.64 | 0.26 | 1.19 | 0.03 | 0.44 | 1.36 | 0.00 | 0.18 | 1.14 | 0.03 | 0.08 | 0.00 | 0.06 |
| CLEC5A | 3.20 | 3.03 | 2.43 | 0.17 | 1.12 | 0.23 | 0.77 | 1.71 | 0.00 | 0.60 | 1.52 | 0.00 | 0.38 | 0.00 | 0.00 |
| CLTA | 1.00 | 0.94 | 0.53 | 0.05 | 1.04 | 0.55 | 0.47 | 1.38 | 0.00 | 0.41 | 1.33 | 0.00 | 0.68 | 0.00 | 0.00 |
| CLUL1 | 2.38 | 2.03 | 2.23 | 0.35 | 1.27 | 0.00 | 0.14 | 1.11 | 0.10 | -0.21 | -1.15 | 0.01 | 0.00 | 0.13 | 0.02 |
| CNDP1 | 0.82 | 0.98 | 1.05 | -0.16 | -1.12 | 0.12 | -0.23 | -1.18 | 0.01 | -0.07 | -1.05 | 0.33 | 0.24 | 0.02 | 0.42 |
| CNPY2 | 0.80 | 0.79 | 0.67 | 0.01 | 1.01 | 0.90 | 0.13 | 1.10 | 0.12 | 0.12 | 1.09 | 0.09 | 0.94 | 0.16 | 0.16 |
| CNST | 2.41 | 1.90 | 2.50 | 0.51 | 1.43 | 0.13 | -0.09 | -1.06 | 0.77 | -0.60 | -1.51 | 0.01 | 0.25 | 0.82 | 0.03 |
| CNTN1 | 1.65 | 1.52 | 1.57 | 0.13 | 1.10 | 0.05 | 0.08 | 1.06 | 0.17 | -0.05 | -1.04 | 0.30 | 0.13 | 0.21 | 0.40 |
| COL18A1 | 6.43 | 6.13 | 5.98 | 0.30 | 1.23 | 0.03 | 0.45 | 1.37 | 0.00 | 0.16 | 1.11 | 0.11 | 0.08 | 0.00 | 0.18 |
| COL1A1 | 0.51 | 0.38 | 0.40 | 0.13 | 1.10 | 0.03 | 0.12 | 1.08 | 0.03 | -0.02 | -1.01 | 0.71 | 0.09 | 0.05 | 0.78 |
| COL4A1 | 4.15 | 3.89 | 3.68 | 0.26 | 1.20 | 0.03 | 0.47 | 1.38 | 0.00 | 0.21 | 1.15 | 0.01 | 0.08 | 0.00 | 0.04 |
| COL6A3 | 6.32 | 5.44 | 5.11 | 0.88 | 1.85 | 0.00 | 1.22 | 2.33 | 0.00 | 0.33 | 1.26 | 0.04 | 0.00 | 0.00 | 0.08 |
| COMP | 5.45 | 4.83 | 4.85 | 0.62 | 1.53 | 0.00 | 0.60 | 1.52 | 0.00 | -0.02 | -1.01 | 0.86 | 0.00 | 0.00 | 0.90 |
| COMT | 3.68 | 3.43 | 3.06 | 0.24 | 1.18 | 0.30 | 0.62 | 1.54 | 0.00 | 0.38 | 1.30 | 0.03 | 0.45 | 0.00 | 0.06 |
| CORO1A | 1.43 | 1.26 | 1.11 | 0.17 | 1.13 | 0.31 | 0.32 | 1.25 | 0.03 | 0.15 | 1.11 | 0.21 | 0.45 | 0.04 | 0.30 |
| CPA1 | 5.97 | 5.59 | 5.65 | 0.38 | 1.31 | 0.03 | 0.33 | 1.25 | 0.04 | -0.06 | -1.04 | 0.66 | 0.09 | 0.06 | 0.74 |
| CPB1 | 2.39 | 1.86 | 1.90 | 0.53 | 1.44 | 0.00 | 0.49 | 1.41 | 0.00 | -0.04 | -1.03 | 0.76 | 0.01 | 0.00 | 0.82 |

| | | | | | | | | | | | | | | | |
|---|---|---|---|---|---|---|---|---|---|---|---|---|---|---|---|
| CR2 | 2.85 | 2.74 | 3.06 | 0.10 | 1.08 | 0.42 | -0.21 | -1.16 | 0.06 | -0.31 | -1.24 | 0.00 | 0.56 | 0.09 | 0.00 |
| CRTAC1 | 1.61 | 1.35 | 1.56 | 0.26 | 1.20 | 0.07 | 0.04 | 1.03 | 0.72 | -0.22 | -1.16 | 0.03 | 0.16 | 0.78 | 0.08 |
| CRX | -0.10 | 0.03 | 0.06 | -0.13 | -1.10 | 0.34 | -0.16 | -1.12 | 0.17 | -0.03 | -1.02 | 0.74 | 0.48 | 0.21 | 0.81 |
| CST3 | 7.74 | 6.96 | 6.74 | 0.78 | 1.72 | 0.00 | 1.00 | 2.00 | 0.00 | 0.22 | 1.16 | 0.11 | 0.00 | 0.00 | 0.18 |
| CST6 | 3.65 | 3.24 | 3.04 | 0.41 | 1.33 | 0.03 | 0.61 | 1.53 | 0.00 | 0.20 | 1.15 | 0.12 | 0.08 | 0.00 | 0.20 |
| CSTB | 3.55 | 3.05 | 2.44 | 0.50 | 1.42 | 0.01 | 1.12 | 2.17 | 0.00 | 0.61 | 1.53 | 0.00 | 0.03 | 0.00 | 0.00 |
| CTF1 | 0.13 | 0.27 | 0.06 | -0.14 | -1.11 | 0.33 | 0.06 | 1.04 | 0.63 | 0.21 | 1.15 | 0.06 | 0.48 | 0.70 | 0.11 |
| CTSB | 3.28 | 3.02 | 2.57 | 0.26 | 1.20 | 0.07 | 0.70 | 1.63 | 0.00 | 0.44 | 1.36 | 0.00 | 0.16 | 0.00 | 0.00 |
| CTSD | 2.30 | 2.26 | 2.02 | 0.04 | 1.03 | 0.72 | 0.28 | 1.22 | 0.00 | 0.24 | 1.18 | 0.00 | 0.81 | 0.01 | 0.01 |
| CTSH | 3.42 | 3.64 | 3.37 | -0.22 | -1.17 | 0.17 | 0.05 | 1.04 | 0.72 | 0.27 | 1.21 | 0.02 | 0.32 | 0.78 | 0.05 |
| CTSL | 5.20 | 5.06 | 4.49 | 0.14 | 1.10 | 0.29 | 0.71 | 1.64 | 0.00 | 0.57 | 1.48 | 0.00 | 0.45 | 0.00 | 0.00 |
| CTSZ | 1.76 | 1.68 | 1.45 | 0.08 | 1.06 | 0.40 | 0.31 | 1.24 | 0.00 | 0.23 | 1.17 | 0.00 | 0.54 | 0.00 | 0.01 |
| CXCL16 | 6.45 | 6.26 | 6.06 | 0.19 | 1.14 | 0.08 | 0.39 | 1.31 | 0.00 | 0.20 | 1.15 | 0.01 | 0.18 | 0.00 | 0.03 |
| CXCL5 | 4.09 | 4.22 | 4.53 | -0.13 | -1.09 | 0.63 | -0.44 | -1.36 | 0.05 | -0.31 | -1.24 | 0.10 | 0.75 | 0.08 | 0.17 |
| CXCL8 | 3.95 | 3.62 | 2.95 | 0.33 | 1.26 | 0.09 | 1.00 | 2.00 | 0.00 | 0.67 | 1.59 | 0.00 | 0.20 | 0.00 | 0.00 |
| DCN | 3.47 | 2.65 | 2.32 | 0.82 | 1.77 | 0.00 | 1.16 | 2.23 | 0.00 | 0.33 | 1.26 | 0.00 | 0.00 | 0.00 | 0.01 |
| DCTPP1 | 3.31 | 3.54 | 2.84 | -0.23 | -1.17 | 0.11 | 0.47 | 1.39 | 0.00 | 0.70 | 1.62 | 0.00 | 0.24 | 0.00 | 0.00 |
| DDC | 4.14 | 4.43 | 4.20 | -0.29 | -1.23 | 0.11 | -0.07 | -1.05 | 0.67 | 0.22 | 1.17 | 0.09 | 0.24 | 0.73 | 0.16 |
| DEFA1 | 1.25 | 0.81 | 0.51 | 0.45 | 1.36 | 0.01 | 0.75 | 1.68 | 0.00 | 0.30 | 1.23 | 0.02 | 0.03 | 0.00 | 0.04 |
| DIABLO | 1.63 | 1.36 | 1.36 | 0.26 | 1.20 | 0.18 | 0.27 | 1.20 | 0.12 | 0.00 | 1.00 | 0.97 | 0.33 | 0.15 | 0.98 |
| DKK3 | 5.07 | 4.42 | 4.25 | 0.66 | 1.58 | 0.00 | 0.82 | 1.76 | 0.00 | 0.16 | 1.12 | 0.10 | 0.00 | 0.00 | 0.16 |
| DLK1 | 4.73 | 4.01 | 3.84 | 0.72 | 1.65 | 0.00 | 0.89 | 1.85 | 0.00 | 0.16 | 1.12 | 0.24 | 0.00 | 0.00 | 0.34 |
| DNAJB8 | 0.25 | 0.20 | 0.23 | 0.05 | 1.03 | 0.44 | 0.03 | 1.02 | 0.63 | -0.02 | -1.02 | 0.61 | 0.58 | 0.70 | 0.70 |
| DOK2 | 1.82 | 1.54 | 1.68 | 0.28 | 1.21 | 0.25 | 0.14 | 1.10 | 0.51 | -0.14 | -1.10 | 0.42 | 0.40 | 0.58 | 0.52 |
| DPP4 | 3.69 | 3.90 | 3.89 | -0.21 | -1.16 | 0.02 | -0.19 | -1.14 | 0.01 | 0.02 | 1.01 | 0.76 | 0.05 | 0.02 | 0.82 |
| DPP7 | 2.68 | 2.81 | 2.65 | -0.13 | -1.09 | 0.35 | 0.03 | 1.02 | 0.83 | 0.16 | 1.11 | 0.12 | 0.50 | 0.86 | 0.19 |
| DPT | 5.33 | 4.66 | 4.55 | 0.67 | 1.59 | 0.00 | 0.77 | 1.71 | 0.00 | 0.10 | 1.07 | 0.39 | 0.00 | 0.00 | 0.49 |
| DUOX2 | 0.06 | 0.03 | -0.01 | 0.03 | 1.02 | 0.73 | 0.07 | 1.05 | 0.34 | 0.04 | 1.03 | 0.50 | 0.82 | 0.40 | 0.60 |
| EDIL3 | 2.36 | 2.22 | 2.20 | 0.14 | 1.10 | 0.14 | 0.16 | 1.11 | 0.06 | 0.02 | 1.01 | 0.81 | 0.27 | 0.08 | 0.86 |
| EFEMP1 | 4.96 | 4.41 | 4.22 | 0.55 | 1.47 | 0.00 | 0.74 | 1.67 | 0.00 | 0.19 | 1.14 | 0.04 | 0.00 | 0.00 | 0.09 |
| EGFR | 2.65 | 2.83 | 2.82 | -0.19 | -1.14 | 0.00 | -0.17 | -1.13 | 0.00 | 0.01 | 1.01 | 0.75 | 0.00 | 0.00 | 0.81 |
| EIF4EBP1 | 4.64 | 4.41 | 3.76 | 0.23 | 1.17 | 0.38 | 0.88 | 1.84 | 0.00 | 0.65 | 1.57 | 0.00 | 0.52 | 0.00 | 0.00 |
| ENG | 4.25 | 4.22 | 4.24 | 0.02 | 1.02 | 0.70 | 0.01 | 1.00 | 0.91 | -0.02 | -1.01 | 0.69 | 0.80 | 0.93 | 0.77 |
| ENPP2 | 1.73 | 1.55 | 1.58 | 0.18 | 1.13 | 0.13 | 0.15 | 1.11 | 0.12 | -0.02 | -1.02 | 0.78 | 0.25 | 0.16 | 0.83 |

| | | | | | | | | | | | | | | | |
|---|---|---|---|---|---|---|---|---|---|---|---|---|---|---|---|
| ENTPD5 | 3.05 | 3.18 | 3.02 | -0.13 | -1.09 | 0.23 | 0.03 | 1.02 | 0.77 | 0.16 | 1.11 | 0.04 | 0.38 | 0.82 | 0.09 |
| ENTPD6 | 1.89 | 1.85 | 1.63 | 0.04 | 1.03 | 0.59 | 0.26 | 1.20 | 0.00 | 0.22 | 1.17 | 0.00 | 0.71 | 0.00 | 0.00 |
| EPHB4 | 4.27 | 3.80 | 3.58 | 0.47 | 1.38 | 0.00 | 0.69 | 1.61 | 0.00 | 0.22 | 1.16 | 0.02 | 0.00 | 0.00 | 0.04 |
| EPHX2 | 0.40 | 0.60 | 0.39 | -0.20 | -1.15 | 0.04 | 0.00 | 1.00 | 0.96 | 0.21 | 1.15 | 0.00 | 0.11 | 0.96 | 0.01 |
| ESAM | 1.84 | 1.53 | 1.38 | 0.31 | 1.24 | 0.01 | 0.45 | 1.37 | 0.00 | 0.14 | 1.11 | 0.08 | 0.03 | 0.00 | 0.15 |
| F7 | 3.32 | 3.50 | 3.63 | -0.18 | -1.13 | 0.07 | -0.31 | -1.24 | 0.00 | -0.13 | -1.10 | 0.06 | 0.16 | 0.00 | 0.12 |
| F9 | 3.63 | 3.71 | 3.60 | -0.08 | -1.06 | 0.37 | 0.03 | 1.02 | 0.75 | 0.11 | 1.08 | 0.11 | 0.52 | 0.80 | 0.18 |
| FABP2 | 3.56 | 3.14 | 2.68 | 0.41 | 1.33 | 0.10 | 0.88 | 1.84 | 0.00 | 0.47 | 1.38 | 0.01 | 0.22 | 0.00 | 0.03 |
| FABP4 | 4.30 | 2.97 | 2.71 | 1.33 | 2.52 | 0.00 | 1.59 | 3.01 | 0.00 | 0.26 | 1.20 | 0.16 | 0.00 | 0.00 | 0.25 |
| FABP6 | 1.57 | 1.08 | 0.96 | 0.49 | 1.40 | 0.00 | 0.61 | 1.52 | 0.00 | 0.12 | 1.09 | 0.24 | 0.01 | 0.00 | 0.34 |
| FADD | 1.90 | 1.82 | 1.53 | 0.08 | 1.06 | 0.63 | 0.37 | 1.29 | 0.01 | 0.29 | 1.23 | 0.01 | 0.75 | 0.01 | 0.03 |
| FAM3C | 6.04 | 5.24 | 4.97 | 0.80 | 1.75 | 0.00 | 1.07 | 2.11 | 0.00 | 0.27 | 1.21 | 0.08 | 0.00 | 0.00 | 0.14 |
| FAP | 0.33 | 0.36 | 0.41 | -0.03 | -1.02 | 0.69 | -0.07 | -1.05 | 0.23 | -0.05 | -1.03 | 0.38 | 0.80 | 0.29 | 0.48 |
| FAS | 6.31 | 5.96 | 5.80 | 0.35 | 1.27 | 0.00 | 0.51 | 1.42 | 0.00 | 0.16 | 1.12 | 0.05 | 0.01 | 0.00 | 0.11 |
| FBP1 | 2.61 | 3.03 | 2.04 | -0.42 | -1.33 | 0.09 | 0.57 | 1.49 | 0.01 | 0.99 | 1.98 | 0.00 | 0.20 | 0.01 | 0.00 |
| FCGR2A | 4.63 | 4.67 | 4.42 | -0.05 | -1.03 | 0.73 | 0.20 | 1.15 | 0.08 | 0.25 | 1.19 | 0.01 | 0.82 | 0.12 | 0.03 |
| FCGR3B | 1.89 | 1.97 | 1.86 | -0.09 | -1.06 | 0.46 | 0.03 | 1.02 | 0.78 | 0.11 | 1.08 | 0.17 | 0.60 | 0.82 | 0.26 |
| FCN2 | 4.91 | 5.23 | 5.31 | -0.33 | -1.26 | 0.01 | -0.40 | -1.32 | 0.00 | -0.08 | -1.05 | 0.43 | 0.05 | 0.00 | 0.52 |
| FCRL1 | 3.33 | 3.38 | 3.46 | -0.05 | -1.03 | 0.69 | -0.13 | -1.10 | 0.21 | -0.09 | -1.06 | 0.34 | 0.80 | 0.26 | 0.44 |
| FETUB | 3.47 | 4.06 | 4.44 | -0.58 | -1.50 | 0.00 | -0.97 | -1.96 | 0.00 | -0.39 | -1.31 | 0.00 | 0.00 | 0.00 | 0.00 |
| FGFR1OP | 1.24 | 1.24 | 0.94 | -0.01 | -1.00 | 0.97 | 0.29 | 1.23 | 0.01 | 0.30 | 1.23 | 0.00 | 0.99 | 0.01 | 0.00 |
| FUCA1 | 4.20 | 4.39 | 4.26 | -0.19 | -1.14 | 0.23 | -0.06 | -1.05 | 0.65 | 0.13 | 1.09 | 0.26 | 0.38 | 0.71 | 0.36 |
| GAS6 | 7.38 | 7.24 | 7.05 | 0.14 | 1.10 | 0.14 | 0.33 | 1.26 | 0.00 | 0.19 | 1.14 | 0.01 | 0.27 | 0.00 | 0.02 |
| GDF15 | 4.66 | 4.14 | 3.33 | 0.52 | 1.43 | 0.03 | 1.33 | 2.51 | 0.00 | 0.81 | 1.75 | 0.00 | 0.09 | 0.00 | 0.00 |
| GDF2 | 2.79 | 2.87 | 2.91 | -0.08 | -1.06 | 0.39 | -0.12 | -1.09 | 0.15 | -0.04 | -1.03 | 0.58 | 0.54 | 0.19 | 0.67 |
| GGH | 2.74 | 3.18 | 3.20 | -0.44 | -1.36 | 0.00 | -0.45 | -1.37 | 0.00 | -0.01 | -1.01 | 0.88 | 0.00 | 0.00 | 0.91 |
| GH1 | 4.30 | 3.90 | 4.28 | 0.40 | 1.32 | 0.26 | 0.02 | 1.01 | 0.95 | -0.39 | -1.31 | 0.13 | 0.41 | 0.96 | 0.21 |
| GHRL | 3.56 | 2.57 | 2.63 | 0.99 | 1.99 | 0.00 | 0.93 | 1.91 | 0.00 | -0.06 | -1.04 | 0.76 | 0.00 | 0.00 | 0.82 |
| GLO1 | 6.52 | 6.54 | 6.04 | -0.01 | -1.01 | 0.96 | 0.48 | 1.40 | 0.01 | 0.50 | 1.41 | 0.00 | 0.99 | 0.02 | 0.01 |
| GLRX | 4.04 | 3.75 | 3.31 | 0.29 | 1.22 | 0.16 | 0.73 | 1.65 | 0.00 | 0.44 | 1.35 | 0.00 | 0.30 | 0.00 | 0.01 |
| GP1BA | 4.53 | 4.32 | 4.36 | 0.21 | 1.15 | 0.04 | 0.17 | 1.12 | 0.05 | -0.04 | -1.03 | 0.58 | 0.10 | 0.07 | 0.67 |
| GP2 | 4.27 | 3.34 | 2.69 | 0.93 | 1.90 | 0.00 | 1.58 | 2.98 | 0.00 | 0.65 | 1.57 | 0.00 | 0.00 | 0.00 | 0.00 |
| GPNMB | 3.43 | 3.30 | 3.26 | 0.13 | 1.09 | 0.20 | 0.17 | 1.12 | 0.05 | 0.04 | 1.03 | 0.57 | 0.35 | 0.08 | 0.67 |
| GPR37 | 4.77 | 3.87 | 3.18 | 0.90 | 1.87 | 0.00 | 1.58 | 2.99 | 0.00 | 0.68 | 1.60 | 0.00 | 0.00 | 0.00 | 0.00 |

| | | | | | | | | | | | | | | | |
|---|---|---|---|---|---|---|---|---|---|---|---|---|---|---|---|
| GRAP2 | 2.61 | 2.26 | 2.50 | 0.36 | 1.28 | 0.24 | 0.12 | 1.08 | 0.66 | -0.24 | -1.18 | 0.27 | 0.39 | 0.72 | 0.36 |
| GRK5 | 0.32 | 0.41 | 0.43 | -0.09 | -1.06 | 0.30 | -0.10 | -1.08 | 0.14 | -0.02 | -1.01 | 0.74 | 0.45 | 0.18 | 0.81 |
| GSTA1 | 4.59 | 5.31 | 4.74 | -0.72 | -1.64 | 0.02 | -0.15 | -1.11 | 0.58 | 0.57 | 1.48 | 0.01 | 0.06 | 0.64 | 0.03 |
| GUSB | 2.46 | 2.64 | 2.32 | -0.18 | -1.13 | 0.23 | 0.13 | 1.10 | 0.30 | 0.31 | 1.24 | 0.00 | 0.38 | 0.36 | 0.01 |
| GYS1 | 2.44 | 2.49 | 2.30 | -0.04 | -1.03 | 0.82 | 0.14 | 1.10 | 0.37 | 0.19 | 1.14 | 0.17 | 0.89 | 0.44 | 0.25 |
| GZMH | 2.67 | 2.72 | 2.48 | -0.05 | -1.04 | 0.76 | 0.19 | 1.14 | 0.19 | 0.24 | 1.18 | 0.05 | 0.84 | 0.24 | 0.09 |
| HEBP1 | -0.21 | -0.15 | -0.35 | -0.06 | -1.04 | 0.53 | 0.14 | 1.10 | 0.09 | 0.20 | 1.15 | 0.00 | 0.67 | 0.13 | 0.01 |
| HK2 | 0.49 | 0.52 | 0.48 | -0.03 | -1.02 | 0.55 | 0.01 | 1.01 | 0.83 | 0.04 | 1.03 | 0.27 | 0.67 | 0.86 | 0.36 |
| HMOX1 | 6.35 | 6.17 | 5.78 | 0.17 | 1.13 | 0.35 | 0.56 | 1.48 | 0.00 | 0.39 | 1.31 | 0.00 | 0.49 | 0.00 | 0.01 |
| HNRNPK | 0.87 | 0.89 | 0.69 | -0.02 | -1.02 | 0.82 | 0.18 | 1.13 | 0.04 | 0.20 | 1.15 | 0.01 | 0.89 | 0.06 | 0.02 |
| HSPB1 | 6.08 | 5.72 | 5.62 | 0.36 | 1.28 | 0.18 | 0.46 | 1.37 | 0.05 | 0.10 | 1.07 | 0.61 | 0.32 | 0.07 | 0.69 |
| HSPG2 | 4.34 | 3.55 | 3.37 | 0.78 | 1.72 | 0.00 | 0.96 | 1.95 | 0.00 | 0.18 | 1.13 | 0.17 | 0.00 | 0.00 | 0.26 |
| HYAL1 | 1.43 | 1.41 | 1.40 | 0.03 | 1.02 | 0.68 | 0.03 | 1.02 | 0.56 | 0.01 | 1.00 | 0.90 | 0.80 | 0.63 | 0.91 |
| HYOU1 | 3.50 | 3.23 | 3.14 | 0.27 | 1.21 | 0.01 | 0.36 | 1.28 | 0.00 | 0.09 | 1.06 | 0.21 | 0.03 | 0.00 | 0.30 |
| ICAM1 | 6.28 | 6.31 | 6.16 | -0.04 | -1.03 | 0.69 | 0.12 | 1.09 | 0.15 | 0.16 | 1.12 | 0.02 | 0.80 | 0.19 | 0.05 |
| ICAM2 | 3.00 | 2.91 | 2.73 | 0.08 | 1.06 | 0.33 | 0.27 | 1.20 | 0.00 | 0.18 | 1.14 | 0.00 | 0.48 | 0.00 | 0.01 |
| ICAM3 | 4.00 | 3.98 | 3.92 | 0.02 | 1.02 | 0.75 | 0.09 | 1.06 | 0.20 | 0.06 | 1.04 | 0.27 | 0.83 | 0.25 | 0.36 |
| ICAM5 | 2.03 | 1.69 | 1.58 | 0.33 | 1.26 | 0.00 | 0.45 | 1.36 | 0.00 | 0.11 | 1.08 | 0.17 | 0.02 | 0.00 | 0.25 |
| IGFBP1 | 7.61 | 6.26 | 4.98 | 1.35 | 2.55 | 0.00 | 2.63 | 6.21 | 0.00 | 1.28 | 2.43 | 0.00 | 0.00 | 0.00 | 0.00 |
| IGFBP2 | 7.67 | 6.86 | 6.38 | 0.81 | 1.75 | 0.00 | 1.29 | 2.44 | 0.00 | 0.48 | 1.40 | 0.00 | 0.00 | 0.00 | 0.01 |
| IGFBP3 | 3.77 | 3.82 | 4.44 | -0.05 | -1.03 | 0.78 | -0.66 | -1.59 | 0.00 | -0.62 | -1.53 | 0.00 | 0.86 | 0.00 | 0.00 |
| IGFBP6 | 5.61 | 4.81 | 4.71 | 0.80 | 1.74 | 0.00 | 0.90 | 1.87 | 0.00 | 0.11 | 1.08 | 0.41 | 0.00 | 0.00 | 0.50 |
| IGFBP7 | 2.55 | 2.06 | 1.90 | 0.48 | 1.40 | 0.00 | 0.64 | 1.56 | 0.00 | 0.16 | 1.12 | 0.07 | 0.00 | 0.00 | 0.14 |
| IGFBPL1 | 1.89 | 1.39 | 1.16 | 0.49 | 1.41 | 0.00 | 0.73 | 1.66 | 0.00 | 0.23 | 1.17 | 0.01 | 0.00 | 0.00 | 0.03 |
| IGSF8 | 2.39 | 2.01 | 1.79 | 0.37 | 1.30 | 0.01 | 0.60 | 1.51 | 0.00 | 0.22 | 1.17 | 0.04 | 0.04 | 0.00 | 0.08 |
| IL18BP | 2.39 | 2.18 | 1.84 | 0.21 | 1.16 | 0.08 | 0.56 | 1.47 | 0.00 | 0.35 | 1.27 | 0.00 | 0.18 | 0.00 | 0.00 |
| IL19 | 3.24 | 1.92 | 1.92 | 1.33 | 2.51 | 0.00 | 1.32 | 2.50 | 0.00 | -0.01 | -1.00 | 0.97 | 0.00 | 0.00 | 0.98 |
| IL1RL1 | 3.68 | 3.38 | 2.34 | 0.30 | 1.23 | 0.13 | 1.34 | 2.54 | 0.00 | 1.04 | 2.06 | 0.00 | 0.26 | 0.00 | 0.00 |
| IL6ST | 2.90 | 2.79 | 2.63 | 0.10 | 1.07 | 0.23 | 0.26 | 1.20 | 0.00 | 0.16 | 1.12 | 0.01 | 0.38 | 0.00 | 0.03 |

**Supplementary Table 4. Statistical comparisons between deceased, intubated and non-severe groups in COVID-19 negative patients for all cardiometabolic proteins.**

| marker | Mean_Estimate_Deceased | Mean_Estimate_Intubated | Mean_Estimate_NonSevere | lgFCH_Deceased_vs_Intubated | FCH_Deceased_vs_Intubated | p_Deceased_vs_Intubated | lgFCH_Deceased_vs_NonSevere | FCH_Deceased_vs_NonSevere | p_Deceased_vs_NonSevere | lgFCH_Intubated_vs_NonSevere | FCH_Intubated_vs_NonSevere | p_Intubated_vs_NonSevere | FDR_Deceased_vs_Intubated | FDR_Deceased_vs_NonSevere | FDR_Intubated_vs_NonSevere |
|---|---|---|---|---|---|---|---|---|---|---|---|---|---|---|---|
| RNASE3 | 4.11 | 2.36 | 1.88 | 1.75 | 3.36 | 0.01 | 2.23 | 4.68 | 0.00 | 0.48 | 1.39 | 0.27 | 0.58 | 0.09 | 0.59 |
| ICAM3 | 4.49 | 3.69 | 3.67 | 0.81 | 1.75 | 0.00 | 0.82 | 1.77 | 0.00 | 0.02 | 1.01 | 0.91 | 0.34 | 0.09 | 0.98 |
| SIGLEC7 | 3.19 | 2.66 | 2.45 | 0.53 | 1.44 | 0.04 | 0.74 | 1.67 | 0.00 | 0.21 | 1.16 | 0.19 | 0.62 | 0.15 | 0.53 |
| TSPAN1 | 1.16 | 0.19 | 0.29 | 0.97 | 1.96 | 0.00 | 0.87 | 1.83 | 0.00 | -0.10 | -1.07 | 0.61 | 0.34 | 0.15 | 0.80 |
| TINAGL1 | 3.94 | 3.58 | 3.35 | 0.36 | 1.29 | 0.11 | 0.59 | 1.51 | 0.00 | 0.23 | 1.17 | 0.10 | 0.62 | 0.16 | 0.44 |
| VWF | 6.16 | 5.57 | 5.22 | 0.59 | 1.50 | 0.10 | 0.95 | 1.93 | 0.00 | 0.36 | 1.28 | 0.11 | 0.62 | 0.16 | 0.44 |
| ACY1 | 4.48 | 3.75 | 3.05 | 0.72 | 1.65 | 0.18 | 1.42 | 2.68 | 0.00 | 0.70 | 1.63 | 0.04 | 0.67 | 0.16 | 0.36 |
| CTSL | 5.25 | 4.76 | 4.31 | 0.50 | 1.41 | 0.17 | 0.94 | 1.92 | 0.00 | 0.44 | 1.36 | 0.05 | 0.67 | 0.16 | 0.38 |
| GPNMB | 4.64 | 3.85 | 3.79 | 0.79 | 1.73 | 0.02 | 0.85 | 1.80 | 0.00 | 0.06 | 1.04 | 0.78 | 0.58 | 0.16 | 0.90 |
| FCN2 | 4.36 | 5.18 | 5.29 | -0.82 | -1.76 | 0.03 | -0.93 | -1.90 | 0.01 | -0.11 | -1.08 | 0.65 | 0.62 | 0.23 | 0.82 |
| IGFBP3 | 3.49 | 4.54 | 4.49 | -1.05 | -2.07 | 0.01 | -1.00 | -2.00 | 0.01 | 0.05 | 1.04 | 0.84 | 0.58 | 0.23 | 0.93 |
| AZU1 | 3.26 | 2.85 | 1.94 | 0.41 | 1.33 | 0.45 | 1.32 | 2.49 | 0.01 | 0.91 | 1.87 | 0.01 | 0.87 | 0.23 | 0.24 |
| LEPR | 0.75 | 0.53 | 0.30 | 0.22 | 1.16 | 0.26 | 0.45 | 1.36 | 0.01 | 0.23 | 1.17 | 0.06 | 0.76 | 0.24 | 0.39 |
| DCTPP1 | 3.23 | 2.59 | 2.40 | 0.64 | 1.55 | 0.08 | 0.83 | 1.78 | 0.01 | 0.19 | 1.14 | 0.39 | 0.62 | 0.24 | 0.67 |
| FETUB | 3.04 | 3.82 | 4.00 | -0.78 | -1.71 | 0.06 | -0.96 | -1.94 | 0.01 | -0.18 | -1.13 | 0.48 | 0.62 | 0.24 | 0.72 |
| RARRES2 | 2.02 | 2.61 | 2.70 | -0.59 | -1.51 | 0.05 | -0.68 | -1.60 | 0.01 | -0.09 | -1.06 | 0.63 | 0.62 | 0.25 | 0.81 |
| TNNI3 | 1.99 | 2.30 | 0.69 | -0.31 | -1.24 | 0.59 | 1.30 | 2.46 | 0.01 | 1.61 | 3.04 | 0.00 | 0.88 | 0.25 | 0.00 |
| LILRB5 | 7.01 | 6.18 | 6.04 | 0.83 | 1.78 | 0.07 | 0.97 | 1.96 | 0.02 | 0.14 | 1.10 | 0.62 | 0.62 | 0.26 | 0.81 |
| PDGFRA | 2.55 | 1.85 | 1.79 | 0.70 | 1.62 | 0.04 | 0.75 | 1.69 | 0.01 | 0.06 | 1.04 | 0.80 | 0.62 | 0.26 | 0.91 |
| PDGFRB | 3.34 | 2.94 | 2.83 | 0.39 | 1.31 | 0.11 | 0.51 | 1.42 | 0.02 | 0.12 | 1.08 | 0.45 | 0.62 | 0.26 | 0.71 |
| PLAT | 4.80 | 4.09 | 3.85 | 0.72 | 1.65 | 0.11 | 0.95 | 1.93 | 0.02 | 0.23 | 1.18 | 0.40 | 0.62 | 0.26 | 0.68 |

| | | | | | | | | | | | | | | |
|---|---|---|---|---|---|---|---|---|---|---|---|---|---|---|
| PLXNB2 | 5.51 | 5.24 | 5.13 | 0.27 | 1.21 | 0.15 | 0.38 | 1.31 | 0.02 | 0.11 | 1.08 | 0.34 | 0.63 | 0.26 | 0.64 |
| ADA2 | 5.84 | 5.16 | 5.17 | 0.68 | 1.61 | 0.03 | 0.67 | 1.60 | 0.02 | -0.01 | -1.01 | 0.96 | 0.62 | 0.26 | 0.98 |
| BLMH | 2.80 | 2.29 | 2.18 | 0.50 | 1.42 | 0.08 | 0.62 | 1.53 | 0.02 | 0.11 | 1.08 | 0.52 | 0.62 | 0.26 | 0.76 |
| CD163 | 4.28 | 3.66 | 3.39 | 0.62 | 1.54 | 0.15 | 0.89 | 1.85 | 0.02 | 0.27 | 1.21 | 0.32 | 0.63 | 0.26 | 0.63 |
| CD97 | 4.69 | 4.36 | 4.06 | 0.33 | 1.26 | 0.26 | 0.63 | 1.55 | 0.02 | 0.30 | 1.23 | 0.12 | 0.76 | 0.26 | 0.44 |
| CNST | 0.90 | 2.38 | 2.45 | -1.48 | -2.79 | 0.05 | -1.56 | -2.94 | 0.02 | -0.07 | -1.05 | 0.87 | 0.62 | 0.26 | 0.95 |
| GAS6 | 7.60 | 6.96 | 6.99 | 0.64 | 1.56 | 0.03 | 0.62 | 1.53 | 0.02 | -0.03 | -1.02 | 0.88 | 0.62 | 0.26 | 0.95 |
| GP2 | 4.64 | 3.51 | 3.07 | 1.13 | 2.19 | 0.12 | 1.57 | 2.97 | 0.02 | 0.44 | 1.36 | 0.34 | 0.62 | 0.26 | 0.64 |
| IL1RL1 | 3.89 | 3.26 | 2.67 | 0.63 | 1.55 | 0.28 | 1.22 | 2.33 | 0.02 | 0.59 | 1.50 | 0.11 | 0.76 | 0.26 | 0.44 |
| SNAP23 | 2.53 | 4.05 | 4.08 | -1.52 | -2.87 | 0.05 | -1.54 | -2.92 | 0.02 | -0.02 | -1.02 | 0.96 | 0.62 | 0.27 | 0.98 |
| PTN | 1.49 | 0.35 | 0.09 | 1.14 | 2.20 | 0.11 | 1.40 | 2.64 | 0.03 | 0.26 | 1.20 | 0.56 | 0.62 | 0.30 | 0.78 |
| USP8 | 1.05 | 2.14 | 1.91 | -1.09 | -2.12 | 0.02 | -0.86 | -1.81 | 0.03 | 0.23 | 1.17 | 0.40 | 0.58 | 0.30 | 0.68 |
| CDH2 | 2.31 | 2.17 | 1.81 | 0.14 | 1.10 | 0.58 | 0.50 | 1.42 | 0.03 | 0.36 | 1.28 | 0.03 | 0.88 | 0.30 | 0.36 |
| CHIT1 | 3.60 | 5.46 | 5.43 | -1.86 | -3.63 | 0.05 | -1.84 | -3.57 | 0.03 | 0.03 | 1.02 | 0.97 | 0.62 | 0.30 | 0.98 |
| GGH | 2.12 | 2.50 | 2.68 | -0.38 | -1.30 | 0.19 | -0.57 | -1.48 | 0.03 | -0.18 | -1.14 | 0.31 | 0.67 | 0.30 | 0.63 |
| NOTCH1 | 6.02 | 5.75 | 5.69 | 0.27 | 1.20 | 0.12 | 0.33 | 1.26 | 0.03 | 0.06 | 1.04 | 0.57 | 0.62 | 0.33 | 0.78 |
| SEMA7A | 4.68 | 4.48 | 4.24 | 0.20 | 1.15 | 0.40 | 0.44 | 1.36 | 0.04 | 0.24 | 1.18 | 0.11 | 0.85 | 0.36 | 0.44 |
| SDC1 | 4.32 | 3.46 | 3.08 | 0.86 | 1.81 | 0.20 | 1.23 | 2.35 | 0.04 | 0.37 | 1.30 | 0.37 | 0.69 | 0.36 | 0.66 |
| MARCO | 4.94 | 4.42 | 4.32 | 0.52 | 1.43 | 0.13 | 0.62 | 1.53 | 0.04 | 0.10 | 1.07 | 0.65 | 0.62 | 0.36 | 0.82 |
| FAP | 1.05 | 0.77 | 0.66 | 0.28 | 1.21 | 0.19 | 0.39 | 1.31 | 0.04 | 0.11 | 1.08 | 0.40 | 0.67 | 0.36 | 0.68 |
| CA13 | 0.86 | 2.19 | 1.84 | -1.33 | -2.52 | 0.02 | -0.98 | -1.97 | 0.04 | 0.36 | 1.28 | 0.30 | 0.58 | 0.38 | 0.61 |
| ITGB1BP2 | 1.35 | 2.60 | 2.47 | -1.25 | -2.38 | 0.05 | -1.12 | -2.17 | 0.05 | 0.13 | 1.10 | 0.73 | 0.62 | 0.39 | 0.88 |
| PCDH17 | 1.90 | 1.52 | 1.40 | 0.37 | 1.30 | 0.19 | 0.50 | 1.41 | 0.05 | 0.12 | 1.09 | 0.48 | 0.67 | 0.39 | 0.72 |
| TCN2 | 7.32 | 7.24 | 6.94 | 0.08 | 1.05 | 0.73 | 0.38 | 1.30 | 0.05 | 0.30 | 1.23 | 0.03 | 0.90 | 0.42 | 0.36 |
| ALCAM | 2.40 | 2.00 | 1.99 | 0.41 | 1.33 | 0.09 | 0.42 | 1.33 | 0.05 | 0.01 | 1.01 | 0.95 | 0.62 | 0.42 | 0.98 |
| DPP4 | 4.16 | 3.79 | 3.78 | 0.37 | 1.30 | 0.10 | 0.38 | 1.30 | 0.06 | 0.01 | 1.00 | 0.96 | 0.62 | 0.46 | 0.98 |
| FBP1 | 1.82 | 1.50 | 1.05 | 0.31 | 1.24 | 0.49 | 0.77 | 1.70 | 0.06 | 0.45 | 1.37 | 0.12 | 0.88 | 0.46 | 0.44 |
| CEACAM8 | 3.83 | 3.39 | 2.92 | 0.44 | 1.36 | 0.42 | 0.90 | 1.87 | 0.06 | 0.47 | 1.38 | 0.17 | 0.85 | 0.46 | 0.51 |
| RCOR1 | 0.62 | 0.54 | 0.41 | 0.08 | 1.06 | 0.52 | 0.21 | 1.16 | 0.06 | 0.13 | 1.09 | 0.11 | 0.88 | 0.46 | 0.44 |
| FABP4 | 4.14 | 4.58 | 3.05 | -0.44 | -1.35 | 0.52 | 1.09 | 2.13 | 0.07 | 1.53 | 2.89 | 0.00 | 0.88 | 0.49 | 0.03 |
| IL6ST | 3.22 | 2.92 | 2.83 | 0.30 | 1.23 | 0.22 | 0.39 | 1.31 | 0.07 | 0.09 | 1.06 | 0.55 | 0.73 | 0.50 | 0.77 |
| VCAM1 | 4.14 | 3.56 | 3.61 | 0.58 | 1.49 | 0.09 | 0.53 | 1.44 | 0.08 | -0.05 | -1.03 | 0.82 | 0.62 | 0.51 | 0.93 |
| WASF1 | 0.28 | 0.87 | 0.75 | -0.59 | -1.50 | 0.06 | -0.47 | -1.38 | 0.10 | 0.12 | 1.09 | 0.54 | 0.62 | 0.51 | 0.77 |
| LILRB1 | 4.17 | 3.61 | 3.76 | 0.56 | 1.48 | 0.04 | 0.41 | 1.33 | 0.09 | -0.15 | -1.11 | 0.37 | 0.62 | 0.51 | 0.66 |

| | | | | | | | | | | | | | | | |
|---|---|---|---|---|---|---|---|---|---|---|---|---|---|---|---|
| MB | 5.73 | 6.89 | 4.71 | -1.16 | -2.23 | 0.08 | 1.03 | 2.04 | 0.08 | 2.19 | 4.55 | 0.00 | 0.62 | 0.51 | 0.00 |
| PCOLCE | 4.53 | 5.26 | 5.16 | -0.73 | -1.65 | 0.07 | -0.63 | -1.55 | 0.08 | 0.09 | 1.07 | 0.71 | 0.62 | 0.51 | 0.87 |
| PEAR1 | 1.49 | 1.34 | 1.22 | 0.15 | 1.11 | 0.41 | 0.27 | 1.21 | 0.09 | 0.13 | 1.09 | 0.26 | 0.85 | 0.51 | 0.59 |
| PRTN3 | 3.18 | 3.15 | 2.54 | 0.04 | 1.02 | 0.94 | 0.65 | 1.56 | 0.10 | 0.61 | 1.53 | 0.03 | 0.97 | 0.51 | 0.36 |
| ACAN | 2.90 | 2.55 | 2.48 | 0.36 | 1.28 | 0.20 | 0.43 | 1.34 | 0.09 | 0.07 | 1.05 | 0.69 | 0.70 | 0.51 | 0.84 |
| ANGPTL3 | 4.04 | 5.01 | 4.53 | -0.97 | -1.96 | 0.00 | -0.49 | -1.41 | 0.10 | 0.48 | 1.39 | 0.02 | 0.50 | 0.51 | 0.36 |
| CXCL5 | 3.51 | 4.48 | 4.48 | -0.97 | -1.97 | 0.13 | -0.97 | -1.96 | 0.09 | 0.00 | 1.00 | 0.99 | 0.63 | 0.51 | 1.00 |
| CXCL8 | 3.95 | 3.26 | 2.86 | 0.69 | 1.62 | 0.34 | 1.09 | 2.13 | 0.09 | 0.40 | 1.32 | 0.38 | 0.82 | 0.51 | 0.67 |
| DEFA1 | 1.58 | 1.25 | 0.83 | 0.34 | 1.26 | 0.49 | 0.76 | 1.69 | 0.08 | 0.42 | 1.34 | 0.17 | 0.88 | 0.51 | 0.51 |
| DKK3 | 4.75 | 4.16 | 4.19 | 0.59 | 1.51 | 0.11 | 0.56 | 1.47 | 0.09 | -0.03 | -1.02 | 0.90 | 0.62 | 0.51 | 0.96 |
| GRAP2 | 1.49 | 2.59 | 2.58 | -1.10 | -2.14 | 0.11 | -1.08 | -2.12 | 0.08 | 0.02 | 1.01 | 0.97 | 0.62 | 0.51 | 0.98 |
| HSPB1 | 4.45 | 5.42 | 5.38 | -0.97 | -1.96 | 0.11 | -0.93 | -1.91 | 0.08 | 0.04 | 1.02 | 0.93 | 0.62 | 0.51 | 0.98 |
| IGFBP1 | 7.19 | 7.70 | 5.89 | -0.51 | -1.42 | 0.56 | 1.31 | 2.47 | 0.09 | 1.82 | 3.52 | 0.00 | 0.88 | 0.51 | 0.05 |
| LRMP | 0.91 | 1.49 | 1.43 | -0.58 | -1.50 | 0.10 | -0.52 | -1.43 | 0.10 | 0.07 | 1.05 | 0.76 | 0.62 | 0.51 | 0.89 |
| MSTN | 0.62 | 0.95 | 1.10 | -0.33 | -1.26 | 0.32 | -0.48 | -1.40 | 0.10 | -0.15 | -1.11 | 0.46 | 0.81 | 0.51 | 0.72 |
| COL4A1 | 4.75 | 4.21 | 4.16 | 0.55 | 1.46 | 0.18 | 0.60 | 1.51 | 0.10 | 0.05 | 1.04 | 0.84 | 0.67 | 0.51 | 0.93 |
| ACP5 | 4.41 | 3.63 | 3.94 | 0.78 | 1.71 | 0.02 | 0.46 | 1.38 | 0.11 | -0.31 | -1.24 | 0.13 | 0.58 | 0.54 | 0.45 |
| ACTA2 | 1.00 | 1.09 | 0.40 | -0.10 | -1.07 | 0.82 | 0.60 | 1.52 | 0.11 | 0.70 | 1.62 | 0.01 | 0.95 | 0.54 | 0.24 |
| ICAM2 | 2.99 | 2.73 | 2.64 | 0.25 | 1.19 | 0.30 | 0.35 | 1.27 | 0.11 | 0.09 | 1.07 | 0.55 | 0.79 | 0.55 | 0.77 |
| STK4 | 0.06 | 0.44 | 0.35 | -0.37 | -1.30 | 0.07 | -0.29 | -1.22 | 0.12 | 0.09 | 1.06 | 0.48 | 0.62 | 0.55 | 0.72 |
| THOP1 | 5.15 | 4.80 | 4.67 | 0.35 | 1.27 | 0.31 | 0.47 | 1.39 | 0.12 | 0.13 | 1.09 | 0.55 | 0.81 | 0.56 | 0.77 |
| CLEC5A | 3.36 | 2.89 | 2.77 | 0.47 | 1.38 | 0.28 | 0.59 | 1.51 | 0.12 | 0.12 | 1.09 | 0.65 | 0.76 | 0.57 | 0.82 |
| ICAM1 | 6.66 | 6.14 | 6.29 | 0.52 | 1.44 | 0.05 | 0.37 | 1.29 | 0.12 | -0.16 | -1.11 | 0.35 | 0.62 | 0.57 | 0.65 |
| SORT1 | 4.46 | 4.41 | 4.14 | 0.05 | 1.04 | 0.83 | 0.32 | 1.25 | 0.14 | 0.27 | 1.20 | 0.08 | 0.95 | 0.57 | 0.44 |
| SOST | 5.08 | 4.70 | 4.64 | 0.38 | 1.30 | 0.28 | 0.44 | 1.35 | 0.16 | 0.06 | 1.04 | 0.78 | 0.76 | 0.57 | 0.90 |
| LILRA5 | 4.66 | 4.25 | 4.25 | 0.41 | 1.33 | 0.19 | 0.41 | 1.33 | 0.14 | 0.00 | 1.00 | 1.00 | 0.67 | 0.57 | 1.00 |
| MSMB | 1.35 | 1.28 | 0.78 | 0.07 | 1.05 | 0.86 | 0.57 | 1.48 | 0.13 | 0.50 | 1.41 | 0.06 | 0.96 | 0.57 | 0.40 |
| NPTXR | 2.82 | 2.59 | 2.41 | 0.24 | 1.18 | 0.47 | 0.42 | 1.33 | 0.16 | 0.18 | 1.13 | 0.39 | 0.88 | 0.57 | 0.67 |
| PROC | 2.86 | 3.47 | 3.24 | -0.60 | -1.52 | 0.04 | -0.38 | -1.30 | 0.14 | 0.23 | 1.17 | 0.21 | 0.62 | 0.57 | 0.55 |
| AOC3 | 2.12 | 2.06 | 1.79 | 0.07 | 1.05 | 0.79 | 0.33 | 1.26 | 0.13 | 0.27 | 1.20 | 0.09 | 0.93 | 0.57 | 0.44 |
| CASP3 | 2.01 | 3.01 | 2.72 | -1.01 | -2.01 | 0.07 | -0.71 | -1.64 | 0.14 | 0.29 | 1.23 | 0.39 | 0.62 | 0.57 | 0.67 |
| CD69 | 3.77 | 4.86 | 4.66 | -1.09 | -2.13 | 0.12 | -0.89 | -1.86 | 0.15 | 0.20 | 1.15 | 0.65 | 0.62 | 0.57 | 0.82 |
| CDH5 | 2.25 | 2.17 | 1.98 | 0.08 | 1.06 | 0.71 | 0.27 | 1.21 | 0.15 | 0.19 | 1.14 | 0.15 | 0.90 | 0.57 | 0.48 |
| CDH6 | 2.05 | 1.63 | 1.57 | 0.43 | 1.34 | 0.26 | 0.48 | 1.40 | 0.15 | 0.06 | 1.04 | 0.81 | 0.76 | 0.57 | 0.92 |

| | | | | | | | | | | | | | | |
|---|---|---|---|---|---|---|---|---|---|---|---|---|---|---|
| DCN | 3.03 | 2.72 | 2.56 | 0.30 | 1.24 | 0.40 | 0.47 | 1.38 | 0.14 | 0.16 | 1.12 | 0.47 | 0.85 | 0.57 | 0.72 |
| DIABLO | 0.74 | 1.35 | 1.33 | -0.61 | -1.53 | 0.19 | -0.60 | -1.51 | 0.15 | 0.02 | 1.01 | 0.95 | 0.67 | 0.57 | 0.98 |
| ENG | 4.68 | 4.55 | 4.42 | 0.13 | 1.09 | 0.51 | 0.26 | 1.20 | 0.13 | 0.13 | 1.10 | 0.28 | 0.88 | 0.57 | 0.59 |
| ENPP2 | 2.08 | 1.87 | 1.74 | 0.21 | 1.16 | 0.41 | 0.34 | 1.26 | 0.14 | 0.13 | 1.09 | 0.42 | 0.85 | 0.57 | 0.69 |
| ENTPD5 | 2.08 | 2.29 | 2.43 | -0.21 | -1.16 | 0.46 | -0.36 | -1.28 | 0.16 | -0.15 | -1.11 | 0.40 | 0.88 | 0.57 | 0.68 |
| FABP6 | 1.90 | 2.46 | 1.15 | -0.56 | -1.48 | 0.34 | 0.75 | 1.68 | 0.15 | 1.31 | 2.48 | 0.00 | 0.82 | 0.57 | 0.03 |
| FCGR3B | 2.35 | 2.12 | 1.96 | 0.24 | 1.18 | 0.43 | 0.39 | 1.31 | 0.14 | 0.16 | 1.12 | 0.39 | 0.85 | 0.57 | 0.68 |
| HEBP1 | -0.10 | -0.12 | -0.39 | 0.02 | 1.01 | 0.93 | 0.30 | 1.23 | 0.13 | 0.28 | 1.21 | 0.05 | 0.97 | 0.57 | 0.37 |
| IGFBP7 | 2.77 | 2.72 | 2.30 | 0.05 | 1.03 | 0.89 | 0.47 | 1.38 | 0.13 | 0.42 | 1.34 | 0.06 | 0.96 | 0.57 | 0.39 |
| TFPI | 4.95 | 4.96 | 4.64 | 0.00 | -1.00 | 0.99 | 0.31 | 1.24 | 0.16 | 0.32 | 1.25 | 0.05 | 0.99 | 0.57 | 0.37 |
| MEGF9 | 4.77 | 4.63 | 4.55 | 0.15 | 1.11 | 0.40 | 0.22 | 1.16 | 0.16 | 0.07 | 1.05 | 0.52 | 0.85 | 0.57 | 0.76 |
| ANG | 6.68 | 7.12 | 7.03 | -0.44 | -1.36 | 0.12 | -0.35 | -1.28 | 0.16 | 0.09 | 1.06 | 0.61 | 0.62 | 0.57 | 0.80 |
| APOM | 2.99 | 3.72 | 3.50 | -0.73 | -1.66 | 0.08 | -0.51 | -1.42 | 0.17 | 0.22 | 1.17 | 0.39 | 0.62 | 0.58 | 0.68 |
| ENTPD6 | 1.67 | 1.49 | 1.46 | 0.18 | 1.13 | 0.30 | 0.21 | 1.16 | 0.17 | 0.03 | 1.02 | 0.76 | 0.79 | 0.58 | 0.89 |
| CLC | 1.78 | 1.55 | 1.33 | 0.23 | 1.17 | 0.54 | 0.45 | 1.37 | 0.17 | 0.22 | 1.17 | 0.34 | 0.88 | 0.58 | 0.64 |
| GDF15 | 4.47 | 4.32 | 3.73 | 0.15 | 1.11 | 0.81 | 0.74 | 1.67 | 0.17 | 0.59 | 1.51 | 0.12 | 0.93 | 0.58 | 0.45 |
| MTPN | -0.01 | 0.12 | 0.09 | -0.13 | -1.09 | 0.12 | -0.10 | -1.07 | 0.18 | 0.03 | 1.02 | 0.58 | 0.62 | 0.59 | 0.79 |
| ADGRG2 | 3.66 | 3.53 | 3.29 | 0.12 | 1.09 | 0.69 | 0.37 | 1.29 | 0.18 | 0.25 | 1.19 | 0.21 | 0.90 | 0.59 | 0.54 |
| DUOX2 | 0.27 | -0.01 | 0.03 | 0.28 | 1.21 | 0.17 | 0.24 | 1.18 | 0.18 | -0.04 | -1.03 | 0.76 | 0.67 | 0.59 | 0.89 |
| PPIB | 0.18 | 1.09 | 0.61 | -0.91 | -1.88 | 0.02 | -0.43 | -1.35 | 0.19 | 0.48 | 1.39 | 0.04 | 0.58 | 0.60 | 0.36 |
| FADD | 1.07 | 1.86 | 1.48 | -0.79 | -1.73 | 0.03 | -0.41 | -1.33 | 0.18 | 0.38 | 1.30 | 0.08 | 0.62 | 0.60 | 0.44 |
| CTSD | 2.27 | 2.03 | 1.98 | 0.24 | 1.18 | 0.33 | 0.29 | 1.23 | 0.19 | 0.05 | 1.04 | 0.75 | 0.82 | 0.60 | 0.89 |
| KITLG | 2.43 | 3.15 | 2.88 | -0.72 | -1.65 | 0.07 | -0.45 | -1.37 | 0.19 | 0.27 | 1.21 | 0.27 | 0.62 | 0.60 | 0.59 |
| MEP1B | 1.07 | 0.65 | 0.62 | 0.42 | 1.33 | 0.28 | 0.44 | 1.36 | 0.19 | 0.03 | 1.02 | 0.90 | 0.76 | 0.60 | 0.97 |
| CDH17 | 0.92 | 0.74 | 0.46 | 0.18 | 1.13 | 0.66 | 0.46 | 1.37 | 0.19 | 0.28 | 1.21 | 0.26 | 0.89 | 0.60 | 0.59 |
| LDLR | 2.99 | 2.76 | 2.58 | 0.22 | 1.17 | 0.52 | 0.40 | 1.32 | 0.20 | 0.18 | 1.13 | 0.42 | 0.88 | 0.61 | 0.69 |
| DOK2 | 1.03 | 1.89 | 1.67 | -0.86 | -1.81 | 0.13 | -0.64 | -1.56 | 0.20 | 0.21 | 1.16 | 0.55 | 0.63 | 0.61 | 0.77 |
| CA3 | 6.56 | 7.46 | 5.85 | -0.90 | -1.87 | 0.15 | 0.71 | 1.63 | 0.20 | 1.61 | 3.05 | 0.00 | 0.64 | 0.62 | 0.01 |
| TYRO3 | 2.80 | 2.71 | 2.56 | 0.09 | 1.07 | 0.67 | 0.25 | 1.19 | 0.21 | 0.15 | 1.11 | 0.27 | 0.90 | 0.62 | 0.59 |
| LTBP2 | 5.24 | 4.89 | 4.71 | 0.35 | 1.27 | 0.47 | 0.53 | 1.45 | 0.21 | 0.18 | 1.14 | 0.54 | 0.88 | 0.62 | 0.77 |
| FUCA1 | 4.33 | 3.84 | 3.84 | 0.49 | 1.40 | 0.27 | 0.49 | 1.40 | 0.21 | 0.00 | -1.00 | 1.00 | 0.76 | 0.62 | 1.00 |
| PLTP | 3.59 | 3.42 | 3.24 | 0.17 | 1.12 | 0.60 | 0.35 | 1.28 | 0.21 | 0.19 | 1.14 | 0.35 | 0.88 | 0.62 | 0.65 |
| CBLIF | 2.27 | 2.16 | 1.68 | 0.11 | 1.08 | 0.84 | 0.58 | 1.50 | 0.21 | 0.47 | 1.39 | 0.15 | 0.95 | 0.62 | 0.49 |
| FCGR2A | 5.24 | 4.80 | 4.84 | 0.44 | 1.35 | 0.24 | 0.40 | 1.32 | 0.22 | -0.03 | -1.02 | 0.88 | 0.76 | 0.65 | 0.96 |

| | | | | | | | | | | | | | | | |
|---|---|---|---|---|---|---|---|---|---|---|---|---|---|---|---|
| DDC | 4.67 | 4.92 | 4.12 | -0.25 | -1.19 | 0.62 | 0.55 | 1.47 | 0.23 | 0.81 | 1.75 | 0.01 | 0.88 | 0.65 | 0.28 |
| OSMR | 0.89 | 0.80 | 0.73 | 0.09 | 1.07 | 0.53 | 0.16 | 1.12 | 0.23 | 0.07 | 1.05 | 0.48 | 0.88 | 0.66 | 0.72 |
| TFRC | 1.73 | 2.28 | 2.11 | -0.55 | -1.47 | 0.14 | -0.38 | -1.31 | 0.24 | 0.17 | 1.12 | 0.47 | 0.63 | 0.66 | 0.72 |
| TSHB | 2.68 | 2.44 | 2.08 | 0.25 | 1.19 | 0.67 | 0.60 | 1.52 | 0.24 | 0.35 | 1.28 | 0.33 | 0.90 | 0.66 | 0.64 |
| NRP1 | 2.40 | 2.24 | 2.16 | 0.16 | 1.12 | 0.48 | 0.24 | 1.18 | 0.24 | 0.08 | 1.06 | 0.58 | 0.88 | 0.66 | 0.79 |
| ANPEP | 1.56 | 1.46 | 1.30 | 0.09 | 1.07 | 0.71 | 0.26 | 1.19 | 0.24 | 0.16 | 1.12 | 0.28 | 0.90 | 0.66 | 0.60 |
| BPIFB1 | 5.54 | 6.22 | 6.08 | -0.67 | -1.59 | 0.19 | -0.53 | -1.45 | 0.24 | 0.14 | 1.10 | 0.66 | 0.67 | 0.66 | 0.82 |
| THBD | 3.69 | 3.54 | 3.30 | 0.15 | 1.11 | 0.70 | 0.39 | 1.31 | 0.25 | 0.24 | 1.18 | 0.31 | 0.90 | 0.67 | 0.63 |
| CXCL16 | 6.57 | 6.62 | 6.26 | -0.05 | -1.04 | 0.85 | 0.30 | 1.23 | 0.25 | 0.36 | 1.28 | 0.06 | 0.95 | 0.67 | 0.39 |
| SERPINE1 | 6.21 | 6.50 | 5.62 | -0.30 | -1.23 | 0.62 | 0.59 | 1.50 | 0.26 | 0.89 | 1.85 | 0.02 | 0.88 | 0.69 | 0.33 |
| PCSK9 | 2.14 | 2.28 | 2.39 | -0.14 | -1.10 | 0.58 | -0.25 | -1.19 | 0.26 | -0.11 | -1.08 | 0.48 | 0.88 | 0.69 | 0.72 |
| PTPRF | 3.77 | 3.86 | 3.53 | -0.09 | -1.06 | 0.72 | 0.24 | 1.18 | 0.27 | 0.32 | 1.25 | 0.03 | 0.90 | 0.70 | 0.36 |
| TIMP1 | 6.83 | 6.66 | 6.48 | 0.17 | 1.12 | 0.63 | 0.35 | 1.27 | 0.27 | 0.18 | 1.13 | 0.43 | 0.89 | 0.71 | 0.69 |
| TIA1 | 0.55 | 0.85 | 0.72 | -0.29 | -1.23 | 0.09 | -0.16 | -1.12 | 0.28 | 0.13 | 1.09 | 0.23 | 0.62 | 0.72 | 0.57 |
| ITGB2 | 5.23 | 5.08 | 5.00 | 0.15 | 1.11 | 0.55 | 0.24 | 1.18 | 0.28 | 0.09 | 1.06 | 0.57 | 0.88 | 0.72 | 0.78 |
| LGALS3 | 4.19 | 4.29 | 3.88 | -0.10 | -1.07 | 0.77 | 0.32 | 1.24 | 0.29 | 0.41 | 1.33 | 0.05 | 0.92 | 0.72 | 0.38 |
| PDGFA | 1.31 | 1.83 | 1.68 | -0.52 | -1.43 | 0.18 | -0.36 | -1.29 | 0.29 | 0.15 | 1.11 | 0.53 | 0.67 | 0.72 | 0.77 |
| PILRB | 3.00 | 3.01 | 2.57 | -0.01 | -1.01 | 0.98 | 0.43 | 1.34 | 0.29 | 0.44 | 1.36 | 0.13 | 0.99 | 0.72 | 0.45 |
| PLA2G2A | 1.97 | 2.57 | 2.67 | -0.60 | -1.52 | 0.42 | -0.70 | -1.62 | 0.29 | -0.10 | -1.07 | 0.83 | 0.85 | 0.72 | 0.93 |
| CGREF1 | 3.16 | 2.78 | 2.69 | 0.38 | 1.30 | 0.44 | 0.47 | 1.38 | 0.28 | 0.09 | 1.06 | 0.77 | 0.87 | 0.72 | 0.90 |
| CSTB | 3.17 | 3.28 | 2.67 | -0.11 | -1.08 | 0.83 | 0.50 | 1.41 | 0.30 | 0.61 | 1.53 | 0.07 | 0.95 | 0.73 | 0.43 |
| SFTPD | 3.48 | 4.14 | 4.01 | -0.66 | -1.58 | 0.25 | -0.53 | -1.44 | 0.31 | 0.14 | 1.10 | 0.71 | 0.76 | 0.73 | 0.87 |
| PGLYRP1 | 4.94 | 5.92 | 5.42 | -0.99 | -1.98 | 0.07 | -0.48 | -1.40 | 0.31 | 0.51 | 1.42 | 0.13 | 0.62 | 0.73 | 0.45 |
| DPT | 4.60 | 5.26 | 4.93 | -0.67 | -1.59 | 0.07 | -0.34 | -1.26 | 0.30 | 0.33 | 1.26 | 0.16 | 0.62 | 0.73 | 0.49 |
| F7 | 3.10 | 3.51 | 3.38 | -0.41 | -1.33 | 0.19 | -0.29 | -1.22 | 0.31 | 0.13 | 1.09 | 0.52 | 0.67 | 0.73 | 0.76 |
| FAS | 6.61 | 6.79 | 6.29 | -0.18 | -1.13 | 0.61 | 0.32 | 1.25 | 0.31 | 0.50 | 1.41 | 0.03 | 0.88 | 0.73 | 0.36 |
| REN | 2.53 | 2.77 | 2.00 | -0.24 | -1.18 | 0.68 | 0.53 | 1.44 | 0.32 | 0.77 | 1.70 | 0.04 | 0.90 | 0.75 | 0.36 |
| CLTA | 0.32 | 0.65 | 0.47 | -0.32 | -1.25 | 0.06 | -0.15 | -1.11 | 0.32 | 0.17 | 1.13 | 0.11 | 0.62 | 0.75 | 0.44 |
| HK2 | 0.40 | 0.35 | 0.49 | 0.05 | 1.03 | 0.65 | -0.09 | -1.07 | 0.32 | -0.14 | -1.10 | 0.04 | 0.89 | 0.75 | 0.36 |
| ICAM5 | 2.51 | 2.32 | 2.25 | 0.20 | 1.15 | 0.51 | 0.26 | 1.20 | 0.32 | 0.07 | 1.05 | 0.72 | 0.88 | 0.75 | 0.87 |
| PRSS27 | 2.73 | 3.30 | 3.07 | -0.56 | -1.48 | 0.14 | -0.33 | -1.26 | 0.32 | 0.23 | 1.17 | 0.34 | 0.63 | 0.75 | 0.64 |
| TNC | 3.67 | 3.60 | 3.29 | 0.08 | 1.05 | 0.86 | 0.39 | 1.31 | 0.33 | 0.31 | 1.24 | 0.27 | 0.96 | 0.75 | 0.59 |
| ZBTB17 | 1.35 | 1.22 | 1.11 | 0.13 | 1.09 | 0.65 | 0.24 | 1.18 | 0.34 | 0.11 | 1.08 | 0.54 | 0.89 | 0.75 | 0.77 |
| BMP6 | 2.61 | 2.44 | 2.41 | 0.17 | 1.13 | 0.47 | 0.20 | 1.15 | 0.34 | 0.03 | 1.02 | 0.85 | 0.88 | 0.75 | 0.93 |

| | | | | | | | | | | | | | | | |
|---|---|---|---|---|---|---|---|---|---|---|---|---|---|---|---|
| CNTN1 | 2.00 | 2.04 | 1.81 | -0.04 | -1.02 | 0.88 | 0.19 | 1.14 | 0.34 | 0.23 | 1.17 | 0.11 | 0.96 | 0.75 | 0.44 |
| EDIL3 | 2.94 | 2.66 | 2.65 | 0.29 | 1.22 | 0.41 | 0.29 | 1.23 | 0.33 | 0.01 | 1.01 | 0.97 | 0.85 | 0.75 | 0.98 |
| EGFR | 2.75 | 2.62 | 2.62 | 0.14 | 1.10 | 0.38 | 0.13 | 1.10 | 0.33 | 0.00 | -1.00 | 0.98 | 0.85 | 0.75 | 1.00 |
| IL19 | 3.17 | 3.11 | 2.65 | 0.06 | 1.04 | 0.92 | 0.52 | 1.44 | 0.34 | 0.46 | 1.38 | 0.23 | 0.97 | 0.75 | 0.58 |
| EIF4EBP1 | 4.44 | 4.60 | 3.91 | -0.16 | -1.12 | 0.80 | 0.53 | 1.44 | 0.34 | 0.69 | 1.61 | 0.09 | 0.93 | 0.76 | 0.44 |
| TGM2 | 4.73 | 4.68 | 4.28 | 0.05 | 1.03 | 0.93 | 0.44 | 1.36 | 0.36 | 0.40 | 1.32 | 0.25 | 0.97 | 0.77 | 0.59 |
| TIE1 | 4.55 | 4.66 | 4.33 | -0.11 | -1.08 | 0.70 | 0.22 | 1.17 | 0.35 | 0.33 | 1.26 | 0.06 | 0.90 | 0.77 | 0.39 |
| AHCY | 1.79 | 1.78 | 1.49 | 0.01 | 1.01 | 0.98 | 0.30 | 1.23 | 0.36 | 0.29 | 1.22 | 0.21 | 0.99 | 0.77 | 0.54 |
| AMY2A | 0.97 | 1.82 | 1.30 | -0.85 | -1.80 | 0.04 | -0.33 | -1.26 | 0.36 | 0.52 | 1.44 | 0.04 | 0.62 | 0.77 | 0.36 |
| BOC | 2.49 | 2.40 | 2.26 | 0.10 | 1.07 | 0.74 | 0.23 | 1.18 | 0.36 | 0.14 | 1.10 | 0.44 | 0.90 | 0.77 | 0.71 |
| CLEC1A | 2.06 | 2.01 | 1.76 | 0.04 | 1.03 | 0.90 | 0.29 | 1.22 | 0.36 | 0.25 | 1.19 | 0.27 | 0.97 | 0.77 | 0.59 |
| TGFBI | 4.83 | 4.76 | 4.66 | 0.08 | 1.05 | 0.73 | 0.18 | 1.13 | 0.37 | 0.10 | 1.07 | 0.47 | 0.90 | 0.77 | 0.72 |
| AMY2B | 0.86 | 1.68 | 1.17 | -0.82 | -1.77 | 0.04 | -0.31 | -1.24 | 0.37 | 0.51 | 1.42 | 0.04 | 0.62 | 0.77 | 0.36 |
| CD55 | 3.90 | 3.76 | 3.60 | 0.14 | 1.10 | 0.72 | 0.30 | 1.23 | 0.38 | 0.16 | 1.12 | 0.50 | 0.90 | 0.78 | 0.74 |
| CORO1A | 0.92 | 1.52 | 1.26 | -0.60 | -1.51 | 0.17 | -0.34 | -1.26 | 0.38 | 0.26 | 1.20 | 0.33 | 0.67 | 0.78 | 0.64 |
| TYMP | 4.23 | 4.06 | 3.85 | 0.16 | 1.12 | 0.73 | 0.37 | 1.29 | 0.38 | 0.21 | 1.15 | 0.49 | 0.90 | 0.78 | 0.73 |
| UMOD | 3.28 | 2.67 | 2.96 | 0.61 | 1.52 | 0.15 | 0.32 | 1.25 | 0.39 | -0.29 | -1.22 | 0.27 | 0.63 | 0.78 | 0.59 |
| PLIN3 | 2.00 | 2.20 | 1.75 | -0.20 | -1.15 | 0.56 | 0.25 | 1.19 | 0.39 | 0.45 | 1.37 | 0.03 | 0.88 | 0.78 | 0.36 |
| AXL | 3.40 | 3.35 | 3.15 | 0.05 | 1.04 | 0.87 | 0.25 | 1.19 | 0.38 | 0.20 | 1.15 | 0.33 | 0.96 | 0.78 | 0.64 |
| C1QTNF1 | 3.45 | 3.24 | 3.27 | 0.21 | 1.16 | 0.38 | 0.18 | 1.13 | 0.40 | -0.03 | -1.02 | 0.85 | 0.85 | 0.79 | 0.93 |
| C2 | 4.64 | 4.67 | 4.43 | -0.03 | -1.02 | 0.93 | 0.21 | 1.15 | 0.40 | 0.23 | 1.17 | 0.18 | 0.97 | 0.79 | 0.52 |
| GRK5 | 0.58 | 0.48 | 0.44 | 0.10 | 1.07 | 0.60 | 0.14 | 1.10 | 0.40 | 0.04 | 1.03 | 0.72 | 0.88 | 0.79 | 0.87 |
| SPON2 | 4.86 | 4.85 | 4.52 | 0.01 | 1.01 | 0.98 | 0.34 | 1.27 | 0.41 | 0.33 | 1.26 | 0.26 | 0.99 | 0.79 | 0.59 |
| LGALS1 | 3.16 | 3.61 | 2.90 | -0.45 | -1.36 | 0.21 | 0.26 | 1.20 | 0.40 | 0.71 | 1.63 | 0.00 | 0.70 | 0.79 | 0.06 |
| NADK | 3.07 | 3.13 | 2.74 | -0.06 | -1.04 | 0.89 | 0.33 | 1.26 | 0.40 | 0.39 | 1.31 | 0.16 | 0.96 | 0.79 | 0.49 |
| SPP1 | 8.54 | 8.43 | 8.15 | 0.11 | 1.08 | 0.83 | 0.40 | 1.32 | 0.41 | 0.28 | 1.22 | 0.41 | 0.95 | 0.80 | 0.68 |
| SEMA3F | 1.90 | 1.97 | 1.72 | -0.06 | -1.05 | 0.80 | 0.18 | 1.13 | 0.43 | 0.24 | 1.18 | 0.13 | 0.93 | 0.81 | 0.46 |
| SERPINA11 | 6.26 | 6.48 | 6.54 | -0.22 | -1.17 | 0.57 | -0.28 | -1.21 | 0.43 | -0.06 | -1.04 | 0.82 | 0.88 | 0.81 | 0.93 |
| MNDA | 2.04 | 2.05 | 1.66 | -0.01 | -1.01 | 0.99 | 0.39 | 1.31 | 0.43 | 0.40 | 1.32 | 0.25 | 0.99 | 0.81 | 0.59 |
| ADAM15 | 4.12 | 3.70 | 3.91 | 0.42 | 1.34 | 0.18 | 0.22 | 1.16 | 0.42 | -0.20 | -1.15 | 0.30 | 0.67 | 0.81 | 0.62 |
| CD2AP | 4.04 | 4.69 | 4.43 | -0.65 | -1.56 | 0.26 | -0.39 | -1.31 | 0.43 | 0.25 | 1.19 | 0.48 | 0.76 | 0.81 | 0.72 |
| CNDP1 | 0.60 | 1.00 | 0.84 | -0.40 | -1.32 | 0.24 | -0.24 | -1.18 | 0.43 | 0.16 | 1.12 | 0.44 | 0.75 | 0.81 | 0.71 |
| CTSH | 3.00 | 2.78 | 2.73 | 0.22 | 1.16 | 0.56 | 0.26 | 1.20 | 0.43 | 0.04 | 1.03 | 0.85 | 0.88 | 0.81 | 0.93 |
| ACE2 | 1.78 | 1.85 | 1.47 | -0.07 | -1.05 | 0.87 | 0.30 | 1.23 | 0.44 | 0.37 | 1.29 | 0.18 | 0.96 | 0.82 | 0.52 |

| | | | | | | | | | | | | | | |
|---|---|---|---|---|---|---|---|---|---|---|---|---|---|---|
| AK1 | 2.40 | 2.42 | 2.05 | -0.02 | -1.01 | 0.97 | 0.36 | 1.28 | 0.46 | 0.37 | 1.30 | 0.27 | 0.99 | 0.84 | 0.59 |
| FAM3C | 5.49 | 6.55 | 5.90 | -1.06 | -2.09 | 0.10 | -0.41 | -1.33 | 0.46 | 0.65 | 1.57 | 0.10 | 0.62 | 0.85 | 0.44 |
| GLRX | 3.53 | 3.75 | 3.24 | -0.22 | -1.16 | 0.63 | 0.29 | 1.22 | 0.46 | 0.51 | 1.42 | 0.07 | 0.88 | 0.85 | 0.43 |
| PRKAR1A | 2.61 | 3.21 | 2.96 | -0.59 | -1.51 | 0.27 | -0.34 | -1.27 | 0.47 | 0.25 | 1.19 | 0.45 | 0.76 | 0.85 | 0.72 |
| PDCD6 | 2.59 | 2.35 | 2.34 | 0.24 | 1.18 | 0.55 | 0.25 | 1.19 | 0.47 | 0.01 | 1.01 | 0.96 | 0.88 | 0.85 | 0.98 |
| ADH4 | 2.24 | 2.82 | 1.80 | -0.58 | -1.49 | 0.40 | 0.44 | 1.36 | 0.47 | 1.02 | 2.03 | 0.02 | 0.85 | 0.85 | 0.35 |
| GP1BA | 4.36 | 4.71 | 4.54 | -0.35 | -1.27 | 0.24 | -0.19 | -1.14 | 0.48 | 0.16 | 1.12 | 0.37 | 0.75 | 0.85 | 0.66 |
| SUSD1 | 0.93 | 1.04 | 1.00 | -0.11 | -1.08 | 0.34 | -0.07 | -1.05 | 0.48 | 0.04 | 1.03 | 0.60 | 0.82 | 0.86 | 0.79 |
| S100A11 | 1.34 | 1.86 | 1.14 | -0.52 | -1.43 | 0.12 | 0.20 | 1.15 | 0.49 | 0.72 | 1.65 | 0.00 | 0.62 | 0.86 | 0.03 |
| PLXNB3 | 0.94 | 0.93 | 0.82 | 0.01 | 1.00 | 0.97 | 0.11 | 1.08 | 0.49 | 0.10 | 1.08 | 0.36 | 0.99 | 0.86 | 0.65 |
| CD14 | 6.39 | 6.60 | 6.62 | -0.21 | -1.16 | 0.58 | -0.23 | -1.18 | 0.49 | -0.02 | -1.02 | 0.92 | 0.88 | 0.86 | 0.98 |
| SCARF1 | 3.33 | 3.82 | 3.54 | -0.49 | -1.40 | 0.16 | -0.21 | -1.15 | 0.49 | 0.28 | 1.21 | 0.20 | 0.66 | 0.86 | 0.53 |
| ANGPTL1 | 1.74 | 1.60 | 1.59 | 0.14 | 1.10 | 0.57 | 0.15 | 1.11 | 0.50 | 0.01 | 1.00 | 0.96 | 0.88 | 0.87 | 0.98 |
| CTSZ | 1.39 | 1.47 | 1.52 | -0.08 | -1.06 | 0.71 | -0.13 | -1.10 | 0.50 | -0.05 | -1.04 | 0.72 | 0.90 | 0.87 | 0.87 |
| GLO1 | 6.15 | 6.31 | 5.84 | -0.16 | -1.12 | 0.75 | 0.30 | 1.23 | 0.51 | 0.47 | 1.38 | 0.15 | 0.91 | 0.88 | 0.49 |
| GUSB | 2.39 | 2.45 | 2.18 | -0.06 | -1.04 | 0.87 | 0.21 | 1.16 | 0.51 | 0.27 | 1.21 | 0.23 | 0.96 | 0.88 | 0.58 |
| SPARCL1 | 3.89 | 4.04 | 3.72 | -0.15 | -1.11 | 0.63 | 0.17 | 1.13 | 0.52 | 0.32 | 1.25 | 0.10 | 0.89 | 0.89 | 0.44 |
| NOTCH3 | 3.01 | 2.89 | 2.84 | 0.11 | 1.08 | 0.71 | 0.17 | 1.12 | 0.53 | 0.06 | 1.04 | 0.76 | 0.90 | 0.90 | 0.89 |
| REG3A | 4.80 | 5.08 | 4.41 | -0.28 | -1.21 | 0.70 | 0.39 | 1.31 | 0.55 | 0.67 | 1.59 | 0.15 | 0.90 | 0.91 | 0.49 |
| THPO | 1.34 | 1.54 | 1.46 | -0.20 | -1.15 | 0.41 | -0.13 | -1.09 | 0.56 | 0.07 | 1.05 | 0.63 | 0.85 | 0.91 | 0.81 |
| PPP1R2 | 1.04 | 1.64 | 1.22 | -0.60 | -1.51 | 0.08 | -0.18 | -1.13 | 0.56 | 0.42 | 1.34 | 0.05 | 0.62 | 0.91 | 0.38 |
| CA1 | 7.82 | 7.81 | 7.50 | 0.01 | 1.01 | 0.98 | 0.32 | 1.25 | 0.55 | 0.31 | 1.24 | 0.42 | 0.99 | 0.91 | 0.69 |
| COMP | 6.05 | 6.24 | 5.84 | -0.20 | -1.14 | 0.62 | 0.21 | 1.15 | 0.55 | 0.40 | 1.32 | 0.10 | 0.88 | 0.91 | 0.44 |
| COMT | 2.90 | 3.87 | 3.18 | -0.96 | -1.95 | 0.07 | -0.28 | -1.21 | 0.55 | 0.69 | 1.61 | 0.04 | 0.62 | 0.91 | 0.36 |
| EPHB4 | 4.02 | 4.00 | 3.84 | 0.02 | 1.01 | 0.95 | 0.18 | 1.13 | 0.55 | 0.16 | 1.12 | 0.46 | 0.98 | 0.91 | 0.72 |
| IL18BP | 2.18 | 1.96 | 1.99 | 0.22 | 1.16 | 0.55 | 0.20 | 1.15 | 0.55 | -0.02 | -1.02 | 0.92 | 0.88 | 0.91 | 0.98 |
| VIM | 1.22 | 1.35 | 1.04 | -0.14 | -1.10 | 0.70 | 0.18 | 1.13 | 0.56 | 0.32 | 1.24 | 0.15 | 0.90 | 0.91 | 0.49 |
| CCDC80 | 4.46 | 4.81 | 4.67 | -0.35 | -1.27 | 0.39 | -0.21 | -1.15 | 0.57 | 0.14 | 1.11 | 0.57 | 0.85 | 0.92 | 0.78 |
| CPA1 | 5.36 | 5.86 | 5.06 | -0.49 | -1.41 | 0.42 | 0.31 | 1.24 | 0.57 | 0.80 | 1.74 | 0.04 | 0.85 | 0.92 | 0.36 |
| QDPR | 3.23 | 3.60 | 3.02 | -0.37 | -1.29 | 0.39 | 0.21 | 1.16 | 0.58 | 0.58 | 1.49 | 0.03 | 0.85 | 0.93 | 0.36 |
| CRTAC1 | 2.38 | 2.70 | 2.58 | -0.32 | -1.24 | 0.44 | -0.20 | -1.15 | 0.58 | 0.12 | 1.08 | 0.65 | 0.87 | 0.93 | 0.82 |
| MFAP5 | 3.48 | 4.12 | 3.69 | -0.64 | -1.56 | 0.15 | -0.21 | -1.16 | 0.58 | 0.43 | 1.35 | 0.12 | 0.63 | 0.93 | 0.44 |
| ITIH3 | 1.83 | 1.95 | 2.00 | -0.12 | -1.09 | 0.73 | -0.17 | -1.12 | 0.59 | -0.05 | -1.03 | 0.83 | 0.90 | 0.93 | 0.93 |
| LBP | 4.05 | 4.19 | 4.25 | -0.14 | -1.10 | 0.73 | -0.20 | -1.15 | 0.59 | -0.06 | -1.04 | 0.83 | 0.90 | 0.94 | 0.93 |

| | | | | | | | | | | | | | | |
|---|---|---|---|---|---|---|---|---|---|---|---|---|---|---|
| SERPINB5 | 0.28 | 0.67 | 0.39 | -0.39 | -1.31 | 0.15 | -0.11 | -1.08 | 0.65 | 0.28 | 1.21 | 0.09 | 0.63 | 0.94 | 0.44 |
| SELE | 6.00 | 5.69 | 5.82 | 0.31 | 1.24 | 0.49 | 0.19 | 1.14 | 0.63 | -0.12 | -1.09 | 0.66 | 0.88 | 0.94 | 0.82 |
| SNX9 | 2.31 | 2.70 | 2.50 | -0.39 | -1.31 | 0.39 | -0.19 | -1.14 | 0.64 | 0.20 | 1.15 | 0.48 | 0.85 | 0.94 | 0.72 |
| STK11 | 2.24 | 2.21 | 2.06 | 0.04 | 1.03 | 0.93 | 0.18 | 1.14 | 0.62 | 0.15 | 1.11 | 0.57 | 0.97 | 0.94 | 0.78 |
| THBS4 | 4.19 | 4.32 | 3.98 | -0.13 | -1.09 | 0.78 | 0.21 | 1.16 | 0.60 | 0.34 | 1.26 | 0.24 | 0.93 | 0.94 | 0.58 |
| SIRPA | 6.41 | 6.28 | 6.26 | 0.13 | 1.09 | 0.72 | 0.15 | 1.11 | 0.64 | 0.02 | 1.01 | 0.93 | 0.90 | 0.94 | 0.98 |
| TNF | 1.29 | 1.23 | 1.41 | 0.06 | 1.04 | 0.83 | -0.12 | -1.09 | 0.63 | -0.19 | -1.14 | 0.31 | 0.95 | 0.94 | 0.63 |
| VAMP5 | 1.02 | 1.12 | 1.12 | -0.11 | -1.08 | 0.66 | -0.11 | -1.08 | 0.62 | 0.00 | -1.00 | 0.99 | 0.90 | 0.94 | 1.00 |
| MCAM | 1.09 | 1.26 | 0.98 | -0.18 | -1.13 | 0.46 | 0.10 | 1.07 | 0.63 | 0.28 | 1.21 | 0.07 | 0.88 | 0.94 | 0.41 |
| NID1 | 3.87 | 4.03 | 3.75 | -0.16 | -1.12 | 0.55 | 0.12 | 1.08 | 0.62 | 0.28 | 1.21 | 0.10 | 0.88 | 0.94 | 0.44 |
| PRSS2 | 2.48 | 3.20 | 2.76 | -0.72 | -1.65 | 0.27 | -0.28 | -1.22 | 0.62 | 0.44 | 1.35 | 0.29 | 0.76 | 0.94 | 0.60 |
| ADAMTS13 | 3.99 | 3.91 | 4.06 | 0.07 | 1.05 | 0.70 | -0.08 | -1.06 | 0.64 | -0.15 | -1.11 | 0.20 | 0.90 | 0.94 | 0.54 |
| CHL1 | 2.52 | 2.51 | 2.42 | 0.01 | 1.01 | 0.96 | 0.10 | 1.07 | 0.65 | 0.09 | 1.06 | 0.57 | 0.99 | 0.94 | 0.78 |
| COL1A1 | 0.50 | 0.71 | 0.59 | -0.21 | -1.16 | 0.32 | -0.09 | -1.07 | 0.62 | 0.12 | 1.09 | 0.37 | 0.81 | 0.94 | 0.66 |
| CST6 | 3.22 | 3.87 | 3.45 | -0.65 | -1.57 | 0.23 | -0.23 | -1.17 | 0.64 | 0.42 | 1.34 | 0.21 | 0.75 | 0.94 | 0.55 |
| DLK1 | 4.60 | 4.90 | 4.82 | -0.30 | -1.23 | 0.58 | -0.22 | -1.17 | 0.64 | 0.08 | 1.06 | 0.82 | 0.88 | 0.94 | 0.93 |
| GH1 | 5.56 | 5.73 | 5.16 | -0.18 | -1.13 | 0.85 | 0.40 | 1.32 | 0.64 | 0.57 | 1.49 | 0.34 | 0.95 | 0.94 | 0.64 |
| GHRL | 3.44 | 3.36 | 3.75 | 0.08 | 1.06 | 0.91 | -0.31 | -1.24 | 0.63 | -0.39 | -1.31 | 0.40 | 0.97 | 0.94 | 0.68 |
| GYS1 | 2.17 | 2.72 | 2.36 | -0.55 | -1.47 | 0.26 | -0.20 | -1.15 | 0.64 | 0.35 | 1.28 | 0.25 | 0.76 | 0.94 | 0.59 |
| GZMH | 1.94 | 1.90 | 1.74 | 0.04 | 1.03 | 0.93 | 0.20 | 1.15 | 0.63 | 0.16 | 1.11 | 0.60 | 0.97 | 0.94 | 0.79 |
| NTproBNP | 3.71 | 4.41 | 3.25 | -0.70 | -1.63 | 0.54 | 0.46 | 1.38 | 0.65 | 1.16 | 2.24 | 0.11 | 0.88 | 0.94 | 0.44 |
| SOD1 | 1.56 | 1.80 | 1.41 | -0.24 | -1.18 | 0.53 | 0.15 | 1.11 | 0.66 | 0.39 | 1.31 | 0.11 | 0.88 | 0.95 | 0.44 |
| ROR1 | 1.48 | 1.54 | 1.36 | -0.06 | -1.04 | 0.85 | 0.12 | 1.09 | 0.68 | 0.18 | 1.13 | 0.38 | 0.95 | 0.95 | 0.67 |
| KYAT1 | 4.13 | 4.63 | 3.94 | -0.50 | -1.41 | 0.34 | 0.19 | 1.14 | 0.68 | 0.69 | 1.62 | 0.04 | 0.82 | 0.95 | 0.36 |
| MET | 4.67 | 4.99 | 4.75 | -0.31 | -1.24 | 0.11 | -0.07 | -1.05 | 0.67 | 0.24 | 1.18 | 0.05 | 0.62 | 0.95 | 0.38 |
| OLR1 | 2.89 | 3.07 | 2.76 | -0.17 | -1.13 | 0.64 | 0.14 | 1.10 | 0.67 | 0.31 | 1.24 | 0.18 | 0.89 | 0.95 | 0.52 |
| PLA2G1B | 3.86 | 5.26 | 4.08 | -1.40 | -2.64 | 0.02 | -0.21 | -1.16 | 0.68 | 1.19 | 2.28 | 0.00 | 0.58 | 0.95 | 0.06 |
| CCL16 | 6.08 | 6.53 | 5.92 | -0.45 | -1.37 | 0.29 | 0.16 | 1.12 | 0.68 | 0.61 | 1.53 | 0.02 | 0.78 | 0.95 | 0.36 |
| HYOU1 | 3.45 | 3.82 | 3.53 | -0.38 | -1.30 | 0.13 | -0.09 | -1.06 | 0.68 | 0.29 | 1.22 | 0.07 | 0.63 | 0.95 | 0.41 |
| ITGB1 | 2.56 | 2.69 | 2.48 | -0.13 | -1.09 | 0.52 | 0.07 | 1.05 | 0.69 | 0.20 | 1.15 | 0.11 | 0.88 | 0.95 | 0.44 |
| NCAM1 | 1.53 | 1.77 | 1.60 | -0.24 | -1.18 | 0.25 | -0.07 | -1.05 | 0.69 | 0.17 | 1.12 | 0.20 | 0.76 | 0.95 | 0.53 |
| TGFBR3 | 4.36 | 4.57 | 4.24 | -0.20 | -1.15 | 0.59 | 0.12 | 1.09 | 0.71 | 0.33 | 1.25 | 0.17 | 0.88 | 0.95 | 0.51 |
| APLP1 | 1.52 | 1.45 | 1.40 | 0.07 | 1.05 | 0.84 | 0.12 | 1.09 | 0.71 | 0.05 | 1.03 | 0.83 | 0.95 | 0.95 | 0.93 |
| CANT1 | 3.39 | 3.47 | 3.32 | -0.08 | -1.06 | 0.70 | 0.07 | 1.05 | 0.70 | 0.15 | 1.11 | 0.25 | 0.90 | 0.95 | 0.59 |

| | | | | | | | | | | | | | | | |
|---|---|---|---|---|---|---|---|---|---|---|---|---|---|---|---|
| CCN3 | 3.38 | 4.31 | 3.60 | -0.93 | -1.91 | 0.15 | -0.22 | -1.16 | 0.70 | 0.71 | 1.64 | 0.08 | 0.63 | 0.95 | 0.43 |
| CDHR5 | 2.49 | 2.32 | 2.38 | 0.17 | 1.13 | 0.58 | 0.11 | 1.08 | 0.70 | -0.06 | -1.05 | 0.74 | 0.88 | 0.95 | 0.88 |
| CHI3L1 | 4.15 | 3.94 | 3.92 | 0.22 | 1.16 | 0.76 | 0.24 | 1.18 | 0.71 | 0.02 | 1.02 | 0.96 | 0.92 | 0.95 | 0.98 |
| CHRDL2 | 2.62 | 2.76 | 2.77 | -0.15 | -1.11 | 0.75 | -0.15 | -1.11 | 0.71 | 0.00 | -1.00 | 0.99 | 0.91 | 0.95 | 1.00 |
| COL6A3 | 6.46 | 6.91 | 6.24 | -0.45 | -1.36 | 0.50 | 0.22 | 1.16 | 0.71 | 0.67 | 1.59 | 0.11 | 0.88 | 0.95 | 0.44 |
| HMOX1 | 5.49 | 5.87 | 5.30 | -0.38 | -1.30 | 0.50 | 0.20 | 1.15 | 0.69 | 0.57 | 1.49 | 0.10 | 0.88 | 0.95 | 0.44 |
| MPHOSPH8 | 0.50 | 0.67 | 0.41 | -0.17 | -1.13 | 0.50 | 0.08 | 1.06 | 0.72 | 0.26 | 1.19 | 0.12 | 0.88 | 0.96 | 0.44 |
| EPHX2 | 0.29 | 0.43 | 0.22 | -0.14 | -1.10 | 0.57 | 0.07 | 1.05 | 0.72 | 0.21 | 1.16 | 0.16 | 0.88 | 0.96 | 0.49 |
| GPR37 | 3.64 | 3.65 | 3.43 | -0.01 | -1.01 | 0.99 | 0.21 | 1.15 | 0.72 | 0.22 | 1.16 | 0.60 | 0.99 | 0.96 | 0.79 |
| GSTA1 | 4.05 | 5.94 | 3.76 | -1.89 | -3.70 | 0.05 | 0.29 | 1.23 | 0.73 | 2.18 | 4.54 | 0.00 | 0.62 | 0.96 | 0.03 |
| ADAMTS16 | 0.47 | 0.44 | 0.53 | 0.04 | 1.02 | 0.85 | -0.06 | -1.04 | 0.73 | -0.09 | -1.07 | 0.44 | 0.95 | 0.97 | 0.71 |
| SSC4D | 2.32 | 2.02 | 2.15 | 0.30 | 1.23 | 0.63 | 0.17 | 1.13 | 0.76 | -0.13 | -1.09 | 0.74 | 0.89 | 0.97 | 0.88 |
| LEP | 1.90 | 2.41 | 2.11 | -0.51 | -1.43 | 0.52 | -0.21 | -1.16 | 0.77 | 0.30 | 1.23 | 0.55 | 0.88 | 0.97 | 0.77 |
| MCFD2 | 2.63 | 2.83 | 2.52 | -0.20 | -1.15 | 0.62 | 0.11 | 1.08 | 0.76 | 0.31 | 1.24 | 0.23 | 0.88 | 0.97 | 0.57 |
| PLPBP | 2.27 | 3.00 | 2.40 | -0.73 | -1.66 | 0.12 | -0.13 | -1.09 | 0.75 | 0.60 | 1.51 | 0.04 | 0.62 | 0.97 | 0.36 |
| PRCP | 2.74 | 2.71 | 2.82 | 0.04 | 1.03 | 0.88 | -0.07 | -1.05 | 0.75 | -0.11 | -1.08 | 0.50 | 0.96 | 0.97 | 0.74 |
| CELA3A | 2.72 | 3.06 | 2.57 | -0.33 | -1.26 | 0.54 | 0.15 | 1.11 | 0.75 | 0.49 | 1.40 | 0.16 | 0.88 | 0.97 | 0.49 |
| CNPY2 | 0.75 | 0.75 | 0.69 | 0.00 | -1.00 | 1.00 | 0.06 | 1.04 | 0.76 | 0.06 | 1.04 | 0.66 | 1.00 | 0.97 | 0.82 |
| CTSB | 3.34 | 3.65 | 3.25 | -0.31 | -1.24 | 0.40 | 0.10 | 1.07 | 0.76 | 0.40 | 1.32 | 0.08 | 0.85 | 0.97 | 0.43 |
| GDF2 | 3.05 | 3.22 | 3.14 | -0.18 | -1.13 | 0.61 | -0.10 | -1.07 | 0.75 | 0.08 | 1.06 | 0.71 | 0.88 | 0.97 | 0.87 |
| NTRK2 | 1.23 | 1.39 | 1.17 | -0.16 | -1.11 | 0.48 | 0.06 | 1.04 | 0.77 | 0.21 | 1.16 | 0.13 | 0.88 | 0.98 | 0.45 |
| PAG1 | 3.15 | 3.63 | 3.25 | -0.49 | -1.40 | 0.24 | -0.10 | -1.08 | 0.77 | 0.38 | 1.30 | 0.14 | 0.75 | 0.98 | 0.47 |
| QPCT | 3.49 | 4.02 | 3.53 | -0.53 | -1.44 | 0.09 | -0.04 | -1.03 | 0.89 | 0.49 | 1.41 | 0.01 | 0.62 | 0.98 | 0.25 |
| RNASET2 | 4.40 | 4.50 | 4.35 | -0.11 | -1.08 | 0.68 | 0.05 | 1.04 | 0.82 | 0.16 | 1.11 | 0.34 | 0.90 | 0.98 | 0.64 |
| REG1A | 5.77 | 6.48 | 5.91 | -0.70 | -1.63 | 0.34 | -0.14 | -1.10 | 0.83 | 0.56 | 1.48 | 0.22 | 0.82 | 0.98 | 0.57 |
| REG1B | 4.08 | 4.63 | 3.98 | -0.54 | -1.46 | 0.49 | 0.10 | 1.07 | 0.88 | 0.65 | 1.56 | 0.20 | 0.88 | 0.98 | 0.53 |
| RETN | 6.61 | 6.84 | 6.49 | -0.23 | -1.17 | 0.69 | 0.12 | 1.09 | 0.82 | 0.35 | 1.28 | 0.34 | 0.90 | 0.98 | 0.64 |
| SDC4 | 2.54 | 2.86 | 2.60 | -0.32 | -1.25 | 0.41 | -0.06 | -1.04 | 0.86 | 0.26 | 1.20 | 0.29 | 0.85 | 0.98 | 0.60 |
| SLITRK6 | 2.34 | 2.73 | 2.39 | -0.38 | -1.30 | 0.22 | -0.04 | -1.03 | 0.88 | 0.34 | 1.27 | 0.08 | 0.72 | 0.98 | 0.43 |
| SSC5D | 2.98 | 3.12 | 3.03 | -0.14 | -1.11 | 0.58 | -0.05 | -1.04 | 0.83 | 0.10 | 1.07 | 0.57 | 0.88 | 0.98 | 0.78 |
| ST6GAL1 | 5.11 | 5.48 | 5.07 | -0.37 | -1.29 | 0.18 | 0.04 | 1.03 | 0.86 | 0.41 | 1.33 | 0.02 | 0.67 | 0.98 | 0.33 |
| TFF3 | 2.38 | 2.71 | 2.30 | -0.33 | -1.26 | 0.58 | 0.08 | 1.06 | 0.88 | 0.41 | 1.33 | 0.27 | 0.88 | 0.98 | 0.59 |
| TIMD4 | 3.45 | 3.33 | 3.36 | 0.12 | 1.09 | 0.80 | 0.09 | 1.06 | 0.83 | -0.03 | -1.02 | 0.92 | 0.93 | 0.98 | 0.98 |
| PTGDS | 4.64 | 5.07 | 4.71 | -0.43 | -1.35 | 0.45 | -0.07 | -1.05 | 0.89 | 0.36 | 1.28 | 0.31 | 0.87 | 0.98 | 0.63 |

| | | | | | | | | | | | | | | | |
|---|---|---|---|---|---|---|---|---|---|---|---|---|---|---|---|
| PTPRS | 2.86 | 3.10 | 2.81 | -0.24 | -1.18 | 0.35 | 0.05 | 1.04 | 0.82 | 0.29 | 1.22 | 0.07 | 0.83 | 0.98 | 0.43 |
| TNFRSF10C | 3.60 | 3.56 | 3.55 | 0.03 | 1.02 | 0.93 | 0.05 | 1.03 | 0.89 | 0.01 | 1.01 | 0.96 | 0.97 | 0.98 | 0.98 |
| TNFSF13B | 4.63 | 4.31 | 4.70 | 0.32 | 1.25 | 0.44 | -0.07 | -1.05 | 0.85 | -0.39 | -1.31 | 0.14 | 0.87 | 0.98 | 0.47 |
| VASN | 2.26 | 2.43 | 2.22 | -0.18 | -1.13 | 0.43 | 0.04 | 1.03 | 0.84 | 0.22 | 1.16 | 0.13 | 0.85 | 0.98 | 0.45 |
| VSTM2L | -0.01 | 0.01 | -0.04 | -0.03 | -1.02 | 0.89 | 0.03 | 1.02 | 0.85 | 0.06 | 1.04 | 0.63 | 0.96 | 0.98 | 0.81 |
| XG | 2.39 | 2.71 | 2.33 | -0.32 | -1.25 | 0.57 | 0.07 | 1.05 | 0.89 | 0.38 | 1.30 | 0.27 | 0.88 | 0.98 | 0.59 |
| KIT | 1.77 | 1.79 | 1.73 | -0.02 | -1.01 | 0.94 | 0.04 | 1.03 | 0.87 | 0.06 | 1.04 | 0.72 | 0.98 | 0.98 | 0.87 |
| LILRB2 | 6.53 | 6.38 | 6.49 | 0.15 | 1.11 | 0.60 | 0.04 | 1.03 | 0.88 | -0.11 | -1.08 | 0.53 | 0.88 | 0.98 | 0.77 |
| MMP7 | 1.62 | 1.77 | 1.56 | -0.15 | -1.11 | 0.70 | 0.06 | 1.05 | 0.85 | 0.21 | 1.16 | 0.37 | 0.90 | 0.98 | 0.66 |
| NECTIN2 | 6.21 | 6.46 | 6.15 | -0.25 | -1.19 | 0.61 | 0.06 | 1.04 | 0.89 | 0.32 | 1.24 | 0.32 | 0.88 | 0.98 | 0.63 |
| NPDC1 | 2.93 | 3.55 | 2.82 | -0.62 | -1.53 | 0.27 | 0.11 | 1.08 | 0.82 | 0.73 | 1.66 | 0.04 | 0.76 | 0.98 | 0.36 |
| AKR1C4 | 0.47 | 1.14 | 0.43 | -0.67 | -1.59 | 0.04 | 0.04 | 1.03 | 0.88 | 0.71 | 1.64 | 0.00 | 0.62 | 0.98 | 0.03 |
| BAG6 | 4.45 | 4.28 | 4.38 | 0.17 | 1.12 | 0.70 | 0.07 | 1.05 | 0.86 | -0.10 | -1.07 | 0.72 | 0.90 | 0.98 | 0.87 |
| CA4 | 2.07 | 2.25 | 2.02 | -0.18 | -1.13 | 0.41 | 0.04 | 1.03 | 0.82 | 0.23 | 1.17 | 0.10 | 0.85 | 0.98 | 0.44 |
| CCL14 | 7.00 | 7.28 | 6.97 | -0.28 | -1.21 | 0.34 | 0.03 | 1.02 | 0.89 | 0.31 | 1.24 | 0.09 | 0.82 | 0.98 | 0.44 |
| CCL15 | 6.63 | 6.97 | 6.53 | -0.34 | -1.27 | 0.42 | 0.10 | 1.07 | 0.80 | 0.44 | 1.35 | 0.10 | 0.85 | 0.98 | 0.44 |
| CCL27 | 3.10 | 2.78 | 3.05 | 0.32 | 1.25 | 0.40 | 0.05 | 1.04 | 0.87 | -0.27 | -1.20 | 0.26 | 0.85 | 0.98 | 0.59 |
| CD59 | 4.31 | 4.56 | 4.21 | -0.25 | -1.19 | 0.63 | 0.10 | 1.07 | 0.83 | 0.35 | 1.28 | 0.28 | 0.88 | 0.98 | 0.59 |
| CDH1 | 2.70 | 2.95 | 2.64 | -0.25 | -1.19 | 0.45 | 0.06 | 1.04 | 0.85 | 0.31 | 1.24 | 0.14 | 0.87 | 0.98 | 0.47 |
| CEBPB | 0.39 | 0.45 | 0.35 | -0.06 | -1.04 | 0.74 | 0.04 | 1.03 | 0.79 | 0.10 | 1.07 | 0.36 | 0.90 | 0.98 | 0.66 |
| CES1 | 3.97 | 4.89 | 4.07 | -0.92 | -1.90 | 0.17 | -0.10 | -1.07 | 0.86 | 0.82 | 1.77 | 0.05 | 0.67 | 0.98 | 0.38 |
| CLUL1 | 2.17 | 2.44 | 2.22 | -0.27 | -1.21 | 0.30 | -0.05 | -1.03 | 0.85 | 0.23 | 1.17 | 0.17 | 0.79 | 0.98 | 0.51 |
| COL18A1 | 6.41 | 6.76 | 6.45 | -0.35 | -1.28 | 0.33 | -0.04 | -1.03 | 0.90 | 0.31 | 1.24 | 0.17 | 0.82 | 0.98 | 0.51 |
| CST3 | 7.23 | 7.71 | 7.17 | -0.48 | -1.39 | 0.33 | 0.07 | 1.05 | 0.88 | 0.54 | 1.46 | 0.08 | 0.82 | 0.98 | 0.43 |
| CTF1 | 0.11 | -0.04 | 0.15 | 0.15 | 1.11 | 0.55 | -0.04 | -1.03 | 0.84 | -0.19 | -1.14 | 0.22 | 0.88 | 0.98 | 0.56 |
| DNAJB8 | 0.26 | 0.28 | 0.30 | -0.02 | -1.02 | 0.90 | -0.04 | -1.03 | 0.81 | -0.02 | -1.01 | 0.89 | 0.97 | 0.98 | 0.96 |
| DPP7 | 2.32 | 2.44 | 2.36 | -0.12 | -1.09 | 0.65 | -0.04 | -1.03 | 0.87 | 0.08 | 1.06 | 0.62 | 0.89 | 0.98 | 0.80 |
| FABP2 | 3.66 | 5.82 | 3.57 | -2.17 | -4.49 | 0.01 | 0.09 | 1.06 | 0.90 | 2.26 | 4.78 | 0.00 | 0.58 | 0.98 | 0.00 |
| FCRL1 | 3.48 | 3.21 | 3.39 | 0.26 | 1.20 | 0.48 | 0.08 | 1.06 | 0.80 | -0.18 | -1.13 | 0.45 | 0.88 | 0.98 | 0.71 |
| HNRNPK | 0.73 | 0.90 | 0.76 | -0.18 | -1.13 | 0.40 | -0.04 | -1.03 | 0.84 | 0.14 | 1.10 | 0.29 | 0.85 | 0.98 | 0.60 |
| HSPG2 | 4.30 | 4.90 | 4.40 | -0.60 | -1.52 | 0.27 | -0.11 | -1.08 | 0.83 | 0.50 | 1.41 | 0.15 | 0.76 | 0.98 | 0.49 |
| IGFBP6 | 5.09 | 5.49 | 5.14 | -0.41 | -1.33 | 0.41 | -0.06 | -1.04 | 0.90 | 0.35 | 1.27 | 0.26 | 0.85 | 0.98 | 0.59 |
| F9 | 3.46 | 3.71 | 3.49 | -0.24 | -1.18 | 0.35 | -0.03 | -1.02 | 0.91 | 0.22 | 1.16 | 0.19 | 0.83 | 0.99 | 0.53 |
| CRX | -0.06 | -0.07 | -0.04 | 0.00 | 1.00 | 0.99 | -0.02 | -1.02 | 0.91 | -0.02 | -1.02 | 0.86 | 1.00 | 0.99 | 0.94 |

| | | | | | | | | | | | | | | |
|---|---|---|---|---|---|---|---|---|---|---|---|---|---|---|
| VSIR | 1.21 | 1.47 | 1.23 | -0.26 | -1.20 | 0.26 | -0.02 | -1.01 | 0.93 | 0.24 | 1.18 | 0.10 | 0.76 | 0.99 | 0.44 |
| MFAP3 | 0.19 | 0.01 | 0.18 | 0.18 | 1.13 | 0.23 | 0.01 | 1.01 | 0.92 | -0.16 | -1.12 | 0.08 | 0.75 | 0.99 | 0.43 |
| PON2 | -0.15 | -0.19 | -0.14 | 0.04 | 1.03 | 0.80 | -0.01 | -1.01 | 0.92 | -0.05 | -1.03 | 0.59 | 0.93 | 0.99 | 0.79 |
| ANXA4 | 0.43 | 0.48 | 0.44 | -0.06 | -1.04 | 0.71 | -0.01 | -1.01 | 0.92 | 0.04 | 1.03 | 0.65 | 0.90 | 0.99 | 0.82 |
| CA5A | 2.12 | 3.07 | 2.05 | -0.96 | -1.94 | 0.21 | 0.07 | 1.05 | 0.92 | 1.02 | 2.03 | 0.03 | 0.70 | 0.99 | 0.36 |
| IGFBP2 | 6.93 | 7.48 | 6.88 | -0.56 | -1.47 | 0.36 | 0.05 | 1.03 | 0.93 | 0.60 | 1.52 | 0.11 | 0.85 | 0.99 | 0.44 |
| TCL1B | 0.12 | 0.07 | 0.13 | 0.05 | 1.03 | 0.79 | -0.01 | -1.01 | 0.94 | -0.06 | -1.04 | 0.60 | 0.93 | 0.99 | 0.79 |
| LPL | 5.13 | 5.04 | 5.11 | 0.09 | 1.06 | 0.84 | 0.02 | 1.01 | 0.96 | -0.07 | -1.05 | 0.81 | 0.95 | 0.99 | 0.92 |
| LRP11 | 2.83 | 2.99 | 2.81 | -0.15 | -1.11 | 0.72 | 0.03 | 1.02 | 0.94 | 0.18 | 1.13 | 0.50 | 0.90 | 0.99 | 0.74 |
| NPPB | 3.70 | 4.21 | 3.65 | -0.51 | -1.43 | 0.65 | 0.05 | 1.03 | 0.96 | 0.56 | 1.47 | 0.42 | 0.89 | 0.99 | 0.69 |
| ACOX1 | 0.14 | 0.27 | 0.13 | -0.12 | -1.09 | 0.51 | 0.01 | 1.01 | 0.95 | 0.13 | 1.10 | 0.26 | 0.88 | 0.99 | 0.59 |
| AGXT | 1.16 | 2.00 | 1.19 | -0.84 | -1.79 | 0.17 | -0.03 | -1.02 | 0.96 | 0.81 | 1.76 | 0.04 | 0.67 | 0.99 | 0.36 |
| ART3 | 2.06 | 2.02 | 2.09 | 0.05 | 1.03 | 0.91 | -0.03 | -1.02 | 0.94 | -0.07 | -1.05 | 0.78 | 0.97 | 0.99 | 0.90 |
| CD93 | 4.61 | 4.92 | 4.62 | -0.31 | -1.24 | 0.38 | -0.02 | -1.01 | 0.96 | 0.30 | 1.23 | 0.19 | 0.85 | 0.99 | 0.53 |
| CR2 | 3.49 | 3.69 | 3.53 | -0.20 | -1.15 | 0.68 | -0.04 | -1.02 | 0.93 | 0.16 | 1.12 | 0.59 | 0.90 | 0.99 | 0.79 |
| ESAM | 1.82 | 2.10 | 1.80 | -0.28 | -1.21 | 0.46 | 0.03 | 1.02 | 0.93 | 0.30 | 1.23 | 0.20 | 0.88 | 0.99 | 0.53 |
| IGFBPL1 | 1.77 | 1.88 | 1.75 | -0.11 | -1.08 | 0.79 | 0.02 | 1.02 | 0.95 | 0.13 | 1.10 | 0.61 | 0.93 | 0.99 | 0.80 |
| IGSF8 | 2.22 | 2.46 | 2.20 | -0.24 | -1.18 | 0.59 | 0.02 | 1.01 | 0.96 | 0.27 | 1.20 | 0.35 | 0.88 | 0.99 | 0.65 |
| SELP | 3.10 | 3.20 | 3.09 | -0.10 | -1.07 | 0.78 | 0.00 | 1.00 | 0.99 | 0.11 | 1.08 | 0.65 | 0.93 | 1.00 | 0.82 |
| LACTB2 | 0.88 | 1.35 | 0.87 | -0.47 | -1.38 | 0.11 | 0.01 | 1.00 | 0.98 | 0.47 | 1.39 | 0.01 | 0.62 | 1.00 | 0.24 |
| NRCAM | 4.39 | 4.33 | 4.38 | 0.05 | 1.04 | 0.89 | 0.00 | 1.00 | 1.00 | -0.05 | -1.04 | 0.83 | 0.96 | 1.00 | 0.93 |
| PAM | 4.26 | 4.52 | 4.25 | -0.26 | -1.20 | 0.28 | 0.00 | 1.00 | 0.98 | 0.27 | 1.20 | 0.08 | 0.76 | 1.00 | 0.44 |
| CD209 | 4.04 | 4.28 | 4.05 | -0.23 | -1.18 | 0.38 | 0.00 | -1.00 | 0.99 | 0.23 | 1.17 | 0.17 | 0.85 | 1.00 | 0.51 |
| CD46 | 4.32 | 4.52 | 4.32 | -0.20 | -1.15 | 0.56 | 0.01 | 1.01 | 0.98 | 0.21 | 1.15 | 0.33 | 0.88 | 1.00 | 0.64 |
| CPB1 | 1.57 | 2.29 | 1.57 | -0.72 | -1.65 | 0.10 | 0.00 | 1.00 | 0.99 | 0.72 | 1.65 | 0.01 | 0.62 | 1.00 | 0.24 |
| EFEMP1 | 4.88 | 5.17 | 4.87 | -0.29 | -1.22 | 0.48 | 0.00 | 1.00 | 0.99 | 0.29 | 1.23 | 0.25 | 0.88 | 1.00 | 0.59 |
| FGFR1OP | 0.91 | 1.13 | 0.92 | -0.21 | -1.16 | 0.50 | 0.00 | -1.00 | 1.00 | 0.21 | 1.16 | 0.29 | 0.88 | 1.00 | 0.60 |
| HYAL1 | 1.53 | 1.62 | 1.53 | -0.09 | -1.07 | 0.61 | 0.00 | -1.00 | 1.00 | 0.09 | 1.06 | 0.42 | 0.88 | 1.00 | 0.69 |

**Supplementary Table 5. Correlation between measured and predicted cardiometabolic protein expression by Pearson correlation.**

| marker | r | p | FDR |
|---|---|---|---|
| HSPG2 | 0.951538 | 1.06E-156 | 3.76E-154 |
| CST3 | 0.940146 | 3.11E-143 | 1.10E-140 |
| FAM3C | 0.937213 | 3.40E-140 | 1.20E-137 |
| COL6A3 | 0.934811 | 8.19E-138 | 2.88E-135 |
| EPHB4 | 0.932494 | 1.34E-135 | 4.70E-133 |
| CD59 | 0.930178 | 1.82E-133 | 6.37E-131 |
| IGFBP6 | 0.926192 | 5.86E-130 | 2.05E-127 |
| CCN3 | 0.92501 | 5.89E-129 | 2.05E-126 |
| ESAM | 0.92477 | 9.36E-129 | 3.25E-126 |
| NECTIN2 | 0.922647 | 5.31E-127 | 1.84E-124 |
| NPDC1 | 0.915493 | 1.93E-121 | 6.66E-119 |
| IL18BP | 0.912541 | 2.74E-119 | 9.43E-117 |
| THBD | 0.911659 | 1.16E-118 | 3.98E-116 |
| CEACAM8 | 0.911659 | 1.16E-118 | 3.98E-116 |
| PTGDS | 0.909113 | 6.95E-117 | 2.37E-114 |
| CD46 | 0.904294 | 1.16E-113 | 3.94E-111 |
| PGLYRP1 | 0.902511 | 1.65E-112 | 5.59E-110 |
| TIMP1 | 0.898024 | 1.03E-109 | 3.48E-107 |
| SPON2 | 0.896359 | 1.05E-108 | 3.54E-106 |
| FAS | 0.894216 | 1.95E-107 | 6.55E-105 |
| IGSF8 | 0.892887 | 1.15E-106 | 3.85E-104 |
| CLEC5A | 0.891963 | 3.93E-106 | 1.31E-103 |
| CD55 | 0.891903 | 4.25E-106 | 1.42E-103 |
| DEFA1_DEFA1B | 0.891328 | 9.06E-106 | 3.01E-103 |
| REG1A | 0.888248 | 4.83E-104 | 1.60E-101 |
| XG | 0.886777 | 3.10E-103 | 1.02E-100 |
| COL18A1 | 0.884235 | 7.24E-102 | 2.38E-99 |
| RETN | 0.880481 | 6.67E-100 | 2.19E-97 |

| | | | |
|---|---|---|---|
| CD93 | 0.877109 | 3.41E-98 | 1.12E-95 |
| CD69 | 0.870534 | 5.29E-95 | 1.72E-92 |
| PDGFA | 0.869655 | 1.37E-94 | 4.45E-92 |
| TGFBR3 | 0.868278 | 6.01E-94 | 1.95E-91 |
| IGFBPL1 | 0.868045 | 7.69E-94 | 2.48E-91 |
| CLEC1A | 0.863199 | 1.21E-91 | 3.90E-89 |
| RNASET2 | 0.863154 | 1.27E-91 | 4.08E-89 |
| SCARF1 | 0.863042 | 1.42E-91 | 4.54E-89 |
| CCL14 | 0.860471 | 1.92E-90 | 6.12E-88 |
| REG3A | 0.86008 | 2.84E-90 | 9.03E-88 |
| EFEMP1 | 0.856211 | 1.28E-88 | 4.06E-86 |
| LGALS1 | 0.854727 | 5.36E-88 | 1.69E-85 |
| IL19 | 0.854514 | 6.58E-88 | 2.07E-85 |
| CXCL5 | 0.854233 | 8.61E-88 | 2.70E-85 |
| CDH1 | 0.853992 | 1.08E-87 | 3.38E-85 |
| PAG1 | 0.853823 | 1.27E-87 | 3.96E-85 |
| GPR37 | 0.851841 | 8.29E-87 | 2.58E-84 |
| CCDC80 | 0.851433 | 1.22E-86 | 3.78E-84 |
| GRAP2 | 0.850125 | 4.11E-86 | 1.27E-83 |
| ST6GAL1 | 0.846558 | 1.08E-84 | 3.33E-82 |
| SNAP23 | 0.846388 | 1.25E-84 | 3.84E-82 |
| LTBP2 | 0.845316 | 3.29E-84 | 1.01E-81 |
| NADK | 0.843643 | 1.46E-83 | 4.45E-81 |
| MCFD2 | 0.843102 | 2.35E-83 | 7.14E-81 |
| REG1B | 0.842755 | 3.19E-83 | 9.67E-81 |
| CST6 | 0.842518 | 3.93E-83 | 1.19E-80 |
| CNST | 0.841472 | 9.81E-83 | 2.95E-80 |
| LRP11 | 0.839787 | 4.23E-82 | 1.27E-79 |
| ROR1 | 0.839298 | 6.44E-82 | 1.93E-79 |
| ALCAM | 0.838389 | 1.40E-81 | 4.17E-79 |
| SEMA3F | 0.837447 | 3.12E-81 | 9.27E-79 |
| HYOU1 | 0.837371 | 3.33E-81 | 9.86E-79 |
| TFF3 | 0.836381 | 7.69E-81 | 2.27E-78 |
| LILRA5 | 0.835309 | 1.89E-80 | 5.56E-78 |

| | | | |
|---|---|---|---|
| CGREF1 | 0.834108 | 5.13E-80 | 1.50E-77 |
| VCAM1 | 0.83306 | 1.22E-79 | 3.56E-77 |
| PDGFRA | 0.830533 | 9.61E-79 | 2.80E-76 |
| SELP | 0.829692 | 1.90E-78 | 5.51E-76 |
| DLK1 | 0.829495 | 2.22E-78 | 6.42E-76 |
| USP8 | 0.829372 | 2.45E-78 | 7.06E-76 |
| PLIN3 | 0.827968 | 7.54E-78 | 2.16E-75 |
| ITGB1BP2 | 0.827294 | 1.29E-77 | 3.69E-75 |
| PTPRS | 0.827234 | 1.35E-77 | 3.85E-75 |
| PAM | 0.827145 | 1.45E-77 | 4.12E-75 |
| PPIB | 0.826233 | 2.98E-77 | 8.43E-75 |
| CA13 | 0.826207 | 3.04E-77 | 8.57E-75 |
| DOK2 | 0.82574 | 4.39E-77 | 1.23E-74 |
| CTSL | 0.825496 | 5.31E-77 | 1.49E-74 |
| SPP1 | 0.824984 | 7.93E-77 | 2.21E-74 |
| CSTB | 0.82308 | 3.48E-76 | 9.67E-74 |
| GDF15 | 0.821948 | 8.31E-76 | 2.30E-73 |
| CANT1 | 0.821508 | 1.16E-75 | 3.20E-73 |
| DCN | 0.817834 | 1.87E-74 | 5.14E-72 |
| CTSZ | 0.817195 | 3.01E-74 | 8.25E-72 |
| PRKAR1A | 0.815813 | 8.38E-74 | 2.29E-71 |
| NOTCH1 | 0.815333 | 1.19E-73 | 3.24E-71 |
| IGFBP7 | 0.814502 | 2.19E-73 | 5.93E-71 |
| DKK3 | 0.814034 | 3.09E-73 | 8.34E-71 |
| DCTPP1 | 0.813616 | 4.19E-73 | 1.13E-70 |
| ACTA2 | 0.813061 | 6.28E-73 | 1.68E-70 |
| ICAM1 | 0.812507 | 9.38E-73 | 2.50E-70 |
| PPP1R2 | 0.812466 | 9.66E-73 | 2.57E-70 |
| CTSB | 0.811914 | 1.44E-72 | 3.82E-70 |
| FABP4 | 0.809906 | 6.06E-72 | 1.60E-69 |
| IL6ST | 0.808432 | 1.72E-71 | 4.52E-69 |
| ANGPTL1 | 0.807228 | 4.02E-71 | 1.05E-68 |
| ART3 | 0.8064 | 7.18E-71 | 1.87E-68 |
| CHI3L1 | 0.805455 | 1.39E-70 | 3.61E-68 |

| | | | |
|---|---|---|---|
| IGFBP2 | 0.804065 | 3.62E-70 | 9.38E-68 |
| SPARCL1 | 0.804011 | 3.75E-70 | 9.68E-68 |
| S100A11 | 0.802824 | 8.48E-70 | 2.18E-67 |
| TINAGL1 | 0.802134 | 1.36E-69 | 3.48E-67 |
| CORO1A | 0.800319 | 4.64E-69 | 1.18E-66 |
| CASP3 | 0.799939 | 5.99E-69 | 1.52E-66 |
| CD97 | 0.798899 | 1.20E-68 | 3.04E-66 |
| CXCL8 | 0.798881 | 1.22E-68 | 3.07E-66 |
| HSPB1 | 0.796987 | 4.28E-68 | 1.07E-65 |
| LBP | 0.79673 | 5.07E-68 | 1.27E-65 |
| SORT1 | 0.789227 | 6.49E-66 | 1.62E-63 |
| GYS1 | 0.78673 | 3.12E-65 | 7.74E-63 |
| DPT | 0.786027 | 4.84E-65 | 1.20E-62 |
| CXCL16 | 0.785122 | 8.48E-65 | 2.09E-62 |
| IL1RL1 | 0.782573 | 4.07E-64 | 9.97E-62 |
| PRTN3 | 0.782141 | 5.30E-64 | 1.29E-61 |
| CD14 | 0.781474 | 7.95E-64 | 1.93E-61 |
| TNFRSF10C | 0.780636 | 1.32E-63 | 3.19E-61 |
| MFAP5 | 0.779838 | 2.14E-63 | 5.16E-61 |
| TFPI | 0.779423 | 2.75E-63 | 6.60E-61 |
| MMP7 | 0.77866 | 4.34E-63 | 1.04E-60 |
| NID1 | 0.777412 | 9.13E-63 | 2.17E-60 |
| WASF1 | 0.776362 | 1.70E-62 | 4.03E-60 |
| LRMP | 0.774524 | 5.02E-62 | 1.18E-59 |
| DIABLO | 0.774419 | 5.34E-62 | 1.25E-59 |
| TNC | 0.773498 | 9.15E-62 | 2.14E-59 |
| OLR1 | 0.772389 | 1.74E-61 | 4.05E-59 |
| PEAR1 | 0.771155 | 3.56E-61 | 8.26E-59 |
| ITGB1 | 0.770959 | 3.98E-61 | 9.19E-59 |
| ACE2 | 0.76954 | 8.98E-61 | 2.07E-58 |
| CDH2 | 0.766336 | 5.52E-60 | 1.26E-57 |
| NRP1 | 0.76579 | 7.49E-60 | 1.71E-57 |
| RNASE3 | 0.764291 | 1.73E-59 | 3.93E-57 |
| PILRB | 0.762448 | 4.80E-59 | 1.08E-56 |

| | | | |
|---|---|---|---|
| COL1A1 | 0.762075 | 5.90E-59 | 1.33E-56 |
| NOTCH3 | 0.760778 | 1.20E-58 | 2.69E-56 |
| TYMP | 0.760398 | 1.48E-58 | 3.30E-56 |
| SDC4 | 0.760372 | 1.50E-58 | 3.33E-56 |
| IGFBP1 | 0.759907 | 1.93E-58 | 4.27E-56 |
| NTproBNP | 0.757411 | 7.46E-58 | 1.64E-55 |
| SERPINE1 | 0.756192 | 1.43E-57 | 3.13E-55 |
| PLXNB2 | 0.754075 | 4.42E-57 | 9.64E-55 |
| GAS6 | 0.752106 | 1.25E-56 | 2.71E-54 |
| BOC | 0.751881 | 1.40E-56 | 3.02E-54 |
| COMT | 0.748454 | 8.32E-56 | 1.79E-53 |
| MSMB | 0.748412 | 8.51E-56 | 1.82E-53 |
| TCN2 | 0.746887 | 1.86E-55 | 3.96E-53 |
| MNDA | 0.746177 | 2.67E-55 | 5.66E-53 |
| MB | 0.745751 | 3.32E-55 | 7.01E-53 |
| PCOLCE | 0.743981 | 8.14E-55 | 1.71E-52 |
| CD163 | 0.741022 | 3.59E-54 | 7.50E-52 |
| LPL | 0.740141 | 5.56E-54 | 1.16E-51 |
| ENTPD6 | 0.738453 | 1.28E-53 | 2.65E-51 |
| PRSS2 | 0.738423 | 1.30E-53 | 2.68E-51 |
| PLA2G2A | 0.737569 | 1.97E-53 | 4.04E-51 |
| SDC1 | 0.737319 | 2.23E-53 | 4.55E-51 |
| CD209 | 0.736424 | 3.45E-53 | 7.00E-51 |
| FGFR1OP | 0.735559 | 5.26E-53 | 1.06E-50 |
| VSIR | 0.734099 | 1.07E-52 | 2.15E-50 |
| CCL15 | 0.733814 | 1.22E-52 | 2.44E-50 |
| STK4 | 0.731978 | 2.95E-52 | 5.87E-50 |
| PLAT | 0.731306 | 4.06E-52 | 8.04E-50 |
| GPNMB | 0.730951 | 4.81E-52 | 9.48E-50 |
| CLC | 0.73077 | 5.24E-52 | 1.03E-49 |
| PTPRF | 0.729756 | 8.47E-52 | 1.65E-49 |
| FADD | 0.727202 | 2.82E-51 | 5.47E-49 |
| ICAM2 | 0.727033 | 3.05E-51 | 5.89E-49 |
| PCSK9 | 0.726724 | 3.52E-51 | 6.76E-49 |

| | | | |
|---|---|---|---|
| BLMH | 0.726533 | 3.85E-51 | 7.35E-49 |
| SNX9 | 0.724494 | 9.91E-51 | 1.88E-48 |
| CLTA | 0.723819 | 1.35E-50 | 2.55E-48 |
| RARRES2 | 0.722146 | 2.91E-50 | 5.47E-48 |
| CCL27 | 0.721487 | 3.94E-50 | 7.37E-48 |
| PLXNB3 | 0.719813 | 8.42E-50 | 1.57E-47 |
| ENG | 0.719328 | 1.05E-49 | 1.94E-47 |
| FETUB | 0.718491 | 1.53E-49 | 2.82E-47 |
| LILRB2 | 0.717614 | 2.27E-49 | 4.15E-47 |
| GP1BA | 0.714996 | 7.28E-49 | 1.32E-46 |
| TIA1 | 0.711734 | 3.06E-48 | 5.54E-46 |
| PCDH17 | 0.709773 | 7.18E-48 | 1.29E-45 |
| SEMA7A | 0.709664 | 7.53E-48 | 1.35E-45 |
| PRSS27 | 0.705761 | 4.03E-47 | 7.17E-45 |
| TNF | 0.705586 | 4.34E-47 | 7.68E-45 |
| GHRL | 0.705415 | 4.67E-47 | 8.22E-45 |
| F9 | 0.703747 | 9.46E-47 | 1.66E-44 |
| SIGLEC7 | 0.703495 | 1.05E-46 | 1.83E-44 |
| VASN | 0.703465 | 1.07E-46 | 1.85E-44 |
| CHRDL2 | 0.70126 | 2.69E-46 | 4.63E-44 |
| MPHOSPH8 | 0.701112 | 2.86E-46 | 4.89E-44 |
| ICAM5 | 0.700997 | 3.00E-46 | 5.10E-44 |
| TIMD4 | 0.696326 | 2.07E-45 | 3.50E-43 |
| LGALS3 | 0.695816 | 2.55E-45 | 4.28E-43 |
| COMP | 0.694957 | 3.63E-45 | 6.06E-43 |
| C1QTNF1 | 0.693905 | 5.56E-45 | 9.23E-43 |
| CA3 | 0.689555 | 3.20E-44 | 5.28E-42 |
| ANPEP | 0.689456 | 3.33E-44 | 5.46E-42 |
| LDLR | 0.688335 | 5.19E-44 | 8.46E-42 |
| AXL | 0.687902 | 6.16E-44 | 9.98E-42 |
| SOST | 0.685964 | 1.32E-43 | 2.13E-41 |
| VWF | 0.685141 | 1.83E-43 | 2.93E-41 |
| CDH6 | 0.682286 | 5.55E-43 | 8.82E-41 |
| CCL16 | 0.680713 | 1.02E-42 | 1.61E-40 |

| | | | |
|---|---|---|---|
| FBP1 | 0.680219 | 1.23E-42 | 1.93E-40 |
| LACTB2 | 0.678185 | 2.68E-42 | 4.18E-40 |
| KITLG | 0.676611 | 4.87E-42 | 7.55E-40 |
| ZBTB17 | 0.673094 | 1.82E-41 | 2.80E-39 |
| ADAM15 | 0.671044 | 3.91E-41 | 5.98E-39 |
| TNFSF13B | 0.67074 | 4.37E-41 | 6.64E-39 |
| VAMP5 | 0.669485 | 6.95E-41 | 1.05E-38 |
| CA4 | 0.668454 | 1.01E-40 | 1.52E-38 |
| SIRPA | 0.666753 | 1.89E-40 | 2.82E-38 |
| OSMR | 0.666106 | 2.39E-40 | 3.54E-38 |
| ADAMTS16 | 0.665968 | 2.51E-40 | 3.69E-38 |
| NCAM1 | 0.664533 | 4.22E-40 | 6.16E-38 |
| TGFBI | 0.663691 | 5.72E-40 | 8.29E-38 |
| NPPB | 0.6616 | 1.21E-39 | 1.74E-37 |
| CD2AP | 0.658901 | 3.15E-39 | 4.50E-37 |
| ITGB2 | 0.654638 | 1.40E-38 | 1.99E-36 |
| BMP6 | 0.653326 | 2.21E-38 | 3.12E-36 |
| ACP5 | 0.651486 | 4.17E-38 | 5.84E-36 |
| ENTPD5 | 0.65076 | 5.35E-38 | 7.44E-36 |
| ITIH3 | 0.64857 | 1.13E-37 | 1.56E-35 |
| PLA2G1B | 0.648414 | 1.19E-37 | 1.63E-35 |
| CNTN1 | 0.647349 | 1.71E-37 | 2.33E-35 |
| FABP6 | 0.645508 | 3.17E-37 | 4.28E-35 |
| TYRO3 | 0.644426 | 4.56E-37 | 6.11E-35 |
| MEGF9 | 0.644065 | 5.15E-37 | 6.85E-35 |
| CDH5 | 0.643265 | 6.72E-37 | 8.87E-35 |
| BAG6 | 0.64023 | 1.83E-36 | 2.40E-34 |
| DPP4 | 0.63691 | 5.44E-36 | 7.07E-34 |
| AOC3 | 0.636813 | 5.61E-36 | 7.24E-34 |
| GLRX | 0.6346 | 1.15E-35 | 1.47E-33 |
| EGFR | 0.634325 | 1.26E-35 | 1.60E-33 |
| AZU1 | 0.631893 | 2.74E-35 | 3.45E-33 |
| TNNI3 | 0.631205 | 3.41E-35 | 4.26E-33 |
| AGXT | 0.631139 | 3.48E-35 | 4.32E-33 |

| | | | |
|---|---|---|---|
| AHCY | 0.628058 | 9.23E-35 | 1.14E-32 |
| C2 | 0.626251 | 1.63E-34 | 1.99E-32 |
| ADA2 | 0.623082 | 4.36E-34 | 5.28E-32 |
| HYAL1 | 0.621635 | 6.81E-34 | 8.17E-32 |
| FABP2 | 0.620504 | 9.64E-34 | 1.15E-31 |
| UMOD | 0.620431 | 9.86E-34 | 1.16E-31 |
| MCAM | 0.618693 | 1.68E-33 | 1.97E-31 |
| ENPP2 | 0.617956 | 2.10E-33 | 2.44E-31 |
| PLTP | 0.617559 | 2.36E-33 | 2.71E-31 |
| IGFBP3 | 0.615623 | 4.24E-33 | 4.83E-31 |
| CBLIF | 0.612278 | 1.15E-32 | 1.30E-30 |
| CRTAC1 | 0.610354 | 2.04E-32 | 2.28E-30 |
| HMOX1 | 0.608684 | 3.34E-32 | 3.71E-30 |
| STK11 | 0.608058 | 4.01E-32 | 4.41E-30 |
| QPCT | 0.607857 | 4.25E-32 | 4.63E-30 |
| ANG | 0.601222 | 2.90E-31 | 3.13E-29 |
| ADGRG2 | 0.601078 | 3.02E-31 | 3.23E-29 |
| APLP1 | 0.600396 | 3.67E-31 | 3.89E-29 |
| EDIL3 | 0.599102 | 5.31E-31 | 5.58E-29 |
| NTRK2 | 0.593181 | 2.80E-30 | 2.91E-28 |
| ADAMTS13 | 0.590467 | 5.94E-30 | 6.12E-28 |
| ANXA4 | 0.588202 | 1.11E-29 | 1.13E-27 |
| SUSD1 | 0.587997 | 1.17E-29 | 1.18E-27 |
| EIF4EBP1 | 0.586991 | 1.54E-29 | 1.54E-27 |
| COL4A1 | 0.586706 | 1.66E-29 | 1.64E-27 |
| GP2 | 0.584919 | 2.70E-29 | 2.65E-27 |
| FCGR2A | 0.58477 | 2.81E-29 | 2.73E-27 |
| GGH | 0.583815 | 3.63E-29 | 3.48E-27 |
| ACY1 | 0.583073 | 4.43E-29 | 4.21E-27 |
| SOD1 | 0.582205 | 5.60E-29 | 5.26E-27 |
| LEPR | 0.581516 | 6.73E-29 | 6.26E-27 |
| PTN | 0.58149 | 6.77E-29 | 6.26E-27 |
| THOP1 | 0.580695 | 8.38E-29 | 7.63E-27 |
| FCN2 | 0.575278 | 3.50E-28 | 3.15E-26 |

| | | | |
|---|---|---|---|
| CTSH | 0.574611 | 4.16E-28 | 3.70E-26 |
| ACAN | 0.574603 | 4.17E-28 | 3.70E-26 |
| KYAT1 | 0.570609 | 1.17E-27 | 1.02E-25 |
| CA1 | 0.567348 | 2.70E-27 | 2.32E-25 |
| TGM2 | 0.563973 | 6.35E-27 | 5.40E-25 |
| PROC | 0.560543 | 1.50E-26 | 1.26E-24 |
| RCOR1 | 0.556802 | 3.77E-26 | 3.13E-24 |
| PDGFRB | 0.556206 | 4.36E-26 | 3.58E-24 |
| GDF2 | 0.555331 | 5.40E-26 | 4.37E-24 |
| TIE1 | 0.551961 | 1.23E-25 | 9.84E-24 |
| CA5A | 0.551656 | 1.32E-25 | 1.04E-23 |
| GLO1 | 0.551498 | 1.37E-25 | 1.07E-23 |
| ADH4 | 0.549986 | 1.97E-25 | 1.52E-23 |
| LILRB1 | 0.548812 | 2.61E-25 | 1.98E-23 |
| VIM | 0.548142 | 3.06E-25 | 2.30E-23 |
| REN | 0.546519 | 4.51E-25 | 3.34E-23 |
| NRCAM | 0.546339 | 4.70E-25 | 3.43E-23 |
| PLPBP | 0.544832 | 6.71E-25 | 4.83E-23 |
| AK1 | 0.543663 | 8.84E-25 | 6.28E-23 |
| CEBPB | 0.543223 | 9.80E-25 | 6.86E-23 |
| MARCO | 0.542908 | 1.06E-24 | 7.31E-23 |
| CDH17 | 0.542791 | 1.08E-24 | 7.34E-23 |
| SFTPD | 0.54223 | 1.24E-24 | 8.31E-23 |
| CPB1 | 0.540777 | 1.74E-24 | 1.15E-22 |
| GSTA1 | 0.538628 | 2.86E-24 | 1.86E-22 |
| F7 | 0.538541 | 2.91E-24 | 1.86E-22 |
| CR2 | 0.536455 | 4.71E-24 | 2.97E-22 |
| CLUL1 | 0.53486 | 6.78E-24 | 4.20E-22 |
| CTSD | 0.534047 | 8.16E-24 | 4.98E-22 |
| GUSB | 0.533513 | 9.22E-24 | 5.53E-22 |
| SLITRK6 | 0.530028 | 2.02E-23 | 1.19E-21 |
| QDPR | 0.529857 | 2.10E-23 | 1.22E-21 |
| GZMH | 0.528169 | 3.07E-23 | 1.75E-21 |
| SERPINA11 | 0.526707 | 4.25E-23 | 2.38E-21 |

| | | | |
|---|---|---|---|
| SELE | 0.526477 | 4.47E-23 | 2.46E-21 |
| GH1 | 0.524074 | 7.60E-23 | 4.10E-21 |
| MET | 0.52111 | 1.45E-22 | 7.69E-21 |
| AMY2A | 0.518325 | 2.66E-22 | 1.38E-20 |
| THBS4 | 0.518273 | 2.69E-22 | 1.38E-20 |
| FUCA1 | 0.517293 | 3.32E-22 | 1.66E-20 |
| HNRNPK | 0.513438 | 7.58E-22 | 3.71E-20 |
| DPP7 | 0.511822 | 1.07E-21 | 5.14E-20 |
| ANGPTL3 | 0.511293 | 1.19E-21 | 5.59E-20 |
| FAP | 0.510488 | 1.41E-21 | 6.49E-20 |
| LILRB5 | 0.50914 | 1.88E-21 | 8.46E-20 |
| FCGR3B | 0.505523 | 3.99E-21 | 1.76E-19 |
| PDCD6 | 0.500242 | 1.18E-20 | 5.07E-19 |
| CDHR5 | 0.498553 | 1.66E-20 | 6.97E-19 |
| CES1 | 0.498204 | 1.78E-20 | 7.30E-19 |
| PRCP | 0.497579 | 2.02E-20 | 8.08E-19 |
| AMY2B | 0.495431 | 3.11E-20 | 1.21E-18 |
| GRK5 | 0.495186 | 3.27E-20 | 1.24E-18 |
| MEP1B | 0.491703 | 6.54E-20 | 2.42E-18 |
| TFRC | 0.489725 | 9.65E-20 | 3.47E-18 |
| KIT | 0.488311 | 1.27E-19 | 4.45E-18 |
| SSC5D | 0.484932 | 2.46E-19 | 8.36E-18 |
| CHL1 | 0.481156 | 5.08E-19 | 1.68E-17 |
| MSTN | 0.479938 | 6.41E-19 | 2.05E-17 |
| ICAM3 | 0.47575 | 1.41E-18 | 4.37E-17 |
| NPTXR | 0.46879 | 5.15E-18 | 1.55E-16 |
| CPA1 | 0.461373 | 1.98E-17 | 5.74E-16 |
| APOM | 0.459067 | 2.98E-17 | 8.34E-16 |
| PON2 | 0.453364 | 8.14E-17 | 2.20E-15 |
| CELA3A | 0.452339 | 9.73E-17 | 2.53E-15 |
| ACOX1 | 0.445245 | 3.29E-16 | 8.23E-15 |
| EPHX2 | 0.443521 | 4.41E-16 | 1.06E-14 |
| THPO | 0.430607 | 3.73E-15 | 8.58E-14 |
| HEBP1 | 0.424231 | 1.04E-14 | 2.29E-13 |

| | | | |
|---|---|---|---|
| DUOX2 | 0.416514 | 3.47E-14 | 7.29E-13 |
| FCRL1 | 0.414567 | 4.68E-14 | 9.36E-13 |
| HK2 | 0.396205 | 7.21E-13 | 1.37E-11 |
| BPIFB1 | 0.39368 | 1.04E-12 | 1.87E-11 |
| DDC | 0.393519 | 1.06E-12 | 1.87E-11 |
| CNDP1 | 0.378631 | 8.47E-12 | 1.36E-10 |
| SSC4D | 0.378339 | 8.81E-12 | 1.36E-10 |
| CNPY2 | 0.368764 | 3.17E-11 | 4.44E-10 |
| TCL1B | 0.368593 | 3.24E-11 | 4.44E-10 |
| MTPN | 0.351004 | 3.05E-10 | 3.66E-09 |
| CTF1 | 0.346819 | 5.10E-10 | 5.61E-09 |
| CRX | 0.328931 | 4.21E-09 | 4.21E-08 |
| AKR1C4 | 0.328641 | 4.35E-09 | 4.21E-08 |
| TSHB | 0.321531 | 9.69E-09 | 7.75E-08 |
| LEP | 0.318763 | 1.32E-08 | 9.24E-08 |
| DNAJB8 | 0.307638 | 4.37E-08 | 2.62E-07 |
| SERPINB5 | 0.290409 | 2.55E-07 | 1.28E-06 |
| TSPAN1 | 0.276532 | 9.71E-07 | 3.88E-06 |
| CHIT1 | 0.273973 | 1.23E-06 | 3.88E-06 |
| VSTM2L | 0.262391 | 3.52E-06 | 7.04E-06 |
| MFAP3 | 0.196388 | 0.000573955 | 0.000573955 |
